# Multi-omics analysis of a traditional fermented food reveals a byproduct-associated subpopulation of *Neurospora intermedia* for waste-to-food upcycling

**DOI:** 10.1101/2024.07.24.604980

**Authors:** Vayu Maini Rekdal, José Manuel Villalobos-Escobedo, Nabila Rodriguez-Valeron, Mikel Olaizola Garcia, Diego Prado Vásquez, Alexander Rosales, Pia M. Sörensen, Edward E. K. Baidoo, Ana Calheiros de Carvalho, Robert Riley, Anna Lipzen, Guifen He, Mi Yan, Sajeet Haridas, Christopher Daum, Yuko Yoshinaga, Vivian Ng, Igor V. Grigoriev, Rasmus Munk, Christofora Hanny Wijaya, Lilis Nuraida, Isty Damayanti, Pablo-Cruz Morales, Jay. D. Keasling

## Abstract

Fungal solid-state fermentation (SSF) of byproducts has promise for increasing food sustainability and security, but fungal waste-to-food upcycling remains poorly understood at the molecular level. Here we use a multi-omics approach to characterize oncom – a fermented food traditionally produced from byproducts in Java, Indonesia – as a model system for understanding fungal waste conversion. Metagenomic sequencing of two oncom types (red and black) indicated that *Neurospora intermedia* is the fungus dominating red oncom. Further transcriptomic, metabolomic, and phylogenomic analysis revealed that oncom-derived *N. intermedia* utilizes pectin and cellulose degradation for substrate conversion and belongs to a distinct byproduct-associated subpopulation that differs from wild strains at the genetic and biochemical level. Finally, we found that *N. intermedia* grew on a range of industrially relevant byproducts, did not encode for any known mycotoxins, and could be used to create foods that were positively perceived by consumers outside Indonesia. This study uncovers the microbial and genetic basis of a traditional upcycled food, sheds light on human domestication of microbes for sustainability challenges, and establishes the edible *N. intermedia* as a promising fungus for byproduct upcycling in SSF and beyond.

## Introduction

Minimizing food waste is critical for improving the resiliency and sustainability of the food system ^1–3^. In industrialized countries such as the U.S, approximately a third of food is wasted, and loss and waste of food accounts for approximately half of total greenhouse gas emissions caused by the food system on a global level^4,5^. Upgrading waste into value-added products (a process termed upcycling) can reduce the climate impact of food production by diverting waste from landfills and increasing resource use efficiency, while also enhancing food security and promoting financial benefits^6–8^. In particular, microbial processes for upcycling have shown promise for converting otherwise wasted substrates into sustainable protein^9^, chemicals^10^, materials^11^, and new foods^12^.

Filamentous fungi – a diverse group of microorganisms that include molds and mushrooms – are ideally suited for microbial upcycling of waste streams (byproducts) generated in the transformation of crops into foods. Owing to their ability to break down polymers such as cellulose, fungi can efficiently upcycle complex byproducts across diverse contexts. For instance, growing fungi on byproducts in large-scale liquid fermentations for heterologous production of animal proteins (precision fermentation) or for edible biomass (mycoprotein) is predicted to offset global greenhouse gas emissions and associated land use changes caused by resource-intensive animal agriculture^9,13^. Similarly, solid-state fermentation (SSF), a process in which fungi are directly grown on solid substrates instead of in submerged tanks, can reduce global land pressure and greenhouse gas emissions while promoting food system circularity, particularly when used to produce human food^1,2,14^. In contrast to large-scale industrial liquid fermentations, SSF is a relatively accessible technology that requires minimal equipment and inputs. Along with the local availability of many byproducts, the low technological barrier gives SSF the added potential as a decentralized solution to improve food security and nutrition for vulnerable populations^15–18^.

Despite the promises of SSF for transforming byproducts into human foods with improved nutrition and sensory appeal^18–20^, rationally engineering byproduct-strain combinations in SSF for desired outcomes remains challenging. Scientific and gastronomic investigations of fungal byproduct SSF food have been growing steadily, but most efforts to date focused on human food have been conducted on a trial-and-error basis with a poor understanding of the underlying mechanisms^12,18–25^. Characterizing fungal waste-to-food conversion in genetic and molecular detail is critical for expanding the engineering possibilities with SSF and enabling design of strain-substrate combinations for specific organoleptic properties and nutritional profiles.

Here, we use multi-omics analyses to characterize the production of oncom, a fermented food traditionally produced through SSF of readily available byproducts on the island of Java, Indonesia. Oncom is classified as either red (oncom merah) or black (oncom hitam) based on the appearance and the ingredients used. Whereas red oncom is produced from soy pulp (commonly known as okara), the main byproduct from soymilk production, black oncom is typically produced from peanut presscake, a byproduct of peanut oil production^26^ (Fig. 1A and B). Oncom production is a small cottage industry mostly confined to the Western part of Java, but these two byproducts are relevant on a global scale, with both okara and peanut oil presscake produced in massive quantities globally each year^16,27,28^. In the traditional process, the byproducts are inoculated by back-slopping from a previous batch, followed by a brief 36-48 hour fermentation period that transforms the byproducts into a firmer cake that is covered by fungal spores (Fig. 1A and 1B) ^29^. Oncom fermentation is assumed to be driven mainly by fungi, and red and black oncom fermentations have been attributed to members of the *Neurospora* and *Rhizopus* genera, respectively, based on appearance, microbial isolation efforts and experiments with pure cultures^16,29–31^. Similar to the more well-known fermented food tempeh, oncom is sold as rectangular blocks and is used as a meat substitute (Fig. 1A and B). Oncom serves diverse culinary roles in the local cuisine, including as a base for sauces, as a standalone protein, or as major ingredient mixed into rice and other dishes (Fig. 1C) ^32^.

**Figure 1.**
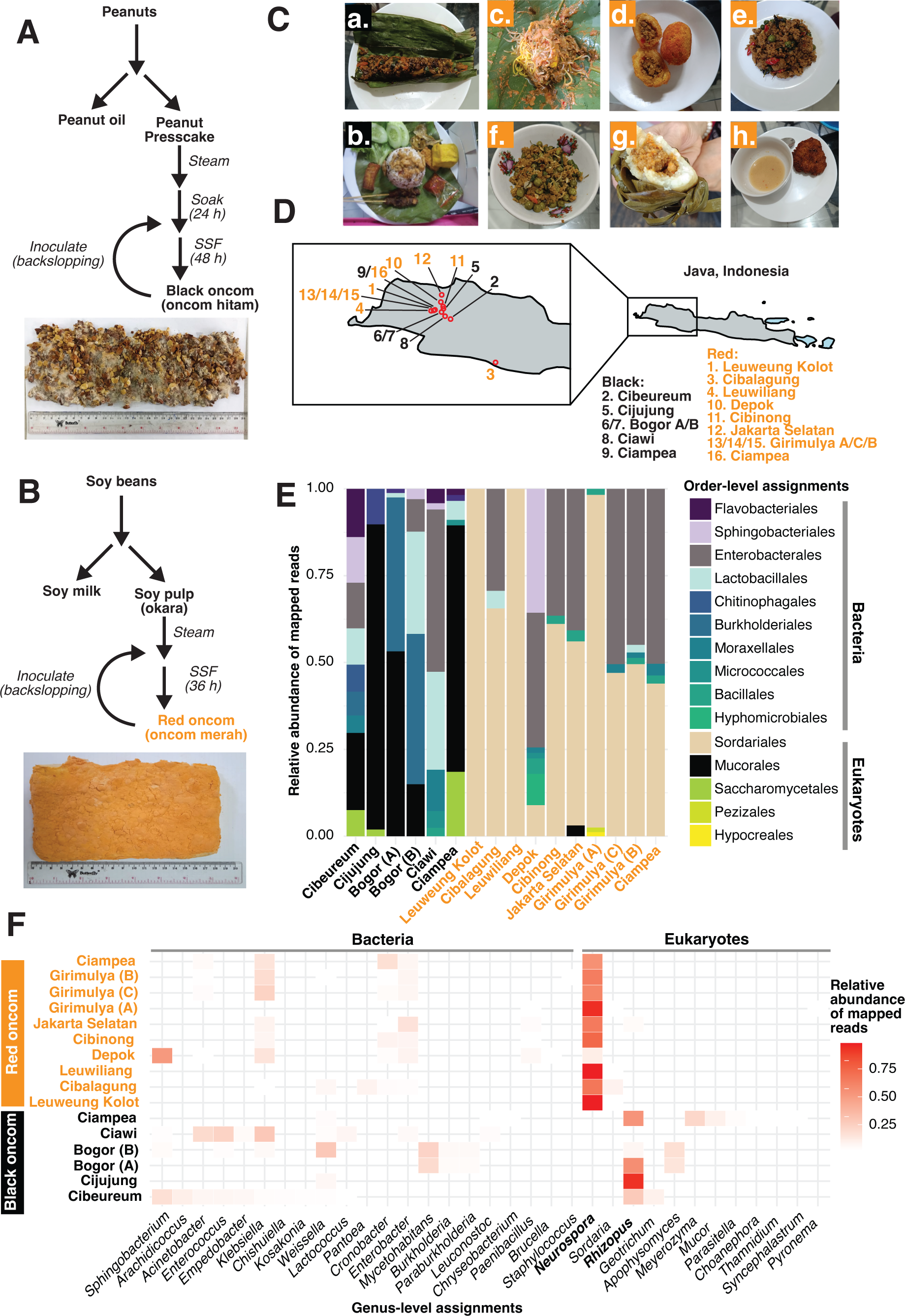
Metagenomic survey of black and red oncom collected from Java, Indonesia. A) traditional production method for black oncom. A typical sample of black oncom is shown (directly after fermentation). B) Traditional production method for red oncom. A typical sample of red oncom is shown (directly after fermentation). C) Typical preparations of oncom in the local cuisine of West Java, Indonesia. a=*pepes oncom* (black oncom cooked in a banana leaf with spices); b=*nasi tutug oncom* (black oncom cooked in rice with spices and served with side dishes and condiments); c=*toge goreng* (red oncom forms the basis of the spicy sauce used to season vegetables, other ingredients); d=*combro* (red oncom is main ingredient in stuffing); e=*krecek oncom* (red oncom stir fried with vegetables, spices); f=*oncom leunca* (red oncom cooked with leunca, a nightshade vegetable); g=*buras oncom* (red oncom is main filling of pouch steamed in banana leaf); h=*perkedel oncom* (fried cakes made from red oncom, spices, and other ingredients). D) Production sites of oncom samples analyzed by metagenome sequencing in this study. The island of Java is shown in grey, and the enlarged picture shows specific production locations. Numbers correspond to where samples were collected from. Orange = red oncom; black = black oncom. Letters indicate that more than one producer was sampled from a specific geographical location. E) Order-level taxonomy assignments using kaiju based on metagenome sequencing data. Beige shows *Sordiarales*, the fungal order to which *<u>Neurospora</u>* belongs; black shows *Mucorales*, the fungal order to which *Rhizopus* belongs. Relative abundances of non-viral mapped reads at >1% abundance threshold are shown. F) Genus-level abundance of the same samples shown in E). Relative abundances of non-viral mapped reads at >1% abundance threshold are shown. *Neurospora* and *Rhizopus*, the fungal organisms traditionally associated with red and black oncom, respectively, are highlighted in bold.

We reasoned that, as an ingenious biotechnological solution to minimizing food waste, oncom could serve as an ideal model system to close the gap in our understanding of fungal waste-to-food conversion and reveal strains, enzymes, and strategies that could inform general efforts to upcycle byproducts into human food. However, at the start of this study, the genetic and biochemical basis of waste conversion into edible oncom remained unknown. In this study, we aimed to characterize the oncom at the microbial community, genetic, and molecular levels, and investigate the broader relevance of these findings for waste-to-food conversion beyond the traditional Indonesian context. We integrated diverse omics methods (metagenomics, transcriptomics, metabolomics, phylogenomic) with biochemical, microbiological, and sensory assays to accomplish these goals, providing detailed insight into oncom as a traditional biotechnology for waste-to-food upcycling.

## Results

### Metagenomic survey of oncom reveals differences across sample type and producer

Although *Neurospora* and *Rhizopus* have been isolated from red and black oncom, respectively, and are traditionally considered the major organisms driving fermentation^16,31^, no culture-independent method have been deployed to characterize oncom samples across different producers, creating an incomplete picture of oncom microbial community composition and diversity. Specifically, we wondered whether a broader microbial consortium could be involved in substrate upcycling, as the traditional process involves minimal controls on microbial community composition beyond steaming and backslopping (Fig. 1A and 1B). To address this gap, we first collected red oncom (n=10) and black oncom (n=6) samples from artisanal producers in Western Java and used shotgun metagenome sequencing to profile microbial composition of the samples, assigning taxonomic identities to the short reads using kaiju^33,34^ (Fig. 1D, Fig. S1 and Table S1).

This metagenomic survey revealed striking differences in microbial community composition across oncom type and producer origin. Black oncom samples displayed significantly more diversity and variability in taxonomic composition between producers (Fig. 1D). For example, *Rhizopus*, assumed to be the main driver of black oncom fermentation, was detected only in five of six samples, with the sample collected from Ciawi harboring no detectable level of *Rhizopus* and instead being dominated by bacteria (Fig. 1F). Within the samples containing *Rhizopus*, the abundance of this fungus varied widely, ranging from complete dominance to comprising less than 10% of total assigned reads. It was also not always the most dominant fungus, and overall, black oncom harbored a diversity of fungi beyond *Rhizopus*, including *Geotrichum*, which is typically associated with dairy fermented foods^35^, *Mucor,* which is found in diverse Asian fermented foods^36^, *Meyerozyma,* a yeast that grows in a variety of habitats^37^, and *Apophysomyces*, a fungus associated with subtropical climates^38^ (Fig. 1E and 1F). Bacterial communities also varied between samples. Notable bacteria included *Burkholderia*^39^ and *Mycetohabitans*^40^, which are known endophytes of *Rhizpous*, as well as lactic acid bacteria such as *Weisella* and *Lactococcus,* which were present in several samples (Fig. 1E and 1F). Such bacteria may be involved in the lactic acid fermentation that takes place during the soaking of the peanut presscake prior to inoculation (Fig. 1A).

In contrast to the fungal variability observed across black oncom samples, *Neurospora* was the main fungus detected across all red oncom samples, and its abundance ranged from complete dominance (>95% of all mapped reads), to co-existing with bacteria and even low levels of other fungi (<9%) in some samples (Fig. 1E and 1F). In all red oncom samples but the one collected from Depok, in which most reads were assigned to the gram-negative bacterium *Sphingobacter*, *Neurospora* was the most abundant single microorganism, making up >50% of all classified reads (Fig. 1F). In the samples that were dominated by *Neurospora* and also harbored bacteria, the main bacterial reads were assigned to Proteobacteria such as *Klebsiella, Enterobacter,* and *Cronobacter.* These bacteria were also found in some black oncom samples (Fig. 1E and 1F). While such organisms are not typically considered major players in the production of human fermented foods, it remains unclear from our data whether these bacteria are actively participating in fermentation or are contaminants and spoilage organisms that appear post-production, which is the major context in which proteobacteria are frequently observed in human food production^41,42^. It is worth noting that the microbial community composition of red oncom differs from that of other microbially characterized soy-based foods from Asia, including miso, dajiang-meju, qu, soy sauce, sufu, and cheonggukjang. These communities do not contain *Neurospora* and instead harbor microbial communities that vary across product type and include fungi such as *Aspergillus, Rhizopus, Penicillium*, as well as diverse bacteria^43–47^.

Finally, we explored the species-level identity of the shotgun metagenomic reads assigned to the *Neurospora* genus in red oncom. Although *Neurospora intermedia* has consistently been the main fungus isolated from red oncom^16,30,31^, the database used to assign taxonomy in kaiju does not contain a *N. intermedia* genome. Thus, to enable greater species-level resolution, we first generated a genome of the oncom-derived reference strain *N. intermedia* FGSC #2613 using long-read sequencing and annotating the genes using transcriptomics. The 39 Mb genome of strain #2613 has a GC-content of 49%, consistent with members of the *Neurospora* genus and the genome quality was higher than the previous best *N. intermedia* genome^48^; it was spread across 21 contigs instead of >1000 and has a 60-fold higher N50 value reflecting its superior contiguity (Table S2). Using this genome to map the metagenomic reads, we found a strong and significant correlation between the abundance of *Neurospora* (as assessed by kaiju) and the percent of reads mapping to *N. intermedia* when this was used as the only query under a stringent (<2) mismatch threshold (Fig. S2). Furthermore, the shotgun metagenome data covered 99% of the *N. intermedia* genome (Fig. S2). These sequence results lend further evidence to previous observations^30,31^ that *N. intermedia* is the major species in red oncom.

Overall, this first metagenomic characterization of oncom samples indicates previously unappreciated microbial diversity in this traditional food. The presence of previously unreported bacteria and fungi in some oncom samples raises questions about their role in product characteristics and the fermentation process. While *Neurospora* appeared to be the major fungus involved in red oncom production, the microbial picture for black oncom fermentation may be more complex and variable than initially appreciated. This difference was further supported by 16s and ITS amplicon sequencing of a subset of black (Cijujung and Cibeureum) and red oncom samples (Leuweung Kolot, Cibalagung, Leuwiliang), which showed a dominance of a single *Sordiarales*, the fungal order to which *N. intermedia* belongs, in red oncom samples, and variable abundance of *Rhizopus* and other fungi and bacteria in the black oncom samples (Fig. S3 and S4). The finding that *Neurospora* is the only microorganisms detected in a subset of red oncom samples suggests that this organism may be sufficient for its production. This conclusion is consistent with a small-scale field trial in Java that have demonstrated that a pure *Neurospora* inoculum can produce red oncom with similar consumer acceptance as oncom produced using traditional back-slopping^16^.

We focused our further characterization efforts on red oncom because of the consistent role *Neurospora* appeared to play in the upcycling process, and the fact that red oncom is the only reported fermented food that involves intentional cultivation of the *Neurospora* genus^49^. While *Rhizopus* is used in tempeh production and other fermented foods in Asia^36^, red oncom represents a unique traditional use of *Neurospora* for human food^49^, making it an ideal model system to understand whether there are specific genetic, biochemical, and phylogenetic features that may makes *Neurospora* uniquely suitable for waste-to-food upcycling.

### *N. intermedia* changes the biochemical and sensory properties of okara and degrades cellulose and pectin during fermentation

We next set out to characterize the transformation of okara, the traditional substrate for red oncom, by *N. intermedia* at the biochemical and genetic levels. We found that SSF modestly increased the protein content (from 25% to 28%, p<0.001, unpaired t-test) and lipid content (from 12% to 14%, p<0.01, unpaired t-test) of okara but did not change the amount of crude fiber (Fig. S4). Furthermore, the overall amino acid composition did not change majorly, and fermented okara retained all essential amino acids contained in the starting material (Fig. S5). In contrast to the unremarkable changes in overall amino acid composition, *N. intermedia* fermentation released free amino acids, most notably high levels of both glutamine and glutamate (Fig. S6). While glutamate could contribute to umami flavor, glutamine has a neutral taste but could participate in Maillard reactions with reducing sugars to enhance the product upon cooking^50,51^. Consistent with this, glutamate has been identified as a contributor to umami taste in red oncom^26^.

We also detected changes in the volatile aroma composition of okara upon *N. intermedia* fermentation. One of the most striking changes was a 40-fold decrease in hexanal to barely detectable levels, an off-flavor that originates from lipid oxidation in leguminous substrates and is considered a major off-flavor in substrates such as okara^52^ (Table S3). Finally, okara SSF produced high levels of ergothioneine, a potent antioxidant that has been associated with several health benefits in humans^53,54^. Ergothioneine was absent in okara but was elevated upon fermentation to levels approximately 25% of those found in oyster mushrooms, the dietary source containing the highest known levels^53,55^ (Fig. S7). While the model fungus *Neurospora crassa* is known to produce ergothioneine^56^, these data indicate *N. intermedia* is a source of ergothioneine when used in SSF, which may further enhance the value of byproducts fermented using this fungus. Taken together, these results suggest that *N. intermedia* fermentation modulates the biochemical composition of okara in ways that potentially make it more palatable and functionally beneficial to humans.

We next sought to identify the genes and enzymes supporting the growth of *N. intermedia* on okara during the fermentation process. As the byproduct of extraction of soluble protein and sugars into soymilk, okara is largely composed of polysaccharides originating from the plant cell wall, including pectin and cellulose^57,58^. The consistent presence of *N. intermedia* in red oncom samples indicates that the fungus may be well-adapted to break down these substrates. Upon further characterization of *Neurospora-*fermented okara, we noted release of free sugars, including arabinose, galactose, and glucose, which were undetectable or found at low levels in the starting material (Fig. 2A). As these free sugars could be derived from breakdown of complex polysaccharides, these results strongly suggested activity of Carbohydrate Active Enzymes (CAZymes), a group of enzymes that are associated with degradation of complex polysaccharides and support fungal growth on plant biomass^59^. We used transcriptomics to identify potential CAZymes involved in okara degradation by *N. intermedia*. First, we grew *N. intermedia* as attached mycelial mats in submerged cultivation using sucrose as the carbon source. Following a growth period, we then washed the mycelia and switched to okara as the only carbon source for a brief period of induction. To identify genes induced by okara, we compared transcriptomes to strains induced on a no carbon control^59^.

**Figure 2.**
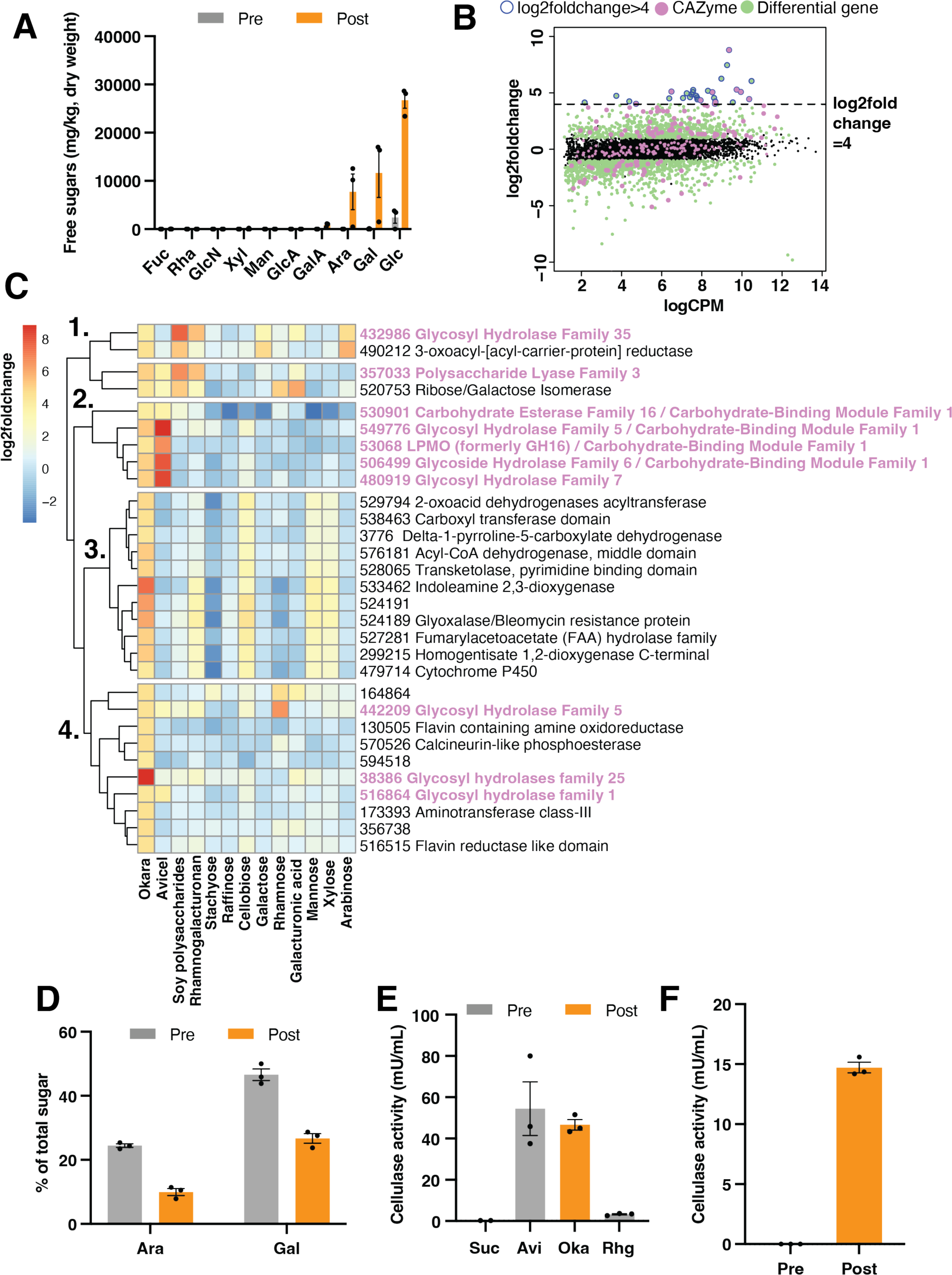
Transcriptomics identifies cellulose and pectin degradation during okara breakdown by *N. intermedia*. A) Free sugars detected in okara subjected to SSF by *N. intermedia,* before and after the fermentation. Fuc = fucose, Rha = rhamnose, GlcN = N-acetylglucosamine, Xyl = xylose, Man = mannose, GlcA = glucuronic acid, GalA = galacturonic acid, Ara = arabinose, Gal = galactose, Glc = glucose. Results are mean and SEM across three biological replicates. B) Transcriptomics identifies CAZymes involved in okara degradation by *N. intermedia.* Results are shown as the log2foldchange compared to a no-carbon control. Green indicates significantly differentially expressed genes. Purple indicates a predicted CAZyme. The blue border around the circle indicates a gene significantly induced above the log2fold change threshold of 4, which captured 10 CAZymes and 21 genes with other predicted functions. C) Expression of the genes that were most highly induced on okara (log2foldchange>4) across purified carbon sources of decreasing complexity. As in B), purple indicates a predicted CAZyme. The substrates were chosen based on their presence in soybeans or okara polymers. Clustering of the data revealed the existence of four separate clusters displaying distinct expression patterns. D) Sugar analysis of the hydrolyzed pectin-containing fraction in okara samples subjected to SSF by *N. intermedia* indicated consumption of both arabinose and galactose. Results are mean and SEM on a dry weight basis from three biological replicates. E) Cellulase activity across different carbon sources in liquid cultures. Oka=okara, Avi=avicel, Suc=Sucrose, Rhg= Rhamnogalacturonan. Results are mean and SEM from three biological replicates. E) Cellulase activity detected in okara subjected to SSF by *N. intermedia,* indicating active fungal secretion during the fermentation process. Results are mean and SEM from three biological replicates.

Okara dramatically changed the global gene expression profile, including significantly upregulating several predicted CAZymes (Fig. 2B). Above a cutoff of log 2-fold change > 4, which we reasoned would capture the most highly expressed genes in response to okara^59^, 10 of these CAZymes remained, in addition to 21 other genes with diverse predicted functions (Fig. 2B, Table S4, and Supplementary Data S1). However, as this is the first transcriptomics effort for enzyme identification in this fungus, and because *N. intermedia* is poorly characterized, assigning these genes to specific metabolic pathways or substrates within the complex okara biomass is challenging using computational prediction alone. To better understand which carbon source within okara was consumed and to better assign the function of the upregulated genes, we repeated the transcriptomics experiment with more defined substrates that were part of the complex okara substrate^58^. Specifically, we exposed the strain to purified mono-, di-, and polysaccharide components of okara, as the sole carbon source (n=12 unique substrates) and then mapped the top genes overexpressed on okara to the transcriptomics data generated on the purified substrates (Fig. 2C). By comparing the genes most upregulated on okara with the genes upregulated on the defined substrates, we sought to get a better look at what specific molecules *N. intermedia* was degrading in the complex plant biomass, while also linking predicted CAZymes to individual molecules^59^.

Initial heatmap analysis using hierarchical clustering revealed a diversity of genes that were either shared with the purified substrates or unique to okara (Fig. 2B and Supplementary Data S1). These genes fell into four groups. For example, group 1 included a predicted pectate lyase (#357033), ribose/galactose isomerase (#520753), beta-galactosidase (#432986), and reductase (#490212). These genes were not only induced on the pectin-rich soluble soy polysaccharides^57^ and purified rhamnogalacturonan, the main pectic fiber found in soy, but also on monosaccharides arabinose, galactose and glucuronic acid. This strongly suggested pectin degradation and potential uptake of released sugars during okara fermentation. Group 2 included a set of five genes that were shared between okara and Avicel (purified cellulose), including three predicted hydrolytic cellulase enzymes (#549776, #480919, and #506499), a lytic monooxygenase predicted to bind cellulose (#53068), and a carbohydrate esterase (#53091). Group 3 included genes mainly predicted to be involved in amino acid metabolism and were shared with cellobiose, mannose, and xylose. Finally, group 4 included three glycosyl hydrolases (#442209, #38386, #516864), predicted proteins with poorly understood functions, or hypothetical proteins. It is notable that some genes induced by okara were also induced by simpler sugars included in the screen, suggesting that simple substrates can induce diverse pathways in *N. intermedia*, as has been observed in the related model ascomycete mold *N. crassa* ^59^ (Fig. 2C).

Co-expression analysis across all conditions confirmed a shared transcriptional module between okara and Avicel as well as a cluster of genes that were unique to okara (Fig. S8). While the shared Avicel and okara cluster contained mainly CAZymes predicted to break down cellulose, the okara cluster harbored a diversity of genes, including predicted pathways for the degradation of phenylalanine and leucine, two of the most abundant amino acids in okara^60^, enzymes involved in the metabolism of other amino acids, and genes such as CAZymes and oxidoreductases with unknown functions (Fig. S8 and Supplementary Data S1). While we did not detect major changes in the global amino acid composition (Fig. S5), these data highlight that *N. intermedia* upregulates predicted pathways for amino acid degradation. In addition to providing a first genomic and biochemical look at okara degradation by *N. intermedia,* these results improve annotation of *N. intermedia* metabolic genes and more broadly uncovers the inner workings of nutrient utilization by this edible fungus.

To biochemically validate the transcriptomics observations, we focused on pectin and cellulose degradation, as these were not only strongly suggested by the transcriptomics substrate comparison but could also be broadly relevant to upcycling of diverse plant-based byproducts for human consumption. To validate the pectin degradation and potential consumption of pectin-bound simple sugars such as galactose, arabinose, and galacturonic acid, we analyzed the sugar composition in the hydrolyzed pectin fraction of okara before and after SSF. These analyses revealed a clear decrease in both arabinose (60% decrease) and galactose (40% decrease), suggesting release of pectin-bound sugars (Fig. 2D and Fig. S9). Liquid cultures using okara as the only carbon source confirmed that arabinose and galactose were initially released to the medium and then taken up by the fungus, with only a small amount remaining, suggesting direct consumption during the fermentation (Fig. S10). These two sugars are abundant in rhamnogalacturonan, the main pectic fiber in soy, and the position of these sugars within the side chain of the polymer makes them accessible for hydrolytic release and microbial degradation, which is consistent with our data^57,61^. We did not detect a significant decrease of galacturonic acid in SSF, which is found in more inaccessible portions of the pectin polymer^61^, or in any other sugar (Fig. S9).

We next turned to validating cellulose degradation, which was suggested by the shared transcriptional module between okara and Avicel (Fig. 2C, Fig. S8, and Supplementary Data S1). To validate the presence of the predicted cellulose-degradation machinery, we assayed the extracellular release of cellulases, as a broad suite of these CAZymes were predicted by the transcriptomics data. Enzymatic assays on culture supernatants confirmed that cellulases were highly produced during growth on Avicel and okara as the sole carbon source, but not on sucrose or rhamnogalacturonan, which were included as negative controls (Fig. 2E). In line with the transcriptomics data, this observation highlights a specificity in *N. intermedia* substrate sensing and enzymatic response. Notably, we also detected high levels of cellulase activity in okara fermented by *N. intermedia* through SSF, and this activity was absent in the starting material, suggesting active fungal secretion during fermentation (Fig. 2F). Taken together, these data provide direct biochemical evidence of some of the activities that were predicted by gene expression profiles and establish both active pectin and cellulose degradation by *N. intermedia* in the metabolism of okara. While cellulose degradation may explain the release of free glucose, pectin metabolism could account for the release of both arabinose and galactose during SSF.

### *Neurospora intermedia* strains used for oncom belong to a genetically distinct subpopulation that is associated with byproducts of human activity

Many fungi traditionally used for food production display genomic adaptations to the substrate on which they are typically grown^62–65^, leading us to wonder about the origins and evolutionary history of *N. intermedia* strains used for oncom. Previous isolation efforts of *N. intermedia* across the globe have indicated the existence of two, distinct types of strains that have macroscopic differences in conidial (asexual spore) pigmentation and size^31^. One type has orange, smaller conidia, the other type has large macroconidia that are yellow-orange in color. Interestingly, the occurrence of these strains strongly correlates with their ecological origin^31^. The orange type has been frequently isolated from burned vegetation, a common site for *Neurospora* isolation. The yellow type, in contrast, has only been isolated from non-burned substrates, including oncom^30,31^. However, any genetic and biochemical differences between the two types have remained unclear.

We sought to understand the genetic basis of strain-level variation among *N. intermedia*. Pan-genome analysis of 29 genomes generated between this study (8 strains) and an unrelated study^66^ revealed that 90% of genes were shared (core genome n=7751 genes) (Fig. 3A and Fig. S11). Using coding sequences of shared *Neurospora* genes, we constructed a comprehensive genome-wide phylogeny which indicated that *N. intermedia* is genetically distinct from other sequenced *Neurospora* species, including the model fungus *N. crassa* (Fig. 3B). Strikingly, *N. intermedia* strains were split into two distinct, closely related groups that segregated based on ecological but not on geographical origin (Fig. 3B and Table S5). Clade A harbored strains isolated from burned vegetation, while Clade B harbored all oncom strains (n=6), as well as strains isolated from fiber-rich byproducts, including sugar cane bagasse, a cellulose-rich byproduct of sugar production, and leftover corn cobs. It is noteworthy that the oncom-derived strain #2559 and burned vegetation-derived strain #8767 segregated in different clades while they were both isolated from the town of Bogor, on the island of Java, Indonesia, reinforcing a genetic rather than geographical distinction between the two clades (Fig. 3B and Table S5). Strains from clades A and B had the previously reported difference in conidial pigmentation, suggesting that they mirror the previously reported orange and yellow strain types (Fig. S12) ^31^. This provides conclusive evidence for the existence of two genetically distinct *N. intermedia* sub-groups and that oncom strains are closely related to other strains associated with byproducts of human activity.

**Figure 3.**
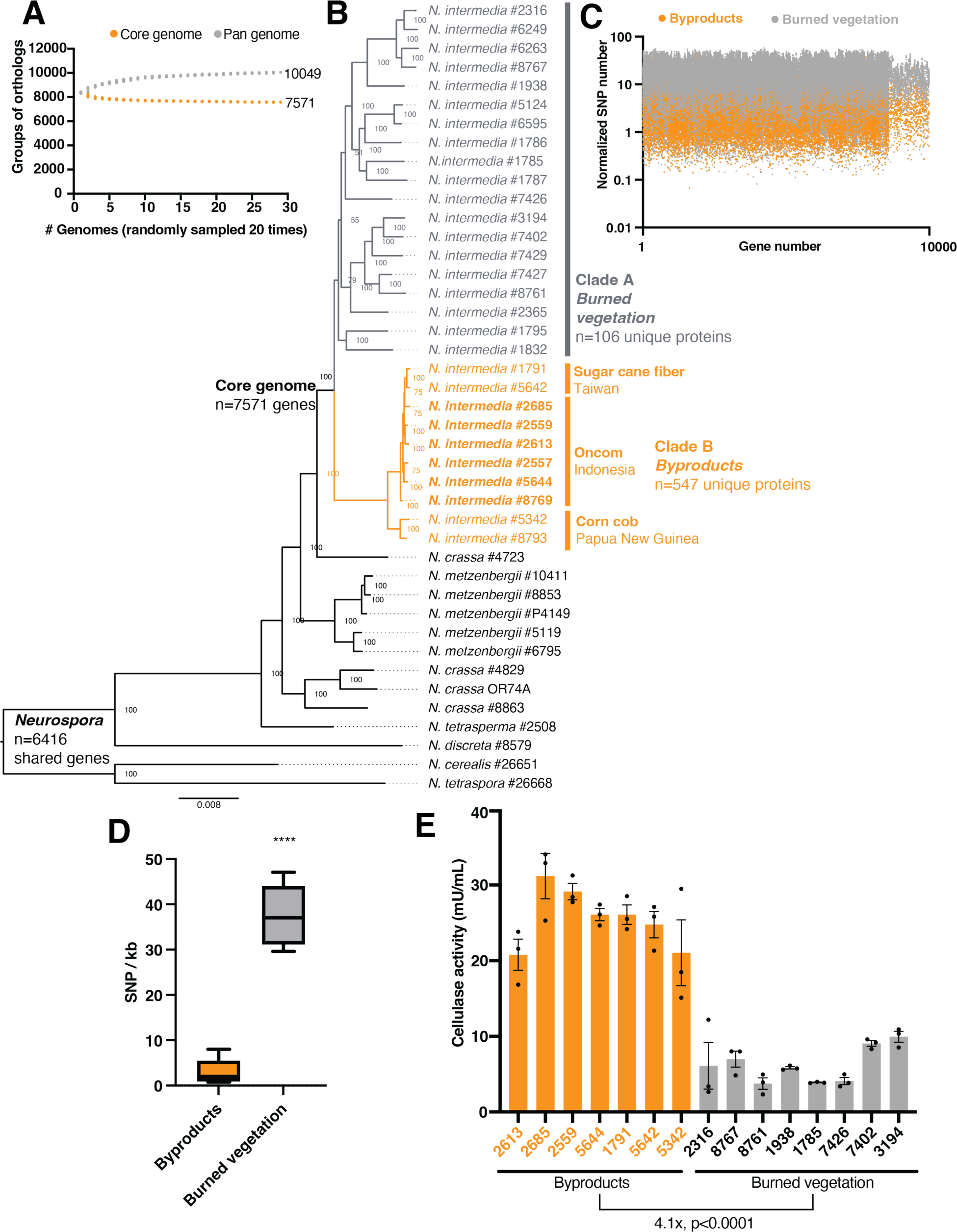
*Neurospora intermedia* strains used for oncom belong to a genetically distinct subpopulation that is associated with byproducts of human activity. A) Core/Pan genome analysis for *N. intermedia*. Grey and orange circles show the number of protein families in the pan and core genomes for a subset of *n* strains (x axis). This calculation was done 20 times for a random sample of *n* genomes. The number of core and unique genes across all strains are shown alongside the curves. B) Phylogenetic tree of *Neurospora* species based on shared coding genes (n=6416). Clade A was associated with burned vegetation, a common site for isolating *Neurospora* species. Clade B included all oncom-derived strains (highlighted in bold) collected from Indonesia, as well as strains isolated from sugar cane bagasse (#1792, #5642), the fibrous byproduct of sugar production, or leftover corn cobs (#5342, #8793). Oncom-derived strain #2559 and burned strain #8767 were both isolated from the town of Bogor, on the island of Java, Indonesia, reinforcing a genetic rather than geographical distinction between the two clades. C) Analysis of single-nucleotide polymorphisms (SNPs) across all genes in *N. intermedia* strains. SNPs were fixed to the reference strain #2613 and were mapped across all shared genes among the remaining 28 strains. Results were normalized to gene length. D) The number of SNPs per kb in the predicted cellulase (#480919, glycosyl hydrolase family 7) that was highly expressed on both okara and Avicel. The intervals in the box and whiskers plot represent the min, max, and the median number of mutations across the groups. E) Secreted cellulase activity between burn-associated and byproduct-associated during growth on okara. Results are mean and SEM across three biological replicates. There was a 4.1 times lower mean secreted cellulase activity among burn-associated strains compared to byproduct-associated strains (****=p<0.0001, unpaired t-test).

Once we established phylogenetic relationships among *N. intermedia* strains, we used comparative and population genomics analyzes to search for genomic features associated with the byproduct-associated lifestyle. The byproduct and burn-associated clades showed both differences in gene content and mutations in shared coding genes. For instance, the pan-genome of the byproduct-associated strains (n=741 genes) was larger compared to that of burn-associated strains (n=132 genes) and was enriched in a range of predicted metabolic functions, including hydrolase activity (n=24 genes, the most enriched) as well as oxidoreductases, transporters, catalytic activity acting on protein, transferases, ATP-dependence, and DNA binding (Fig. S11). However, only the functions of a few genes could be predicted. Furthermore, the genome-wide phylogenetic tree strongly suggested that the observed divergence may arise from differences in shared genes (Fig. 3B). To understand these potential changes, we identified and analyzed single nucleotide polymorphisms (SNP) in coding sequences among all strains, using the oncom-derived strain #2613 as a reference (Supplementary Data S2). This analysis revealed that the two clades represent two distinct subpopulations, with the burn-associated strains clearly segregating from the waste associated strains by their SNP frequency on a genome-wide level (Fig. 3C). The genes harboring the most SNPs per kb (n=209, >2 standard deviations beyond the genome’s average mutation rate) included transcription factors, transporters, an ergothioneine biosynthetic enzyme, a polyketide synthase, a predicted plant toxin, as well as several CAZymes, including those predicted to be involved in lignin and cellulose deconstruction (Supplementary Data S2). The SNP pattern was generally similar across all waste-associated strains, despite their diverse geographical origins from different islands (Taiwan, Papua New Guinea, and Indonesia) and association with different substrates (Fig. 3C). These results demonstrate a strong genomic signature associated with byproducts of human activity.

Cross-referencing the genes harboring the most SNPs with the full transcriptomics dataset identified a highly mutated predicted cellulase (Glycosyl Hydrolase family 7, JGI protein ID #48919 in Fig. 3C and Table S4) that was highly expressed on both okara and Avicel (Fig. 2C, Fig. S13, and Supplementary Data S2). Across all strains, the burn-associated group had an average of 37 SNPs per kb in this enzyme, compared to the <3 for the waste-group (Figure 3D, p<0.0001, unpaired t-test). Structural modeling located the mutations away from the predicted active site (Fig. S14), suggesting that the mutations could impact cellulose degradation by affecting substrate binding or assembly of complexes that are critical for efficient cellulose metabolism in fungi^67–69^. To assess whether the two clades of *N. intermedia* also harbored associated differences in active cellulose degradation, we assayed the secreted cellulase activities of a subset of strains from each group (n=7 byproduct-associated strains, n=8 burn-associated strains). Consistent with the genomic data, we found that secreted cellulase activity was >4-fold higher among the waste associated strains during growth on okara (Fig. 3E, p<0.0001, unpaired t-test). There was no difference in overall biomass yield of the two groups of strains (Fig. S15). This indicates that the elevated cellulase activity is explained by biochemical rather than growth differences, and that there is a nutritional redundancy in nutrient utilization under these conditions as higher enzyme activity did not translate into elevated growth. While we cannot conclude that the increased cellulase activity directly results from a mutation in the cellulase enzyme without further genetic experiments, these results highlight that the two ecologically and evolutionarily distinct subpopulations of *N. intermedia* display genetic and biochemical differences in metabolic enzymes that are active during growth on byproducts.

### *Neurospora intermedia* grows on diverse industrial byproducts from food and agricultural processing

The observation that the strains used for oncom production belong to a genetically distinct waste-associated subpopulation raised questions about the broader ability of *N. intermedia* to upcycle plant-based byproducts beyond okara for applications in non-traditional contexts. To evaluate this possibility, we assembled a library of n=30 industrially relevant byproduct substrates, including plant-based milk byproducts, fruit and vegetable byproducts, grain and agricultural processing byproducts, brewing byproducts, oilseed presscakes, and others. We then conducted a SSF screen of oncom-derived *N. intermedia* #2613 to assess its growth across these substrates of diverse origins (Supplementary Data S3). Following 72 h of SSF of inoculated substrates, we found that *N. intermedia* grew on all byproducts but grape pomace, olive pomace, and buckwheat hulls, as assessed by the development of mycelia and conidia (Fig. 4). However, the extent of growth varied between substrates. There was a moderate amount of growth on coffee grounds, almond hulls, banana peels, pineapple core, oat bran, and almond milk byproduct, while only minimal growth was observed on hazelnut skins and spent grain (Fig. 4 and Table S6). All others supported a high level of growth comparable to that observed on okara, the traditionally used substrate. Upon analyzing the protein content of a subset of byproducts where *N. intermedia* displayed high growth in SSF, we found an increase in protein content as a result of the fermentation (Table S7). This indicates that *N. intermedia* can alter the nutritional value of substrates beyond okara. While poorly characterized *N. intermedia* strains have been previously demonstrated to grow on the byproducts peanut press cake and stale bread in SSF^21,70^, these results illuminate the broad substrate range of oncom-derived *N. intermedia* and suggest potential opportunities for upcycling across diverse byproducts.

**Figure 4.**
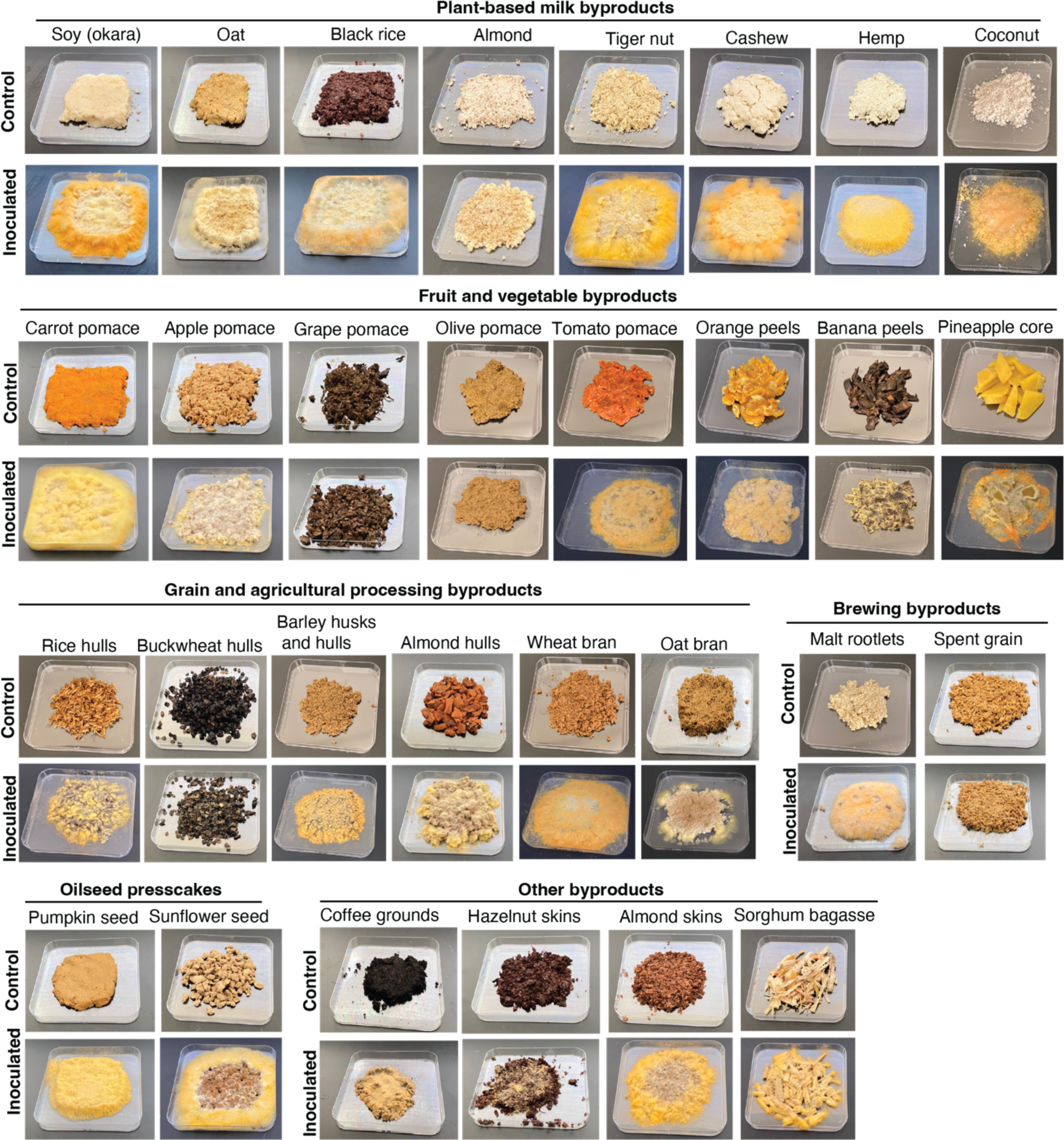
*Neurospora intermedia* grows on diverse industrial byproducts from food and agricultural processing. SSF screen of the oncom-derived *N. intermedia* #2613 growth across diverse industrially relevant byproducts. Okara was included as a positive control and benchmark for the fungal growth. *N. intermedia* grew on all substrates tested except for grape pomace, olive pomace, and buckwheat hulls, as assessed by the development of mycelia and / or conidia. Growth scores were assigned to each substrate and are shown in Table S6. Large photos from controls and inoculated substrates can be found in Supplementary Data S3.

### *Neurospora intermedia* does not encode for any known mycotoxins and is positively perceived outside Indonesia

The growth on diverse byproducts available across a variety of geographical locations suggests promise for *N. intermedia* as a fungus for waste-to-food conversion for new applications beyond red oncom production. However, any such future applications must account for potential issues surrounding the safety (mycotoxin production) as well as the palatability of *Neurospora* fermentations among consumers outside the traditional setting of Java, Indonesia.

We first investigated the potential for toxin production. A major concern with fungal fermentation is the potential production of secondary metabolites such as mycotoxins that could be harmful to human health^71^. While many of these secondary metabolites may not be produced on the substrates typically used for fermentation, the genomic potential to produce these are concerning when exposing fungi to new substrates or growth conditions^71,72^. As a first step towards assessing the broader safety profile of *N. intermedia*, we analyzed the capacity for secondary metabolite production using our reference genome of strain #2613 (Fig. 5A and Table S8). Compared to other sequenced, edible Ascomycete fungi frequently used for food production, including in SSF upcycling, *N. intermedia* encoded for a strikingly small number of predicted natural products. For example, *Aspergillus oryzae* (koji mold), a mold that is frequently explored for byproduct SSF across scientific and gastronomic contexts^12,24^, harbored >100 natural product gene clusters, while *N. intermedia* encoded for only 22, with 8 of these being predicted polyketide synthases and 9 being predicted non-ribosomal-peptide synthases.

**Figure 5.**
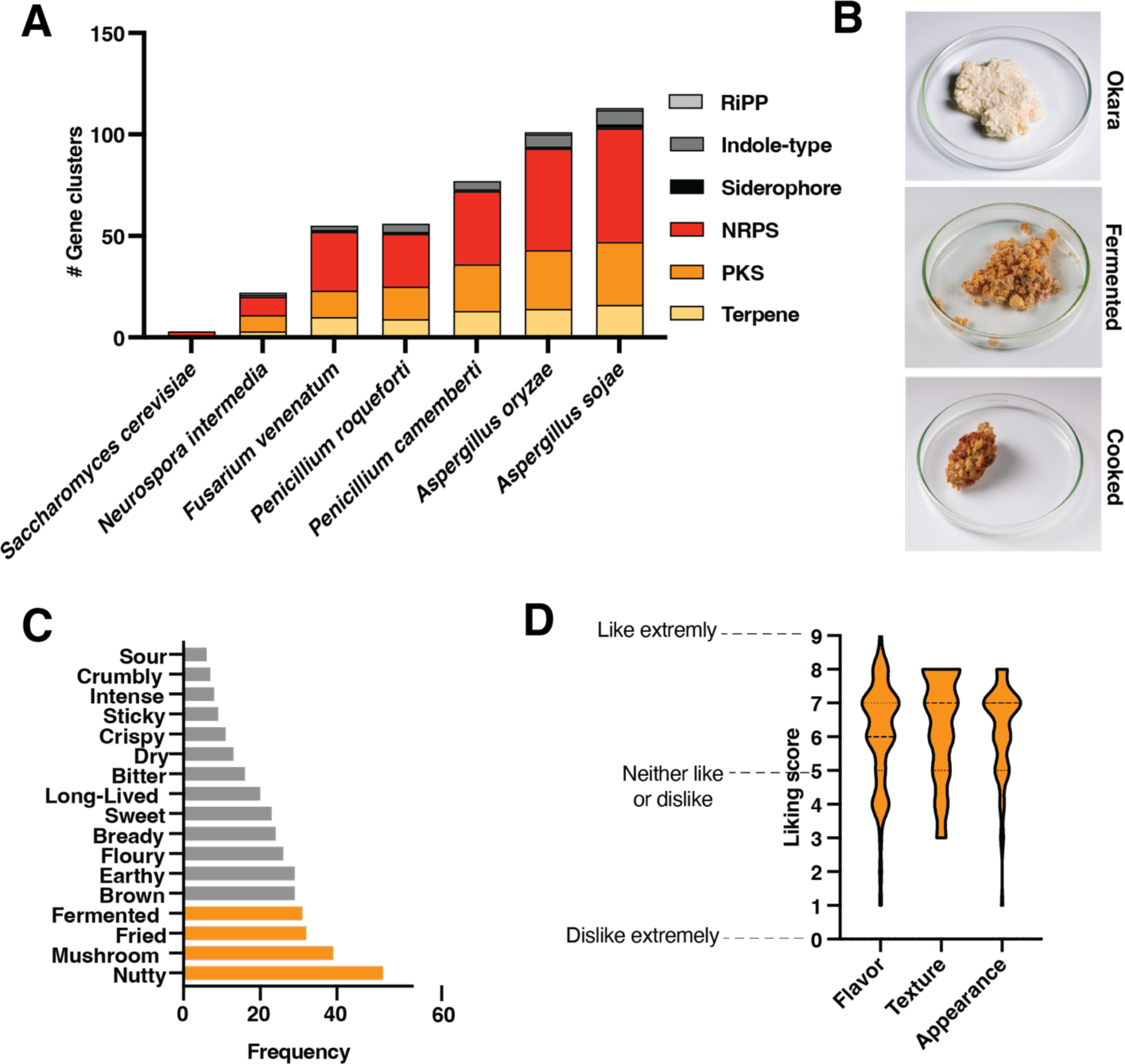
*Neurospora intermedia* has low potential for secondary metabolite production and is positively perceived by consumers outside Indonesia. A) Predicted natural products production capacity of *N. intermedia* and other sequenced Ascomycete molds used for food production. RiPP = ribosomally synthesized and post-translationally modified peptide; NRPS = nonribosomal peptide synthase; PKS = polyketide synthase. Compared to other ascomycete molds used for food production, *N. intermedia* encodes for a small number of predicted natural products (n=22) and does not have the genomic capacity to produce any known mycotoxins. 8 of these were predicted polyketide synthases, and 9 were predicted non-ribosomal-peptide synthases. The full results of the analysis are shown in Table S8. B) preparation of fermented, pan-fried okara using *N. intermedia.* The pan-fried okara was used in sensory trials C) Results of a Check-All-That-Apply (CATA) sensory analysis with n=61 adult Danish participants who had never tried oncom. Okara was subjected to fermentation by *N. intermedia* followed by a CATA evaluation of the cooked product. Sensory attributes assigned by >10% of participants are shown in the graph, and those characteristics assigned by >50% of participants are highlighted in orange. D) Liking score by the same sensory panel tasting the fermented, cooked okara. Liking scores are shown on the y-axis. 0=Dislike extremely, 1=Dislike extremely, 2=Dislike very much, 3=Dislike moderately, 4=Dislike slightly, 5=Neither like nor dislike, 6=Like slightly, 7=Like moderately, 8=Like very much, 9=Like extremely. The sensory panel scored the flavor, texture, and appearance was rated >6, indicating a liking towards the product. Violin plots show distribution of ratings for each category for the n=61 participants. Results +/− SEM of flavor, texture, appearance were 6.1+/−0.2, 6.3+/−0.2, and 6.2+/−0.2, respectively.

In terms of the number of gene clusters, *N. intermedia* was most similar to *Saccharomyces cerevisiae,* a single-cell yeast used for bread and brewing that is known for its safety. Unlike *Fusarium venenatum,* a fungus used for alternative meat production in Quorn^73^, *and Aspergillus and Penicillium* molds used for fermented foods^62,72^, *N. intermedia* did not encode for any known mycotoxins. Additionally, we could not detect active production of any known mycotoxins or natural products during SSF of okara by *N. intermedia* using untargeted metabolomics (see methods for details). While the *Neurospora* genus has been proposed as safe based on historical evidence and its traditional use in oncom^49^, these results provide initial genomic evidence for *N. intermedia* safety profile. Interestingly, *N. intermedia* is predicted to encode for the same number and type of natural products as *N. crassa*, which was recently demonstrated to be safe as food in extensive animal trials^74^.

Ultimately, the broader potential of *N. intermedia* for upcycling will depend on sensory appeal outside the traditional context in new applications and settings. As a starting point to assess this potential, we conducted a Check-All-That-Apply (CATA) consumer survey^75,76^ of fermented, cooked okara among Danish consumers (n=61 participants) who had never tried oncom previously (Fig. 5B). These participants were asked to assign a set of attributes (negative and positive descriptors) and indicate their liking of the food, which was presented to them as a pan-fried product (Table S9). These participants assigned mainly positive attributes to the cooked product, characterizing it as mainly *nutty, mushroom, fermented, brown*, and *earthy* (attributes selected by >50% of respondents) (Fig. 5C). Importantly, the consumers also overall liked the appearance, flavor, and texture of the cooked product (Fig. 5D, mean liking score >6 out of 9 across all flavor, texture, and appearance). The favorable rating is in stark contrast to consumer attitudes towards okara, which has a relatively low acceptance threshold even when used as a minor ingredient^77,78^. Both the liking score and the positive attributes suggest that *N. intermedia* byproduct fermentation can create foods that are liked outside the context where red oncom is traditionally produced and consumed, highlighting the potential of *N. intermedia* for fungal waste-to-food upcycling in new contexts and applications.

## Discussion

Fungi have promise for upcycling of food waste for increased sustainability and food security, particularly in the conversion of byproducts into human food through solid state fermentation (SSF) ^1,7,14^. However, engineering fungal byproduct transformation for specific outcomes remains challenging, in part due to the limited availability of sequenced, characterized, edible fungal strains, and our poor understanding of fungal waste-to-food conversion at the genetic and molecular level. Here we used multi-omics analysis of oncom – a traditional, Indonesian fermented food produced from readily available byproducts – as a model system to identify strains, enzymes, and processes that could enable byproduct upcycling in SSF.

We initially surveyed black and red oncom microbial communities using metagenomics, but focused further efforts on red oncom due to its reduced microbial complexity, the dominance of *N. intermedia* as the major fungus involved in the fermentation, and the fact that red oncom is a unique traditional food involving intentional cultivation of *Neurospora*^49^. We conclude that oncom-derived *N. intermedia* changes the biochemical and aroma composition of okara, utilizes pectin and cellulose degradation during substrate breakdown, and belongs to a distinct subpopulation that is associated with plant-based byproducts. These byproduct-associated strains harbor genetic and biochemical differences in cellulose degradation, among other genomic changes. We also uncovered that *N. intermedia* readily grows on diverse industrial byproducts, does not encode for any known mycotoxins, and can produce food from waste that is positively perceived among consumers outside of Indonesia. Overall, this study enhances our understanding of red oncom production at the genetic, biochemical and microbial levels, and establishes *Neurospora intermedia* as a promising fungus for waste-to-food upcycling.

Our study leaves several key questions unanswered, creating opportunities for further investigation. First, the microbial and genetic basis of black oncom remains poorly characterized and needs to be explored further. Additionally, although the complete dominance of *Neurospora* in some red oncom samples suggest that this fungus may be sufficient for making oncom, other microbes detected in some samples could play unappreciated roles in the fermentation process and the final product, which warrants further investigation. It is also unclear from our data whether the dominance of *Neurospora* among fungi arises from its rapid growth, outcompeting other organisms, or from active killing and inhibition of other organisms, raising questions about microbe-microbe interactions during the fermentation. Finally, while we observed differences in genes and enzymatic activities related to cellulose degradation between the burn-associated and byproduct-associated subpopulations of *N. intermedia*, genetic experiments are needed to establish whether the enhanced cellulase activity among byproduct-associated *N. intermedia* originate from the mutations in the predicted cellulase rather than other broader genome-wide changes. Mechanistically characterizing the genes, enzymes, and SNPs that distinguish the two *N. intermedia* subpopulations is therefore an important avenue for further study. The close relatedness between *N. intermedia* and *N. crassa,* a model fungus that has decades of fundamental genetic work and widely available genetic tools and protocols^79,80^, may make *N. intermedia* particularly amenable to such genetic investigations in the future.

In addition to providing insight into oncom, our study sheds light on human domestication of microorganisms for sustainability challenges. Although many fungi have been domesticated from their wild relatives for food production throughout human history, *Neurospora intermedia* stands out among characterized fungi in that it appears to have been uniquely recruited for waste-to-food conversion^62–65,81,82^. Recent work has revealed that wild *Neurospora* strains likely exist as plant endophytes^83,84^, and we propose that the genomic changes among byproduct-associated *N. intermedia* may reflect a transition to a new lifestyle and ecological niche associated with the human generation of byproducts. The observed genomic and biochemical changes among byproduct-associated *N. intermedia* strains are consistent with observations from other fungi that have adapted to particular substrates as a result of human selection in food production^62,64,82^. For example, compared to its close relatives, the edible fungus *A. oryzae –* traditionally used in shoyu, sake, and miso production in East Asia – displays increases in hydrolase capacity, mutations in key genomic regions involved in secreted enzymes, and has increased secreted amylase activity which supports its growth on grains and legumes in traditional fermented foods^62,81,85^.

We envision several possible applications for *N. intermedia* in waste-to-food conversion moving forward. In addition to its low capacity for secondary metabolite production and ability to grow on diverse complex plant-derived substrates, *N. intermedia* is a particularly fast-growing fungus^86^, which may give it advantages for SSF upcycling compared to many mushroom-forming fungi that are also able to break down complex plant substrates in upcycling but suffer from slow growth under cultivation conditions^11^. Working with chefs, we have recently found that *N. intermedia* can be readily used to create new fermented foods from non-waste substrates across different applications^87^. A separate study recently revealed that poorly characterized *N. intermedia* can be used in SSF of stale bread to produce a burger-like product that is positively perceived among Western consumers^20^. We anticipate that these previous findings, alongside our unique dataset of growth on diverse byproducts presented in this study, could guide the creation of new foods from waste using *N. intermedia* SSF in the near future. Additionally, we have also recently found that *N. intermedia* CAZymes can transform flavor and texture in fermented foods^87^, suggesting that the diversity of CAZymes encoded and expressed by the edible *N. intermedia* may have applications in food processing across culinary and industrial applications. Longer term, we envision that the robust growth of *N. intermedia* on byproducts could make it suitable for production of biomass for sustainable meat alternatives or as a host for secreted protein production, where growth on byproducts instead of highly purified substrates as glucose is essential for decreasing costs to alleviate the environmental burden of industrial animal agriculture^9,13^. However, future applications involving *N. intermedia* for waste conversion in new contexts will benefit from improved tools to understand and manipulate its genome and metabolism, assessment of potential life cycle impacts, as well as overcoming potential challenges in scale up, consumer acceptance, and the regulatory landscape.

Future applications must also consider the origins of *N. intermedia* and oncom as indigenous biotechnologies. This study serves as another reminder that traditional foods may hold clues to addressing global challenges in sustainability^3,11,88,89^ and highlights the importance of preserving and characterizing these traditions in the face of rapid global change and food system industrialization. While such traditional biological resources could benefit the world, it is critical that new exploration and innovations centered on these practices procure benefits for the original inventors moving forward.

## Acknowledgements

We thank Dr Jennifer Wang at the Harvard Center for Mass Spectrometry for technical support with aroma analysis; Daniela Rago from the analytical lab at Novo Nordisk Foundation Center for Biosustainability for technical support and LC-MS/MS instruments operation; **and** the Scheller group and Yi-Chun Chen at the Joint Bioenergy Institute for help with HPAEC assay. This work was part of the DOE Joint BioEnergy Institute (https://www.jbei.org) supported by the U.S. Department of Energy, Office of Science, Office of Biological and Environmental Research under contract <u>DE-AC02-05CH11231</u> between Lawrence Berkeley National Laboratory and the U.S. Department of Energy, and by the Philomathia Foundation. VMR was supported by the Miller Institute at University of California, Berkeley. PCM and ACC were supported by Novo Nordisk Foundation grant no. NNF20CC0035580. JMVE acknowledges funding from m-CAFEs Microbial Community Analysis & Functional Evaluation in Soils project, a Science Focus Area led by Lawrence Berkeley National Laboratory and supported by the U.S. Department of Energy, Office of Science, Office of Biological & Environmental Research under contract number DE-AC02-05CH11231. The specific work (proposal: 10.46936/10.25585/60001019) conducted by the U.S. Department of Energy Joint Genome Institute (https://ror.org/04xm1d337), a DOE Office of Science User Facility, is supported by the Office of Science of the U.S. Department of Energy operated under Contract No. DE-AC02-05CH11231. Finally, we thank the restaurants and industry contributors who donated diverse byproducts for the evaluation of *N. intermedia* growth, and professors John Taylor and Louise Glass for helpful discussion.

## Author contributions

VMR conducted all experiments and wrote the manuscript and conceptualized the study together with JDK. JMVE conducted bioinformatics analysis of metagenomics and transcriptomics data. NRV, MOG, DPV, RM conducted sensory trials. AR conducted the solid state fermentation screen. NRV and PMS conducted volatile aroma analysis. ID, LN, CHW, collected and analyzed physiochemical properties of oncom samples from local producers and cataloged the production process by artisans in Java, Indonesia. EEKB contributed to metabolite analysis. ACC conducted untargeted metabolomics analysis of natural products and mycotoxins. RR, AL, GH, MY, SH, CD, YY, VN, IVG conducted whole-genome sequencing, assembly, and annotation of *N. intermedia* FGSC #2613, as well as transcriptomics sequencing, at the Joint Genome Institute. PCM conducted all genomic and phylogenetic analyses.

## Data availability

The genome assembly and annotation of *Neurospora intermedia* FGSC #2613 is available through the MycoCosm portal at https://mycocosm.jgi.doe.gov/Neuin1 and GenBank (available upon publication). The genomic reads have also been deposited to the Sequence Read Archive (SRA) under #1294447. Additionally, the transcriptomics data from *Neurospora intermedia* FGSC #2613 grown across carbon source has been deposited to the Sequence Read Archive (SRA); the specific access information is specified in Table S10 in the supplemental materials. The genomes and reads for *N. intermedia* strains #1791, #2557, #2559, #2685, #5342, #5642, and #5644 are deposited at the Genbank under Bioproject PRJNA996151; the sequencing information can be found in Table S11 in the supplemental materials. The 16s and ITS amplicon sequencing, as well as the metagenome sequencing, has also been deposited at the Genbank under Bioproject PRJNA996151.

## References

1 Springmann, M. et al. Options for keeping the food system within environmental limits. Nature 562, 519–525 (2018). 10.1038/s41586-018-0594-0

2 van Zanten, H. H. E. et al. Circularity in Europe strengthens the sustainability of the global food system. Nature Food 4, 320–330 (2023). 10.1038/s43016-023-00734-9

3 Jahn, L. J., Rekdal, V. M. & Sommer, M. O. A. Microbial foods for improving human and planetary health. Cell 186, 469–478 (2023). 10.1016/j.cell.2022.12.002

4 Zhu, J. et al. Cradle-to-grave emissions from food loss and waste represent half of total greenhouse gas emissions from food systems. Nature Food 4, 247–256 (2023). 10.1038/s43016-023-00710-3

5 Service, U. E. R. Estimates of Food Loss at the Retail and Consumer Levels, <https://www.ers.usda.gov/data-products/food-availability-per-capita-data-system/food-loss/> (2020).

6 Zhaojun Wang, T. G., Zhiyong He, Maomao Zeng, Fang Qin, Jie Chen. Reduction of off-flavor volatile compounds in okara by fermentation with four edible fungi. LWT 155 (2022). 10.1016/j.lwt.2021.112941

7 Sandström, V. et al. Food system by-products upcycled in livestock and aquaculture feeds can increase global food supply. Nature Food 3, 729–740 (2022). 10.1038/s43016-022-00589-6

8 Reynolds, C. J., Piantadosi, J. & Boland, J. Rescuing Food from the Organics Waste Stream to Feed the Food Insecure: An Economic and Environmental Assessment of Australian Food Rescue Operations Using Environmentally Extended Waste Input-Output Analysis. Sustainability 7, 4707–4726 (2015).

9 Humpenoder, F. et al. Projected environmental benefits of replacing beef with microbial protein. Nature 605, 90–96 (2022). 10.1038/s41586-022-04629-w

10 Protzko, R. J. et al. Engineering Saccharomyces cerevisiae for co-utilization of d-galacturonic acid and d-glucose from citrus peel waste. Nature Communications 9, 5059 (2018). 10.1038/s41467-018-07589-w

11 McBee, R. M. et al. Engineering living and regenerative fungal-bacterial biocomposite structures. Nat Mater 21, 471–478 (2022). 10.1038/s41563-021-01123-y

12 B. Noval, D. P. Manual para la revalorización de descartes. (Basque Culinary Center, http://www.bculinarylab.com/2018/12/23/la-gastronomia-como-via-del-desarrollo-sostenible/, 2018).

13 Jarvio, N. et al. Ovalbumin production using Trichoderma reesei culture and low-carbon energy could mitigate the environmental impacts of chicken-egg-derived ovalbumin. Nat Food 2, 1005–1013 (2021). 10.1038/s43016-021-00418-2

14 Javourez, U., Rosero Delgado, E. A. & Hamelin, L. Upgrading agrifood co-products via solid fermentation yields environmental benefits under specific conditions only. Nat Food 3, 911–920 (2022). 10.1038/s43016-022-00621-9

15 Piercy, E. et al. A sustainable waste-to-protein system to maximise waste resource utilisation for developing food- and feed-grade protein solutions. Green Chemistry 25, 808–832 (2023). 10.1039/D2GC03095K

16 Sastraatmadja, D. D., Tomita, F. & Kasai, T. Production of High-Quality Oncom, a Traditional Indonesian Fermented Food, by the Inoculation with Selected Mold Strains in the Form of Pure Culture and Solid Inoculum. J Grad Sch Agr Hokkaido Univ 70, 111–127 (2002). 10.11501/3178528

17 Dirar, H. A. Kawal, Meat Substitute from Fermented Cassia-Obtusifolia Leaves. Econ Bot 38, 342–349 (1984). Doi 10.1007/Bf02859013

18 Wang, J. et al. Fungal solid-state fermentation of crops and their by-products to obtain protein resources: The next frontier of food industry. Trends in Food Science & Technology 138, 628–644 (2023). 10.1016/j.tifs.2023.06.020

19 Wang, J. et al. Solid-State Fermentation of Soybean Meal with Edible Mushroom Mycelium to Improve Its Nutritional, Antioxidant Capacities and Physicochemical Properties. Fermentation 9, 322 (2023).

20 Hellwig, C., Gmoser, R., Lundin, M., Taherzadeh, M. J. & Rousta, K. Fungi Burger from Stale Bread? A Case Study on Perceptions of a Novel Protein-Rich Food Product Made from an Edible Fungus. Foods 9 (2020). 10.3390/foods9081112

21 Gmoser, R. et al. From stale bread and brewers spent grain to a new food source using edible filamentous fungi. Bioengineered 11, 582–598 (2020). 10.1080/21655979.2020.1768694

22 Šelo, G. et al. Bioconversion of Grape Pomace with Rhizopus oryzae under Solid-State Conditions: Changes in the Chemical Composition and Profile of Phenolic Compounds. Microorganisms 11 (2023). 10.3390/microorganisms11040956

23 Muniz, C. E. S. et al. Solid-state fermentation for single-cell protein enrichment of guava and cashew by-products and inclusion on cereal bars. Biocatalysis and Agricultural Biotechnology 25, 101576 (2020). 10.1016/j.bcab.2020.101576

24 Ong, A. & Lee, C.-L. K. Cooperative metabolism in mixed culture solid-state fermentation. LWT 152, 112300 (2021). 10.1016/j.lwt.2021.112300

25 Wang, Z. et al. Reduction of off-flavor volatile compounds in okara by fermentation with four edible fungi. LWT 155, 112941 (2022). 10.1016/j.lwt.2021.112941

26 Andayani, S. N., Lioe, H. N., Wijaya, C. H. & Ogawa, M. Umami fractions obtained from water-soluble extracts of red oncom and black oncom-Indonesian fermented soybean and peanut products. J Food Sci 85, 657–665 (2020). 10.1111/1750-3841.14942

27 Kamble, D. B. & Rani, S. Bioactive components, in vitro digestibility, microstructure and application of soybean residue (okara): a review. Legume Science 2, e32 (2020). 10.1002/leg3.32

28 Zhao, T., Ying, P., Zhang, Y., Chen, H. & Yang, X. Research Advances in the High-Value Utilization of Peanut Meal Resources and Its Hydrolysates: A Review. Molecules 28 (2023). 10.3390/molecules28196862

29 Surono, I. S. in Ethnic Fermented Foods and Alcoholic Beverages of Asia (ed Jyoti Prakash Tamang) 341–382 (Springer India, 2016).

30 Ho, C. C. Identity and characteristics of Neurospora intermedia responsible for oncom fermentation in Indonesia. Food Microbiology,, 115–132 (1986).

31 Turner, B. C. Two Ecotypes of Neurospora Intermedia. Mycologia 79, 425–432 (1987). 10.1080/00275514.1987.12025400

32 Shurtleff, W. & Aoyagi, A. The book of tempeh. Vol. 1 (Soyinfo Center, 1979).

33 Landis Elizabeth, A., et al. Microbial Diversity and Interaction Specificity in Kombucha Tea Fermentations. mSystems 7, e00157–00122 (2022). 10.1128/msystems.00157-22

34 Menzel, P., Ng, K. L. & Krogh, A. Fast and sensitive taxonomic classification for metagenomics with Kaiju. Nature Communications 7, 11257 (2016). 10.1038/ncomms11257

35 Kamilari, E., Stanton, C., Reen, F. J. & Ross, R. P. Uncovering the Biotechnological Importance of Geotrichum candidum. Foods 12, 1124 (2023).

36 Hesseltine, C. W. in Frontiers in Industrial Mycology (ed Gary F. Leatham) 24–39 (Springer US, 1992).

37 Ghasemi, R. et al. Meyerozyma guilliermondii species complex: review of current epidemiology, antifungal resistance, and mechanisms. Braz J Microbiol 53, 1761–1779 (2022). 10.1007/s42770-022-00813-2

38 Chakrabarti, A. et al. Apophysomyces elegans: epidemiology, amplified fragment length polymorphism typing, and in vitro antifungal susceptibility pattern. J Clin Microbiol 48, 4580–4585 (2010). 10.1128/jcm.01420-10

39 Lackner, G., Moebius, N., Partida-Martinez, L. P., Boland, S. & Hertweck, C. Evolution of an endofungal Lifestyle: Deductions from the Burkholderia rhizoxinica Genome. BMC Genomics 12, 210 (2011). 10.1186/1471-2164-12-210

40 Yang, S., Anikst, V. & Adamson, P. C. Endofungal Mycetohabitans rhizoxinica Bacteremia Associated with Rhizopus microsporus Respiratory Tract Infection. Emerg Infect Dis 28, 2091–2095 (2022). 10.3201/eid2810.220507

41 Mladenović, K. G. et al. Enterobacteriaceae in food safety with an emphasis on raw milk and meat. Appl Microbiol Biotechnol 105, 8615–8627 (2021). 10.1007/s00253-021-11655-7

42 Rodrigues, C. et al. High Prevalence of Klebsiella pneumoniae in European Food Products: a Multicentric Study Comparing Culture and Molecular Detection Methods. Microbiology Spectrum 10, e02376–02321 (2022). 10.1128/spectrum.02376-21

43 Xie, M. et al. An integrated metagenomic/metaproteomic investigation of microbiota in dajiang-meju, a traditional fermented soybean product in Northeast China. Food Research International 115, 414–424 (2019). 10.1016/j.foodres.2018.10.076

44 Tamang, J. P. et al. Shotgun metagenomics of Cheonggukjang, a fermented soybean food of Korea: Community structure, predictive functionalities and amino acids profile. Food Research International 151, 110904 (2022). 10.1016/j.foodres.2021.110904

45 Caffrey, E. B., Olm, M. R., Kothe, C. I., Evans, J. & Sonnenburg, J. L. MiFoDB, a workflow for microbial food metagenomic characterization, enables high-resolution analysis of fermented food microbial dynamics. bioRxiv, 2024.2003.2029.587370 (2024). 10.1101/2024.03.29.587370

46 Chen, C. et al. Metagenomic and metaproteomic analyses of microbial amino acid metabolism during Cantonese soy sauce fermentation. Front Nutr 10, 1271648 (2023). 10.3389/fnut.2023.1271648

47 He, W. & Chung, H. Y. Exploring core functional microbiota related with flavor compounds involved in the fermentation of a natural fermented plain sufu (Chinese fermented soybean curd). Food Microbiol 90, 103408 (2020). 10.1016/j.fm.2019.103408

48 Svedberg, J. et al. Convergent evolution of complex genomic rearrangements in two fungal meiotic drive elements. Nat Commun 9, 4242 (2018). 10.1038/s41467-018-06562-x

49 Perkins, D. D. & Davis, R. H. Evidence for safety of Neurospora species for academic and commercial uses. Appl Environ Microbiol 66, 5107–5109 (2000). 10.1128/aem.66.12.5107-5109.2000

50 Niquet, C. & Tessier, F. J. Free glutamine as a major precursor of brown products and fluorophores in Maillard reaction systems. Amino Acids 33, 165–171 (2007). 10.1007/s00726-006-0388-9

51 Ninomiya, K. Science of umami taste: adaptation to gastronomic culture. Flavour 4, 13 (2015). 10.1186/2044-7248-4-13

52 Mellor, N., Bligh, F., Chandler, I. & Hodgman, C. Reduction of off-flavor generation in soybean homogenates: a mathematical model. J Food Sci 75, R131–138 (2010). 10.1111/j.1750-3841.2010.01733.x

53 Beelman, R. B. et al. Health consequences of improving the content of ergothioneine in the food supply. FEBS Lett 596, 1231–1240 (2022). 10.1002/1873-3468.14268

54 Halliwell, B. & Cheah, I. Ergothioneine, where are we now? FEBS Lett 596, 1227–1230 (2022). 10.1002/1873-3468.14350

55 Maini Rekdal, V., et al. Edible mycelium bioengineered for enhanced nutritional value and sensory appeal using a modular synthetic biology toolkit. Nature Communications 15, 2099 (2024). 10.1038/s41467-024-46314-8

56 Bello, M. H., Barrera-Perez, V., Morin, D. & Epstein, L. The Neurospora crassa mutant NcDeltaEgt-1 identifies an ergothioneine biosynthetic gene and demonstrates that ergothioneine enhances conidial survival and protects against peroxide toxicity during conidial germination. Fungal Genet Biol 49, 160–172 (2012). 10.1016/j.fgb.2011.12.007

57 Yamaguchi, F., Ota, Y. & Hatanaka, C. Extraction and purification of pectic polysaccharides from soybean okara and enzymatic analysis of their structures. Carbohydrate Polymers 30, 265–273 (1996). 10.1016/S0144-8617(96)00046-X

58 Li, B., Lu, F., Nan, H. & Liu, Y. Isolation and structural characterisation of okara polysaccharides. Molecules 17, 753–761 (2012). 10.3390/molecules17010753

59 Wu, V. W. et al. The regulatory and transcriptional landscape associated with carbon utilization in a filamentous fungus. Proc Natl Acad Sci U S A 117, 6003–6013 (2020). 10.1073/pnas.1915611117

60 Kumar, V., Rani, A. & Hussain, L. Investigations of Amino Acids Profile, Fatty Acids Composition, Isoflavones Content and Antioxidative Properties in Soy Okara. Asian Journal of Chemistry 28, 903–906 (2016). 10.14233/ajchem.2016.19548

61 Benz, J. P. et al. A comparative systems analysis of polysaccharide-elicited responses in Neurospora crassa reveals carbon source-specific cellular adaptations. Mol Microbiol 91, 275–299 (2014). 10.1111/mmi.12459

62 Gibbons, John G. et al. The Evolutionary Imprint of Domestication on Genome Variation and Function of the Filamentous Fungus Aspergillus oryzae. Current Biology 22, 1403–1409 (2012). 10.1016/j.cub.2012.05.033

63 Ropars, J. et al. Domestication of the Emblematic White Cheese-Making Fungus Penicillium camemberti and Its Diversification into Two Varieties. Current Biology 30, 4441–4453.e4444 (2020). 10.1016/j.cub.2020.08.082

64 Varela, J. A. et al. Origin of Lactose Fermentation in Kluyveromyces lactis by Interspecies Transfer of a Neo-functionalized Gene Cluster during Domestication. Current Biology 29, 4284–4290.e4282 (2019). 10.1016/j.cub.2019.10.044

65 Sierra-Patev, S. et al. A global phylogenomic analysis of the shiitake genus Lentinula. Proc Natl Acad Sci U S A 120, e2214076120 (2023). 10.1073/pnas.2214076120

66 Svedberg, J. et al. Convergent evolution of complex genomic rearrangements in two fungal meiotic drive elements. Nature Communications 9, 4242 (2018). 10.1038/s41467-018-06562-x

67 Yan, S., Xu, Y. & Yu, X.-W. From induction to secretion: a complicated route for cellulase production in Trichoderma reesei. Bioresources and Bioprocessing 8, 107 (2021). 10.1186/s40643-021-00461-8

68 Sineiro, J., Dominguez, H., Núñez, M. J. & Lema, J. M. Hydrolysis of microcrystalline cellulose by cellulolytic complex of Trichoderma reesei in low-moisture media. Enzyme and Microbial Technology 17, 809–815 (1995). 10.1016/0141-0229(94)00107-3

69 Sonoda, M. T. et al. Structure and dynamics of Trichoderma harzianum Cel7B suggest molecular architecture adaptations required for a wide spectrum of activities on plant cell wall polysaccharides. Biochimica et Biophysica Acta (BBA) - General Subjects 1863, 1015–1026 (2019). 10.1016/j.bbagen.2019.03.013

70 Beuchat, L. R. Fungal Fermentation of Peanut Press Cake. Econ Bot 30, 227–234 (1976). Doi 10.1007/Bf02909731

71 Blumenthal, C. Z. Production of toxic metabolites in Aspergillus niger, Aspergillus oryzae, and Trichoderma reesei: justification of mycotoxin testing in food grade enzyme preparations derived from the three fungi. Regul Toxicol Pharmacol 39, 214–228 (2004). 10.1016/j.yrtph.2003.09.002

72 Dubey, M. K. et al. PR Toxin - Biosynthesis, Genetic Regulation, Toxicological Potential, Prevention and Control Measures: Overview and Challenges. Front Pharmacol 9, 288 (2018). 10.3389/fphar.2018.00288

73 Trinci, A. P. J. Myco-protein: A twenty-year overnight success story. Mycological Research 96, 1–13 (1992). 10.1016/S0953-7562(09)80989-1

74 Bartholomai, B. M. et al. Safety evaluation of Neurospora crassa mycoprotein for use as a novel meat alternative and enhancer. Food Chem Toxicol 168, 113342 (2022). 10.1016/j.fct.2022.113342

75 Jaeger, S. R. et al. Check-all-that-apply (CATA) questions for sensory product characterization by consumers: Investigations into the number of terms used in CATA questions. Food Quality and Preference 42, 154–164 (2015). 10.1016/j.foodqual.2015.02.003

76 Rahmawati, D., Astawan, M., Putri, S. P. & Fukusaki, E. Gas chromatography-mass spectrometry-based metabolite profiling and sensory profile of Indonesian fermented food (tempe) from various legumes. Journal of Bioscience and Bioengineering 132, 487–495 (2021). 10.1016/j.jbiosc.2021.07.001

77 Genta, H. D., Genta, M. L., Álvarez, N. V. & Santana, M. S. Production and acceptance of a soy candy. Journal of Food Engineering 53, 199–202 (2002). 10.1016/S0260-8774(01)00157-1

78 Xie, M., Huff, H., Hsieh, F. & Mustapha, A. Puffing of okara/rice blends using a rice cake machine. J Food Sci 73, E341–348 (2008). 10.1111/j.1750-3841.2008.00905.x

79 Beadle, G. W. & Tatum, E. L. Genetic Control of Biochemical Reactions in Neurospora. Proc Natl Acad Sci U S A 27, 499–506 (1941). 10.1073/pnas.27.11.499

80 McCluskey, K., Wiest, A. & Plamann, M. The Fungal Genetics Stock Center: a repository for 50 years of fungal genetics research. J Biosci 35, 119–126 (2010). 10.1007/s12038-010-0014-6

81 Watarai, N., Yamamoto, N., Sawada, K. & Yamada, T. Evolution of Aspergillus oryzae before and after domestication inferred by large-scale comparative genomic analysis. DNA Res 26, 465–472 (2019). 10.1093/dnares/dsz024

82 Dumas, E. et al. Independent domestication events in the blue-cheese fungus Penicillium roqueforti. Mol Ecol 29, 2639–2660 (2020). 10.1111/mec.15359

83 Wang, Y., Li, H., Feng, G., Du, L. & Zeng, D. Biodegradation of diuron by an endophytic fungus Neurospora intermedia DP8-1 isolated from sugarcane and its potential for remediating diuron-contaminated soils. PLoS One 12, e0182556 (2017). 10.1371/journal.pone.0182556

84 Kuo, H.-C. et al. Secret lifestyles of Neurospora crassa. Scientific Reports 4, 5135 (2014). 10.1038/srep05135

85 Kjærbølling, I. et al. A comparative genomics study of 23 Aspergillus species from section Flavi. Nature Communications 11, 1106 (2020). 10.1038/s41467-019-14051-y

86 Tannenbaum, S. R., Wang, D. I. C. Single-Cell Protein II. (MIT press, 1975).

87 Maini Rekdal, V., et al. From lab to table: Expanding gastronomic possibilities with fermentation using the edible fungus Neurospora intermedia. International Journal of Gastronomy and Food Science 34, 100826 (2023). 10.1016/j.ijgfs.2023.100826

88 Gilbert, C. et al. Living materials with programmable functionalities grown from engineered microbial co-cultures. Nature Materials 20, 691–700 (2021). 10.1038/s41563-020-00857-5

89 Yamashita, H. Koji Starter and Koji World in Japan. J Fungi (Basel) 7 (2021). 10.3390/jof7070569

