## Supplemental information for "Multi-omics analysis of a traditional fermented food reveals a byproduct-associated subpopulation of *Neurospora intermedia* for waste-to-food upcycling"

Vayu Maini Rekdal et al.

**Table S1. Physicochemical and geographical data for oncom samples collected from traditional producers in Java, Indonesia.** Samples were collected from regional producers throughout Western Java. The site where the sample was collected is indicated in the table. pH was measured using a pH meter, and hardness was measured using a penetrometer. Compactness was determined visually (see methods for details). Results are mean and SD of two replicates for pH, five replicates for hardness. + (poor mold growth, non-compact texture); ++ (decent growth of mold, texture is quite compact and dense); +++ (good mold growth, compact and dense texture); ++++ (very good mold growth, very compact and dense texture). See methods for description of compactness measurements.

| **Sample number** | **Oncom type** | **Production site** | **Hardness (mm/5 sec)** | **compactness of texture** | **pH** |
| --- | --- | --- | --- | --- | --- |
| 1 | Red | Leuweung Kolot | 22.42 ± 3.08 | ++ | 5.50 ± 0.30 |
| 2 | Black | Cibeureum | 7.78 ± 0.14 | ++++ | 4.71 ± 0.01 |
| 3 | Red | Cibalagung | 15.69 ± 0.69 | ++++ | 5.41 ± 0.25 |
| 4 | Red | Leuwiliang | 14.93 ± 0.72 | ++++ | 5.84 ± 0.44 |
| 5 | Black | Cijujung | 13.85 ± 0.58 | + | 6.18 ± 0.46 |
| 6 | Black | Bogor (producer A) | 11.88 ± 1.30 | ++ | 6.73 ± 0.08 |
| 7 | Black | Bogor (producer B) | 11.12 ± 1.45 | + | 6.50 ± 0.09 |
| 8 | Black | Ciawi | 14.04 ± 3.91 | + | 7.08 ± 0.02 |
| 9 | Black | Ciampea | 7.58 ± 3.37 | +++ | 7.58 ± 0.42 |
| 10 | Red | Depok | 13.92 ± 1.15 | ++++ | 5.83 ± 0.07 |
| 11 | Red | Cibinong | 14.72 ± 1.08 | ++++ | 5.94 ± 0.01 |
| 12 | Red | Jakarta Selatan | 15.12 ± 2.13 | +++ | 5.79 ± 0.02 |
| 13 | Red | Girimulya (producer A) | 19.22 ± 2.50 | +++ | 6.63 ± 0.07 |
| 14 | Red | Girimulya (producer C | 17.60 ± 1.98 | ++ | 6.88 ± 0.02 |
| 15 | Red | Girimulya (producer B) | 17.64 ± 2.26 | ++ | 6.55 ± 0.01 |
| 16 | Red | Ciampea | 13.18 ± 2.93 | ++++ | 6.56 ± 0.15 |

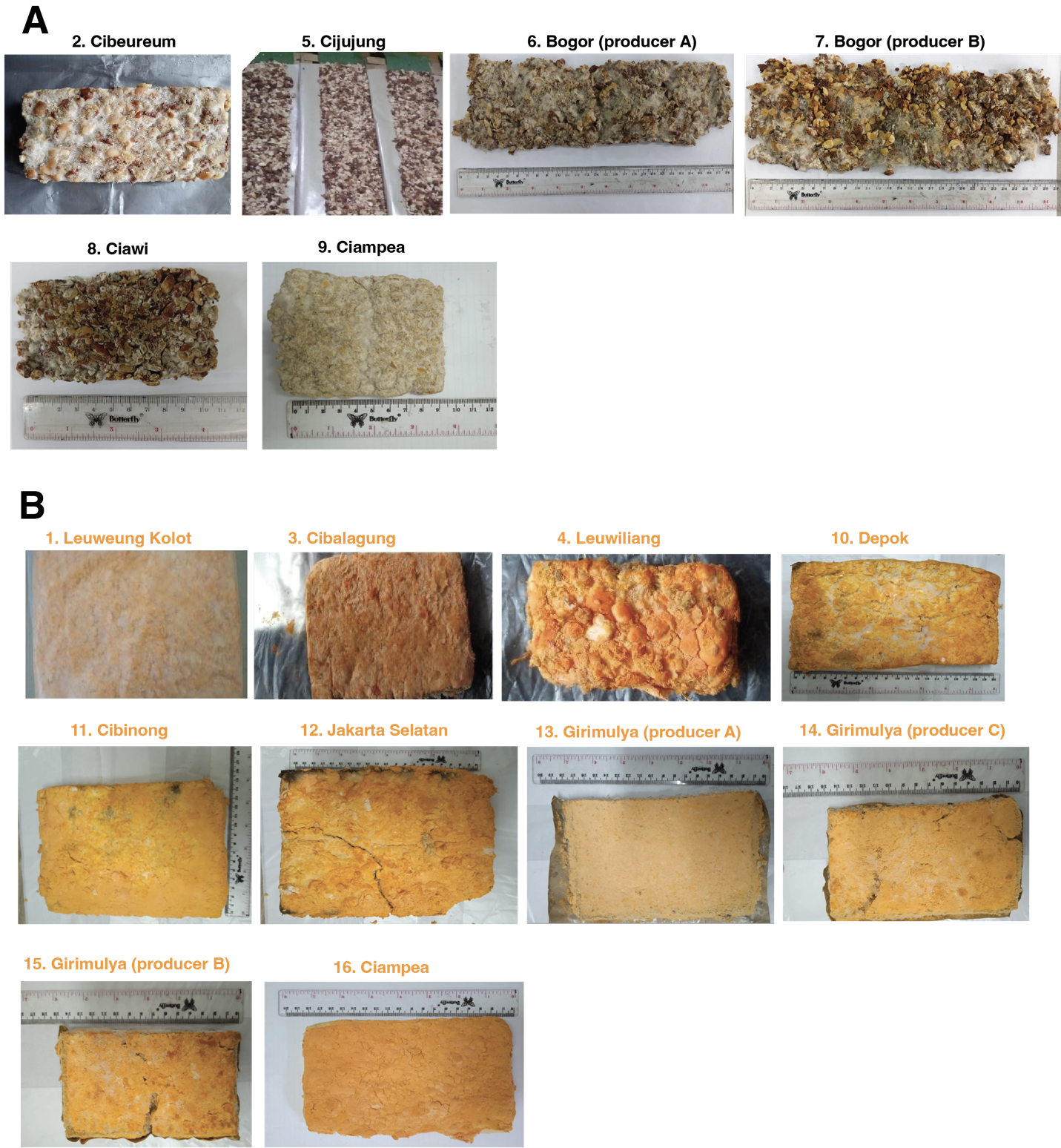

**Figure S1. Photos of black and red oncom samples collected from different producers in Java, Indonesia.** The numbers refer to the locations shown in Table S1 and in Fig. 1 in the main text. All these samples were subjected to metagenome sequencing for taxonomic profiling. Three red and two black oncom samples were also subjected to amplicon sequencing (Fig. S3, S4).

**Table S2. Genome statistics for oncom-derived *Neurospora intermedia* FGSC #2613.** *Neurospora intermedia* FGSC #2613, which was isolated from oncom samples, was subjected to whole genome sequencing using PacBio and was annotated using transcriptomics, generating a high-quality draft genome that is publicly available through the Joint Genome Institute (JGI). The completeness of the genome outperforms the previous best *N. intermedia* genome, which was sequenced in an unrelated study^1^. *N. crassa,* the model ascomycete and the first *Neurospora* to be sequenced, is included for comparison.

|  | ***N. intermedia* #2613** | ***N. intermedia* #8793 (Previous best)** | ***N. crassa* OR74A** |
| --- | --- | --- | --- |
| **Length (bp)** | 39262359 | 39486933 | 41102378 |
| **Contigs** | 21 | 1147 | 21 |
| **N50** | 4,276,046 | 70635 | 6000761 |
| **GC content** | 0.49 | 0.49 | 0.48 |

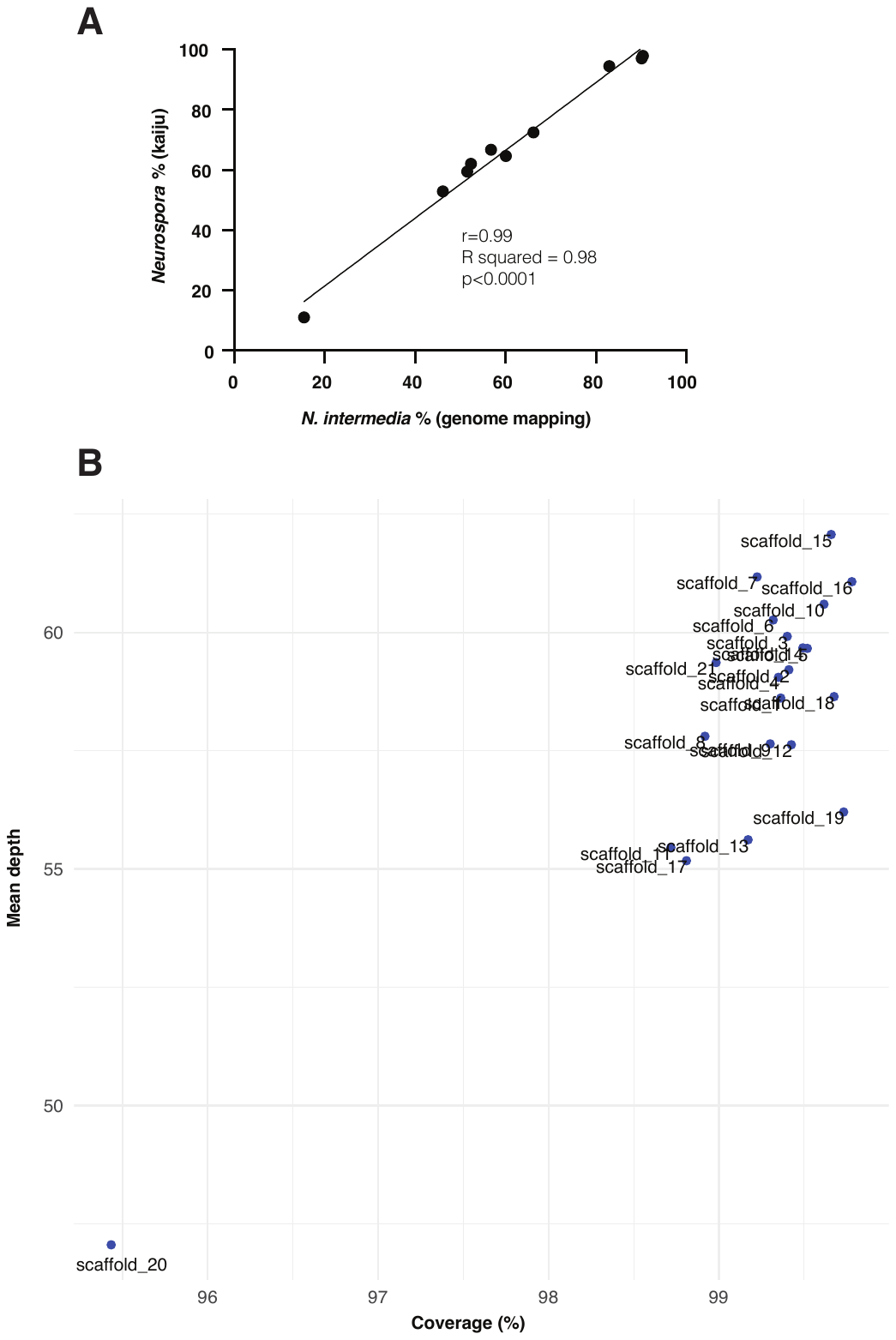

**Figure S2. Mapping of shotgun metagenome data to *N. intermedia* genome**. A) Correlation between % reads assigned to the *Neurospora* genus using the kaiju database, which does not contain a genome for *Neurospora intermedia*, and the % of reads mapped to our sequenced *N. intermedia* FGSC #2613 genome using this as the only query (<2 mismatches allowed). There was a strong correlation (r=0.99, R squared = 0.98, p<0.0001) between the % abundance by these two methods, indicating that a majority of the % reads assigned to *Neurospora* by kaiju also readily map to the sequenced *N. intermedia* FGSC #2613 genome. B) Coverage of the *N. intermedia* FGSC #2613 genome by scaffold. The shotgun metagenome sequences from all samples were combined and mapped against the *N. intermedia* genome. The results indicate >98.5% coverage for all scaffolds but scaffold 20 (mean coverage=99%). This indicates that the oncom samples harbor sequences that align with the *N. intermedia* genome.

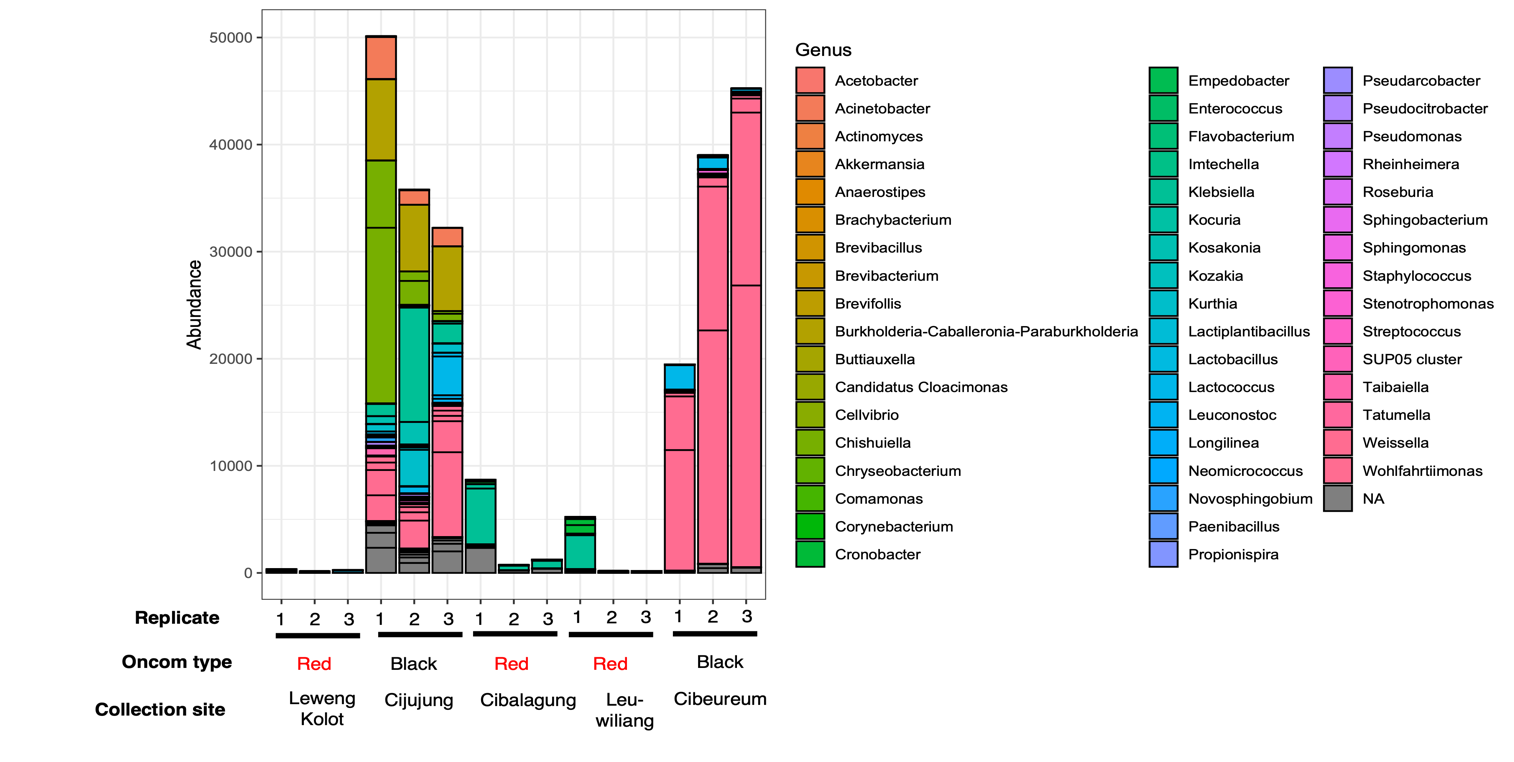
**Figure S3. Bacterial abundance based on 16s amplicon sequencing of a subset of red and black oncom samples collected from traditional producers in Java, Indonesia.** The graph displays the bacterial abundance at the genus level. DNA concentrations were normalized prior to sequencing. Analysis was conducted using phyloseq. Bars indicate the number of reads assigned to a specific genus. Three replicates from each of the five oncom products collected from different producers were sequenced. While the black oncom sample from Cijujung had a diverse microbial community, including *Weisella*, *Brevibacterium*, *Klebsiella*, and *Acetobacter,* the black oncom sample from Cibeureum was mostly dominated by the lactic acid bacterium *Weissella,* which is consistent with the substrates being soaked and subjected to a brief lactic acid fermentation prior to inoculation. In contrast, red oncom samples did not harbor a diverse microbial community. Some red oncom samples harbored enterobacteria such as *Klebsiella.*

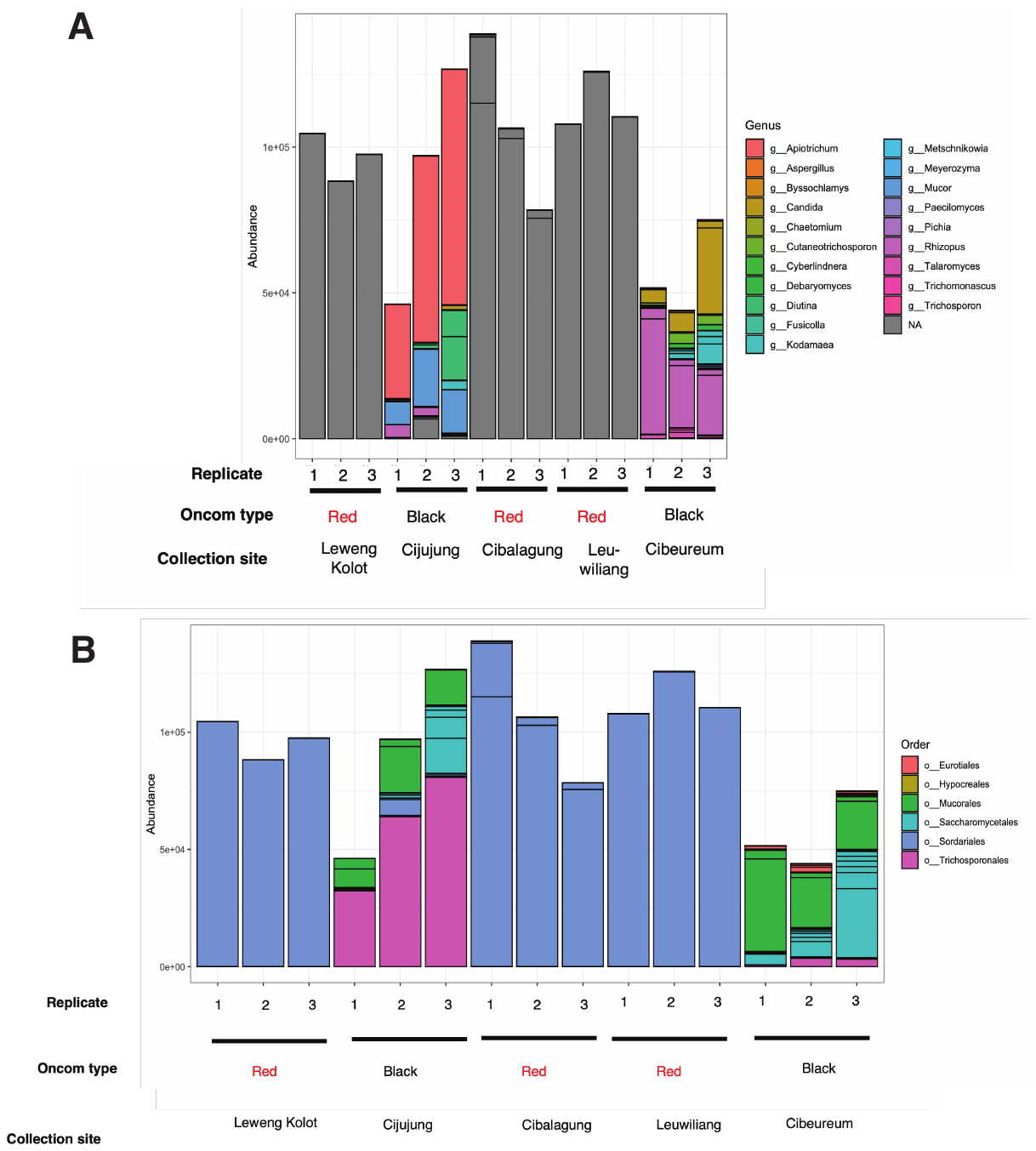

**Figure S4. Fungal abundance based on ITS amplicon sequencing of a subset of red and black oncom samples collected from traditional producers in Java, Indonesia.** The graph displays the fungal abundance at the genus (A) and order (B) levels. DNA concentrations were normalized prior to sequencing. Analysis was conducted using phyloseq. Bars indicate the number of reads assigned to a specific group. Three replicates from each of the five oncom “cakes” collected from different producers were sequenced. A) At the genus level, black and red oncom samples harbored different fungal communities, and while red oncom appeared to be dominated by a single genus, the two black oncom samles harbored different fungal communities. However, the specific genus identity in the red oncom samples could not be assigned using the analysis pipeline and database. Black oncom from Cijujung was dominated by *Apotrichium* but also harbored *Mucor* and *Debaromyces* members. The other black oncom sample was dominated by *Rhizopus*. B) Order-level assignment revealed that the unassigned major fungus at the genus level belonged to the *Sordiarales* order. Comparison of these results with the metagenome sequencing (see Figure 1 in main text) suggests that the major fungus is *Neurospora*. There was also a small number of a fungus belonging to the *Sordiaria* genus detected based on metagenomic sequencing (Figure 1 in main text), which may explain the presence of two distinct *Sordiarales* fungi (based on ITS amplicon sequencing data) at the order level in the Cibalagung sample.

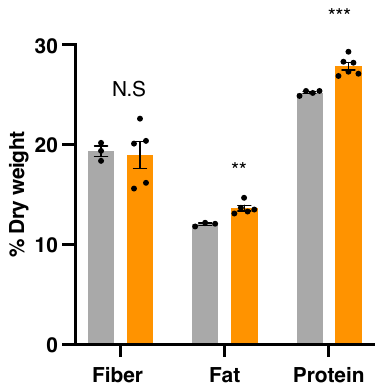

**Figure S4. Nutrient profile of raw and fermented okara.** While the fiber did not change during the fermentation, both protein (***p<0.001, unpaired t-test) and lipid content (**=p<0.01, unpaired t-test) significantly increased, from 25% to 28% and from 12-14%, respectively (dry weight basis). For protein, results are mean and SEM from n=6 biological for fermented and n=4 biological replicates for unfermented samples. For lipids and fiber, results are mean and SEM from n=3 biological for fermented and n=5 biological replicates for unfermented samples.

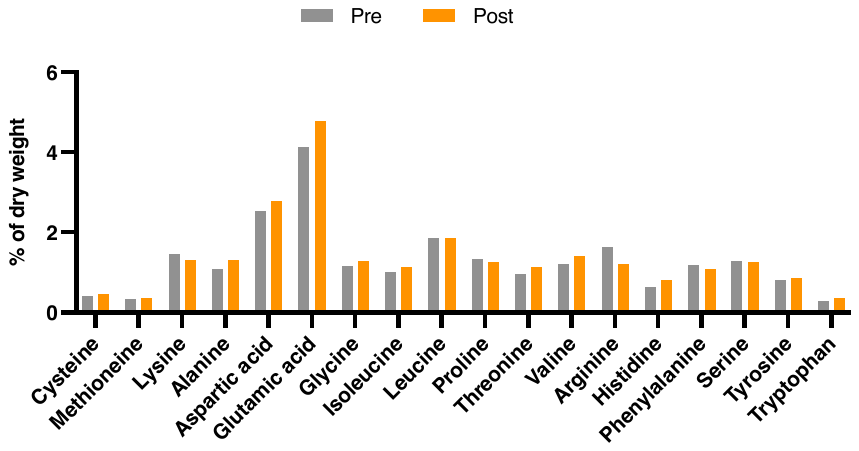

**Figure S5. Amino acid composition of raw and fermented okara.** Okara samples were fermented with *N. intermedia* for 48 hours in solid-state. Lyophilized, hydrolyzed samples were subjected to amino acid profiling. The graph demonstrates that there were no major changes in the global amino acid composition before and after fermentation. However, like raw okara, fermented okara contained all the essential amino acids (histidine, isoleucine, leucine, lysine, methionine, phenylalanine, threonine, tryptophan, and valine). One lyophilized sample of each of okara substrate and fermented okara was sequenced.

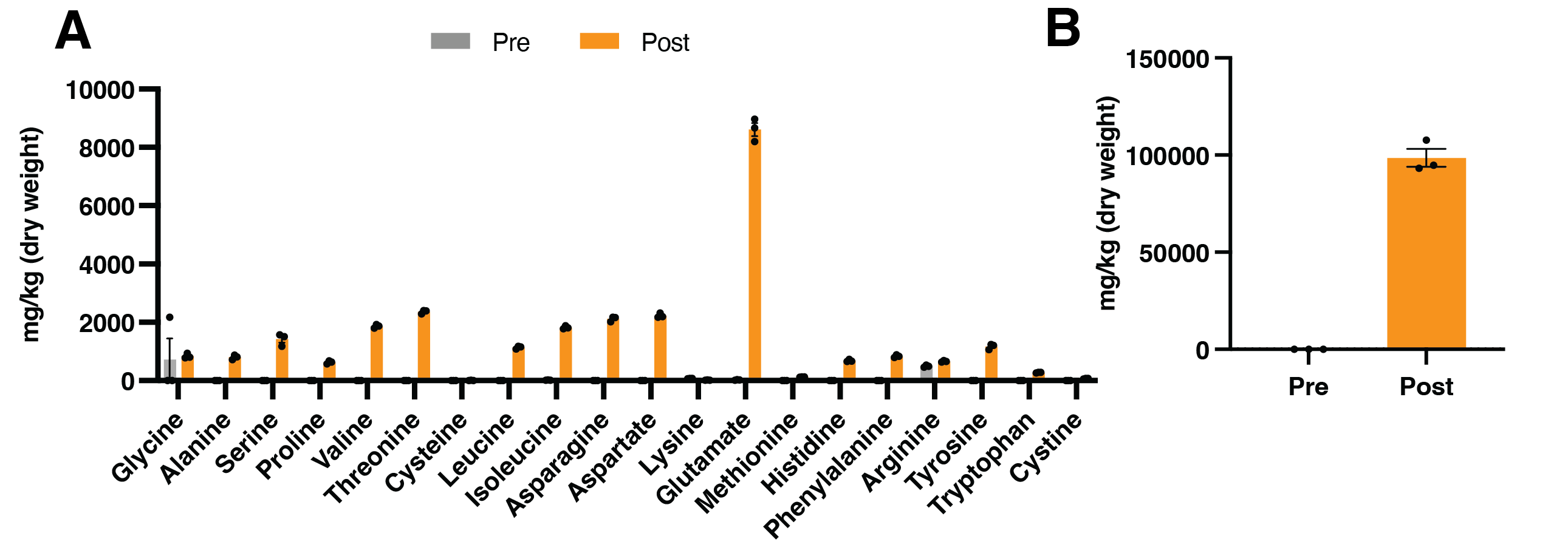

**Figure S6. Free amino acids in raw and fermented okara.** Okara samples were fermented with *N. intermedia* in solid-state. Lyophilized samples were extracted and subjected to amino acid analysis by LC-MS. Results are mean and SEM of three biological replicates and are normalized to the dry weight. A) shows all analyzed amino acids except for glutamine, which was produced at such high levels that it obscured changes in the other amino acids. Aside from glutamine, glutamate (8610 mg/kg) was by far the most produced amino acids. Only arginine was present at similar amounts pre and post fermentation. B) Levels of glutamine pre and post fermentation. This was produced at the highest levels of all amino acids (98571 mg/kg).

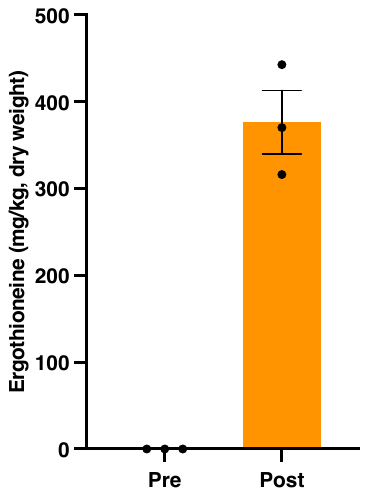

**Figure S7. Ergothioneine levels in raw and fermented okara.** Okara samples were fermented with *N. intermedia* for 48 hours in solid-state. The fermentation produced ergothioneine, a potent antioxidant that is associated with several health benefits. Ergothioneine was absent in the starting material, suggesting active fungal biosynthesis during the fermentation process. Results are mean and SEM on a dry weight basis from n=3 biological replicates and were analyzed by LC-MS.

**Table S3. GC-MS analysis of volatile aroma compounds in raw okara as well as raw and cooked okara fermented with *N. intermedia*.**  Values are expressed in mean and standard deviation (SD). Values are expressed as 2-methyl-3-heptanone equivalent (µg/L). Compounds were identified by comparison of the MS spectra with the NIST library as well as by comparison of RI (Kovat indices). Retention index is based on Thermo TG-5SILMS column using C7-C27 as external references. The p-value is based on ANOVA results and post-hoc test (Tukey HSD) in volatile aroma compounds for each sample. Different letter(s) (a, b, or ab) within the same row (compound) indicates statistically significant differences in quantity between the samples (p-value <0.05). For example, a row where raw oncom, cooked oncom, and okara have the same letter, they are not significantly different. A total of 30 volatile compounds were found in all the samples, including one acid compound, three aldehydes, one ester, two ethers, four ketones, 17 hydrocarbons and one phenol. The most abundant compounds in the okara samples were hexanal, 1,3-bis(1,1-dimethylethyl)-benzene, although this last compound was found in a high concentration in the cooked oncom and at lower concentration in oncom. The flavor of the original okara is described as having grassy, green and beany aromas, all undesirable notes normally found in bean products^2^. These attributes can be traced to hexanal, a product of enzymatic oxidation due to the presence of lipoxygenase during processing of the raw materials^3^. After the fermentation, the abundance was reduced 40-times from 86.86 µg/L in okara to 2.47 µg/L in oncom. Additionally, the acid compound 2-methyl butyric acid was found in okara but disappeared during the fermentation. It is produced during anaerobic fermentation of short-chain fatty acids^4^ and is responsible for cheesy and rancid aromas^5^. In addition, the cooked fermented okara developed hydrocarbon compounds related to the cooking method, such as 2-ethyl-1,4-dimethyl-benzene (the most abundant), dodecane, and others^6^. Moreover, ethylcyclohexane developed during cooking; it is responsible for pineapple and pear aromas, and is directly related to roasted attributes developing via the Maillard reaction^7^.

|  |  |  |  | **Concentration (µg/L)** | | | | | |  |
| --- | --- | --- | --- | --- | --- | --- | --- | --- | --- | --- |
| **Calculated RI^1^** | **Library RI** | **Retention time (min)** | **Putative Compounds** | **Cooked Oncom** |  | **Oncom** |  | **Okara** |  | ***p*-value** |
|  |  |  | **Acid** |  |  |  |  | |  |  |
| 852 | 861 | 5.81 | 2-methyl-butyric acid | 0.124 a | ±  0.05 | 0.002 b | ±0.00 | 0.009 b | ±0.00 | **0.009** |
|  |  |  | **Alcohols** |  |  |  |  | |  |  |
| NA | 739 | 3.10 | 2-methyl-1-butanol | 0.474 b | ±  0.25 | 4.124a | ±1.75 | 0.118 b | ±0.02 | **0.022** |
| 887 | 872 | 6.61 | 1,3-dimethylcyclopentanol | 4.240 b | ±0.62 | 12.01a | ±1.64 | 1.502 b | ±0.57 | **0.000** |
| 982 | 981 | 9.25 | vinyl amyl carbinol | 7.373 a | ±0.81 | 2.233 b | ±0.25 | 3.288 b | ±0.54 | **0.000** |
|  |  |  | **Aldehydes** |  |  |  |  | |  |  |
| NA | 652 | 2.07 | 3-methyl-butanal | 0.127 a | ±0.07 | 0.003 a | ±0.00 | 0.001 a | ±0.00 | 0.042 |
| 780 | 800 | 4.30 | hexanal | 16.373 ab | ±3.60 | 2.468 b | ±1.77 | 86.857a | ±57.59 | **0.044** |
|  |  |  | **Esters** |  |  |  |  | |  |  |
| 1596 | 1612 | 26.42 | methyl n-tridecanoate | 0.000 b | ±0.00 | 0.036 a | ±0.00 | 0.000 b | ±0.00 | **<0,0001** |
|  |  |  | **Ethers** |  |  |  |  | |  |  |
| NA | 703 | 2.53 | 2-ethyl-furan | 0.352 a | ±0.13 | 0.210 a | ±0.09 | 1.402 a | ±1.48 | 0.260 |
| 712 | 742 | 3.34 | 2,4,5-trimethyl-1,3-dioxolane | 3.057 a | ±0.77 | 2.032 a | ±0.43 | 0.063 b | ±0.01 | **0.005** |
|  |  |  | **Ketones** |  |  |  |  | |  |  |
| 961 | 1175 | 8.65 | 1-phenyl-1,2-propanedione | 27.509 a | ±6.67 | 4.666 b | ±2.35 | 0.489 b | ±0.48 | **0.001** |
| 1262 | 1249 | 17.21 | 1,3-bis(1,1-dimethylethyl)-benzene | 472.537 a | ±29.29 | 34.416 b | ±12.26 | 60.102 b | ±28.41 | **<0,0001** |
| 1464 | 1445 | 23.27 | 1b,5,5,6a-Tetramethyl-octahydro-1-oxa-cyclopropa[a]inden-6-one | 0.182 a | ±0.04 | 0.018 b | ±0.01 | 0.007 b | ±0.00 | **0.000** |
| NA | 555 | 1.69 | Butanone | 1.826 a | ±0.32 | 3.114 a | ±0.77 | 0.175 b | ±0.02 | **0.005** |
|  |  |  | **Miscellaneous and hydrocarbons** |  |  |  |  | |  |  |
| 714 | 708 | 3.38 | 2-chloro-pentane | 0.023 a | ±0.00 | 0.012 a | ±0.00 | 0.014 a | ±0.01 | 0.088 |
| 821 | 813 | 5.07 | Ethylcyclohexane | 0.919 a | ±0.39 | 0.126 b | ±0.02 | 0.276 ab | ±0.01 | **0.029** |
| 863 | 850 | 6.05 | 2,6-dimethyl-2-heptene | 0.099 a | ±0.05 | 3.040 a | ±3.04 | 5.236 a | ±1.02 | 0.124 |
| 1126 | 1133 | 13.37 | 1,2,4,5-tetramethylbenzene | 4.387 a | ±0.24 | 0.411 b | ±0.09 | 0.251 b | ±0.15 | **<0,0001** |
| 1129 | 1119 | 13.45 | 2-ethyl-1,4-dimethyl-benzene | 19.180 a | ±1.05 | 1.787 b | ±0.56 | 0.648 b | ±0.40 | **<0,0001** |
| 1163 | 1145 | 14.29 | 2,3-dihydro-4-methyl-1H-Indene | 0.422 a | ±0.04 | 0.083 b | ±0.02 | 0.051 b | ±0.04 | **<0,0001** |
| 1166 | 1145 | 14.37 | 1,2,3,4-tetramethyl-benzene | 9.329 a | ±0.79 | 1.171 b | ±0.31 | 0.725 b | ±0.46 | **<0,0001** |
| 1206 | 1189 | 15.38 | Naphthalene | 2.356 a | ±0.40 | 0.898 b | ±0.26 | 0.394 b | ±0.24 | **0.002** |
| 1221 | 1200 | 15.88 | Dodecane | 2.817 a | ±0.02 | 0.057 b | ±0.02 | 0.052 b | ±0.03 | **<0,0001** |
| 1285 | 1300 | 17.96 | Tridecane | 1.510 a | ±0.34 | 0.196 b | ±0.07 | 0.094 b | ±0.04 | **0.001** |
| 1318 | 1385 | 18.98 | (E)-3-Tetradecene | 1.156 a | ±0.65 | 0.612 a | ±0.25 | 0.238 a | ±0.03 | 0.153 |
| 1326 | 1409 | 19.24 | 1,1,3-trimethyl-2-(3-methylpentyl)-cyclohexane | 0.864 a | ±0.44 | 0.396 a | ±0.04 | 0.206 a | ±0.02 | 0.096 |
| 1331 | 1318 | 19.38 | 2,3,5,8-tetramethyl-decane | 0.747 a | ±0.25 | 0.287 ab | ±0.12 | 0.111 b | ±0.01 | **0.018** |
| 1419 | 1400 | 22.00 | tetradecane | 1.430 a | ±0.22 | 0.238 b | ±0.07 | 0.096 b | ±0.03 | **0.000** |
| 1511 | 1505 | 24.54 | 1R,3Z,9s-4,11,11-Trimethyl-8-methylenebicycloundec-3-ene | 0.006 ab | ±0.00 | 0.009 a | ±0.00 | 0.001 b | ±0.00 | **0.036** |
| 1964 | 1939 | 32.53 | 1,3,6,10-Cyclotetradecatetraene, 3,7,11-trimethyl-14-(1-methylethyl)-, [S-(E,Z,E,E)]- | 0.203 a | ±0.04 | 0.001 b | ±0.00 | 0.000 b | ±0.00 | **0.000** |
|  |  |  | **Phenols** |  |  |  |  | |  |  |
| 1518 | 1519 | 24.69 | 2,4-Di-tert-butylphenol | 0.040 a | ±0.01 | 0.069 a | ±0.01 | 0.059 a | ±0.03 | 0.365 |

**Table S4. Genes induced by *N. intermedia* in response to okara.** Genes induced above a cutoff of log2foldchange>4 are shown and a statistical cutoff of FDR<0.05. Results are from three biological replicates with okara as the sole carbon source in (1% w/v), compared to a no carbon control, in liquid cultures. CAZyme, BLASTgo, and degradPlantBio annotations are displayed for functional predictions. degradPlantBio is based on homologous genes involved in plant biomass degradation in the related fungus and model ascomycete *Neurospora crassa.* Okara induced a number of predicted CAZymes. Full results for okara and all other carbon sources (n=12) can be found in Supplementary file S1. The JGI protein ID comes from the genome assembly and annotation of Neurospora intermedia FGSC #2613, which is available through the MycoCosm portal at <https://mycocosm.jgi.doe.gov/Neuin1>. The RNA sequencing data across all carbon sources (log2foldchange>1, FDR<0.05) can be found in Supplementary Data S1.

| **JGI protein ID** | **log2foldchange** | **CAZyme annotation** | **goName** | **degraPlantBio** |
| --- | --- | --- | --- | --- |
| 526964 | 11.1785107 | #N/A | response to stress | #N/A |
| 38386 | 8.801304002 | Glycoside Hydrolase Family 25 protein | lysozyme activity | #N/A |
| 533462 | 7.462822093 | #N/A | #N/A | #N/A |
| 546421 | 6.4642685 | #N/A | #N/A | #N/A |
| 524191 | 6.260404518 | #N/A | #N/A | #N/A |
| 524189 | 6.052509787 | #N/A | 4-hydroxyphenylpyruvate dioxygenase activity | #N/A |
| 549776 | 5.28967937 | Carbohydrate-Binding Module Family 1 /  Glycoside Hydrolase Family 5 protein | hydrolase activity, hydrolyzing O-glycosyl compounds | probable cellulase precursor |
| 576181 | 5.259496838 | #N/A | acyl-CoA dehydrogenase activity | #N/A |
| 299215 | 5.187802849 | #N/A | homogentisate 1,2-dioxygenase activity | #N/A |
| 357033 | 5.117855807 | Polysaccharide Lyase Family 3 protein | extracellular region | probable pectate lyase |
| 506499 | 5.084668567 | Carbohydrate-Binding Module Family 1 /  Glycoside Hydrolase Family 6 protein | hydrolase activity, hydrolyzing O-glycosyl compounds | probable cellulose  1,4-beta-cellobiosidase II |
| 527281 | 5.066336691 | #N/A | catalytic activity | #N/A |
| 480919 | 5.062161083 | Glycoside Hydrolase Family 7 protein | hydrolase activity, hydrolyzing O-glycosyl compounds | probable endo-1,4-beta-glucanase  (cellulase) |
| 528065 | 4.965505081 | #N/A | catalytic activity | #N/A |
| 529794 | 4.867243277 | #N/A | dihydrolipoamide branched chain acyltransferase activity | #N/A |
| 356738 | 4.741348328 | #N/A | #N/A | #N/A |
| 570526 | 4.738908222 | #N/A | nucleic acid binding | #N/A |
| 490212 | 4.619127297 | #N/A | catalytic activity | #N/A |
| 538463 | 4.585136467 | #N/A | methylcrotonoyl-CoA carboxylase activity | #N/A |
| 173393 | 4.552213915 | #N/A | ornithine-oxo-acid transaminase activity | #N/A |
| 164864 | 4.545190813 | #N/A | #N/A | #N/A |
| 130505 | 4.51423994 | #N/A | electron transport | #N/A |
| 442209 | 4.448756486 | Glycoside Hydrolase Family 5 protein | hydrolase activity, hydrolyzing O-glycosyl compounds | #N/A |
| 516515 | 4.427878462 | #N/A | #N/A | #N/A |
| 53068 | 4.345697664 | Lytic polysaccharide monooxygenase (formerly GH61) /  Carbohydrate-Binding Module Family 1 | hydrolase activity, hydrolyzing O-glycosyl compounds | related to endoglucanase B  (cellulase) |
| 520753 | 4.250252935 | #N/A | ribose-5-phosphate isomerase activity | #N/A |
| 3776 | 4.237032617 | #N/A | 1-pyrroline-5-carboxylate dehydrogenase activity | #N/A |
| 516864 | 4.182198374 | Glycoside Hydrolase Family 1 protein | hydrolase activity, hydrolyzing O-glycosyl compounds | probable beta-glucosidase |
| 479714 | 4.162143113 | #N/A | monooxygenase activity | #N/A |
| 594518 | 4.15113167 | #N/A | #N/A | #N/A |
| 432986 | 4.034999715 | Glycoside Hydrolase Family 35 protein | beta-galactosidase activity | #N/A |
| 530901 | 4.032914093 | Carbohydrate-Binding Module Family 1 /  Carbohydrate Esterase Family 16 protein | hydrolase activity, hydrolyzing O-glycosyl compounds | #N/A |

**Figure S8. Global co-expression analysis reveals a shared transcriptional module between okara and avicel as well as genes uniquely expressed on okara.**  A matrix of gene expression of 11280 genes predicted from the genome was used to construct a co-expression network using the WGCNA software integrating the full transcriptomic data across all carbon sources. The scale-free network was built with the blockwiseModules function, with a power = 8. To define the modules of the network, the minimum size was established at 30 genes per module and the threshold for merging similar modules was established at 0.15. Furthermore, the weighted adjacency matrix was transformed into a topological overlap measurement matrix (TOM) to estimate connectivity in the network. The network was visualized using the Cytoscape v.3.9.1 program. Network metrics were obtained by Cytoscape v.3.9.1. To determine enriched metabolic pathways in the modules we used the KEGG tool (<https://www.genome.jp/kegg/>). A) shows the global co-expression network, which revealed a shared transcriptional module between okara and avicel, as well as a transcriptional module that was unique to okara. In the network, the values of log2 Fold-change are observed in the comparison okara vs no carbon (FDR<0.05). B) Among other pathways, the unique okara cluster harbored predicted genes involved in the degradation of the amino acids tyrosine (also phenylalanine) as well as leucine. The predicted metabolic pathways are shown. Red indicates that the gene was induced in response to okara. The adjacent table indicates the gene ID, and the log2fold change across okara, avicel, raffinose, galactose, and soy polyssacharides (from left to right). The color corresponds to the level of fold change. The full gene list for each of the two clusters can be found in Supplementary data S1.

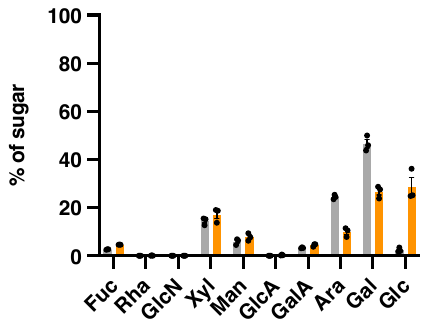

**Figure S9. Sugar composition in the hydrolyzed pectin fraction of raw and fermented okara.** Okara samples were fermented with *N. intermedia* for 48 hours in solid-state. Lyophilized samples were extracted and subjected to sugar analysis by HPAEC after processing and hydrolysis to obtain the pectin-containing fraction. Results are mean and SEM of three biological replicates and are expressed as the mol % of all sugars analyzed. Fuc = fucose, Rha = rhamnose, GlcN = N-acetylglucosamine, Xyl = xylose, Man = mannose, GlcA = glucuronic acid, GalA = galacturonic acid, Ara = arabinose, Gal = galactose, Glc = glucose. There was a decrease in both arabinose and galactose as a result of the fermentation. Other sugars did not decrease significantly. The increase in glucose is likely attributed to glucans, which make up the majority of the fungal cell wall and can be found in the pectin fraction after fermentation. Arabinose and galactose are abundant in rhamnogalacturonan, the main pectic fiber in soy, and the position of these sugars within the side chain of the polymer makes them accessible for hydrolytic release and microbial degradation, which is consistent with the data. There was not a significant decrease of galacturonic acid, which is found in more inaccessible portions of the pectin polymer. Grey = okara substrate; Orange=post fermentation.

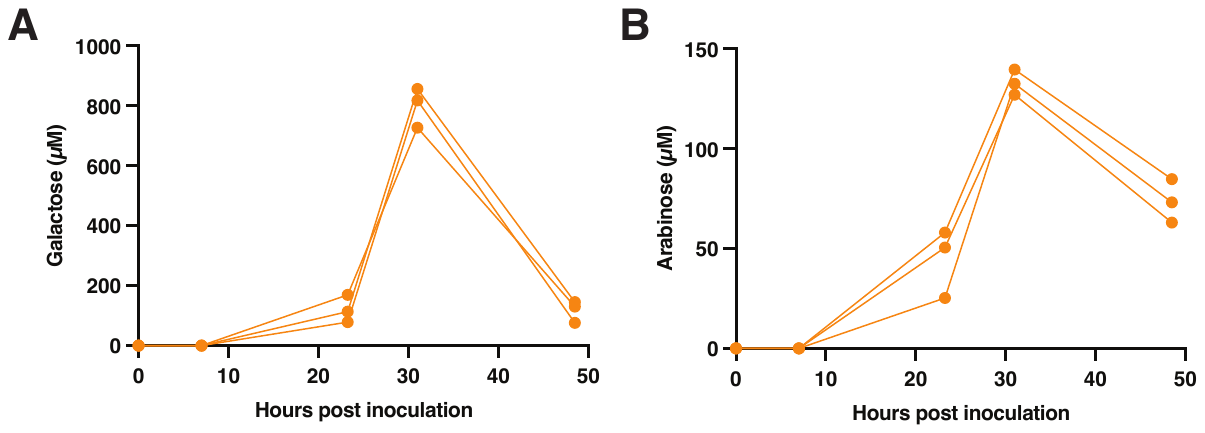

**Figure S10. Profiling of arabinose and galactose during *N. intermedia* growth on okara as the sole carbon source in liquid cultures.** Results display three biological replicates. Supernatants of VMM medium liquid cultures harboring okara as the sole carbon source (1 % w/v) were analyzed for secreted galactose (A) and arabinose (B) by HPAEC. Analysis of okara SSF had suggested both of these sugars are released during the fermentation (Figure 2 in main text) and are also depleted from the pectin fraction, suggesting active release from pectin and subsequent fungal consumption. The graph above supports this conclusion as it reveals initial release of these sugars from the okara substrate, followed by disappearance from the culture medium, suggesting active fungal hydrolysis and subsequent uptake.

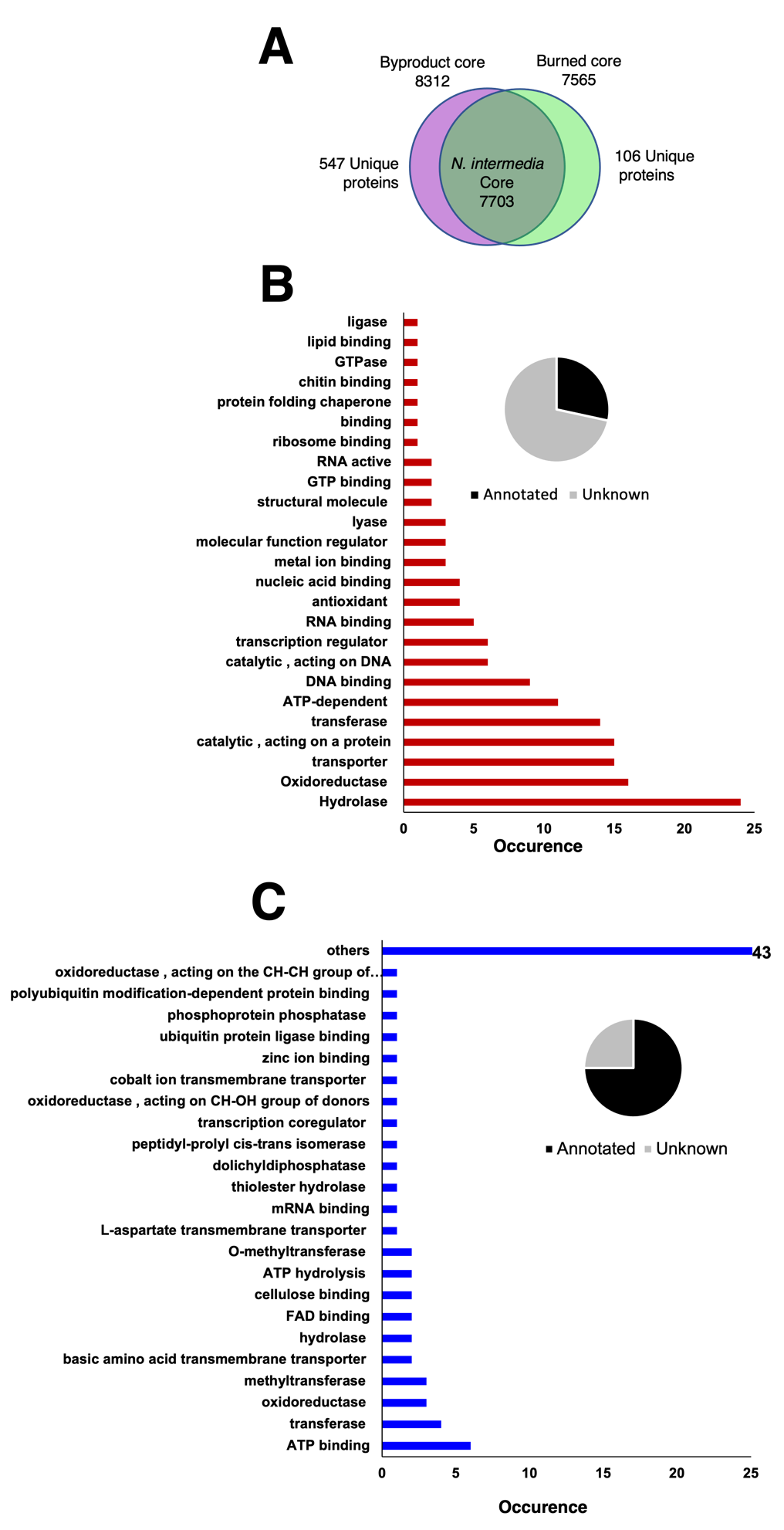

**Figure S11. Pan-genome analysis of two genetically distinct subpopulations of *Neurospora intermedia* associated with byproducts and burned vegetation.** Phylogenetic analysis (Figure 3 in main text) revealed the existence of two closely related, genetically distinct subpopulations of *N. intermedia* associated with burned vegetation and byproducts, respectively. A) The core *N. intermedia* genome consists of 7703 shared protein-coding genes. The pan-genome of the byproduct-associated strains is several-fold larger compared to the burn-associated strains, suggesting potential genetic expansion among the byproduct clade. Functional enrichment analysis of the byproduct associated (B) and burn-associated (C) pan-genomes. BLASTGO was used to identify the predicted metabolic functions among the genes that are unique to each clade. Hydrolytic activity was the most enriched clade among the byproduct-associated strains (B). However, less than half of all genes could be functionally annotated, suggesting many genes with unknown functions. C) In contrast to the byproduct-associated pan-genome, the burn-associated pangenome was enriched in diverse functions and was more disperse in terms of the number of genes for each functionally assigned category. The burn-associated pan-genome also had a higher amount of annotated functions.

**Table S5. *Neurospora* strains analyzed in this study.** Strains of *N. intermedia* experimentally evaluated in this study (see Fig. 3E for exact strains that were evaluated in this study) were obtained from the Fungal Genetics Stock Center (FGSC) ^8^ and their corresponding FGSC numbers are shown. Clade refers to the two clades identified in Figure 3 in the main text. The ecological association data for clades A and B was obtained from FGSC. Svedberg et al (2018) refers to the following paper^1^.

| **Organism** | **Origin** | **Ecological association** | **Genome source** | **Clade** |
| --- | --- | --- | --- | --- |
| *N. intermedia* #1785 | Papua New Guinea | Burned | Svedberg et al (2018) | A |
| *N. intermedia* #1786 | Papua New Guinea | Burned | Svedberg et al (2018) | A |
| *N. intermedia* #1787 | Papua New Guinea | Burned | Svedberg et al (2018) | A |
| *N. intermedia* #1795 | Bali, Indonesia | Not burned, near temple offerings | Svedberg et al (2018) | A |
| *N. intermedia* #1832 | Australia | Burned | Svedberg et al (2018) | A |
| *N. intermedia* #1938 | Papua New Guinea | Burned | Svedberg et al (2018) | A |
| *N. intermedia* #2316 | USA | Burned | Svedberg et al (2018) | A |
| *N. intermedia* #2365 | Hawaii | Burned | Svedberg et al (2018) | A |
| *N. intermedia* #3194 | Papua New Guinea | Burned | Svedberg et al (2018) | A |
| *N. intermedia* #5124 | Tahiti | Burned | Svedberg et al (2018) | A |
| *N. intermedia* #6249 | Haiti | Burned | Svedberg et al (2018) | A |
| *N. intermedia* #6263 | Ivory Coast | Burned | Svedberg et al (2018) | A |
| *N. intermedia* #6595 | Tahiti | Burned | Svedberg et al (2018) | A |
| *N. intermedia* #7402 | Taiwan | No information | Svedberg et al (2018) | A |
| *N. intermedia* #7426 | Malaysia | Burned | Svedberg et al (2018) | A |
| *N. intermedia* #7427 | Taiwan | Burned | Svedberg et al (2018) | A |
| *N. intermedia* #7429 | Taiwan | Burned | Svedberg et al (2018) | A |
| *N. intermedia* #8761 | Taiwan | Burned | Svedberg et al (2018) | A |
| *N. intermedia* #8767 | Bogor, Indonesia | Burned | Svedberg et al (2018) | A |
| *N. intermedia* #8769 | Indonesia | Oncom | Svedberg et al (2018) | B |
| *N. intermedia* #8793 | Papua New Guinea | Corn cob | Svedberg et al (2018) | B |
| *N. intermedia* #2559 | Bogor, Indonesia | Oncom | This study | B |
| *N. intermedia* #2685 | Indonesia | Oncom | This study | B |
| *N. intermedia* #5644 | Indonesia | Oncom | This study | B |
| *N. intermedia* #2557 | Indonesia | Oncom | This study | B |
| *N. intermedia* #1791 | Taiwan | Sugar cane bagasse | This study | B |
| *N. intermedia* #5642 | Taiwan | Sugar cane bagasse | This study | B |
| *N. intermedia* #5342 | Papua New Guinea | Corn cob | This study | B |
| *N. intermedia* #2613 | Indonesia | Oncom | This study | B |
| *N. crassa* #4723 | NA | NA | Svedberg et al (2018) | NA |
| *N. metzenbergii* #10411 | NA | NA | Svedberg et al (2018) | NA |
| *N. metzenbergii* #8853 | NA | NA | Svedberg et al (2018) | NA |
| *N. metzenbergii* P4149 | NA | NA | Svedberg et al (2018) | NA |
| *N. metzenbergii* #5119 | NA | NA | Svedberg et al (2018) | NA |
| *N. metzenbergii* #6795 | NA | NA | Svedberg et al (2018) | NA |
| *N. crassa* #4829 | NA | NA | Svedberg et al (2018) | NA |
| *N. crassa* OR74A | GCF 000182925.2 | NA | Genbank (GCF_000182925.2) ^9^ | NA |
| *N. crassa* #8863 | NA | NA | Svedberg et al (2018) | NA |
| *N. tetrasperma* #2508 | GCF 000213175.1 | NA | Genbank (GCF_000213175.1) ^10^ | NA |
| *N. discreta* #8579 | GCA 009805215 | NA | Genbank (GCA_009805215) (Under Bioproject PRJNA487060) | NA |
| *N. cerealis* #26651 | GCA 009806005.1 | NA | Genbank (GCA_009806005.1) (Under Bioproject PRJNA487060) | NA |
| *N. tetraspora* #26668 | GCA 009806155.1 | NA | Genbank (GCA_009806155.1) (Under Bioproject PRJNA487060) | NA |

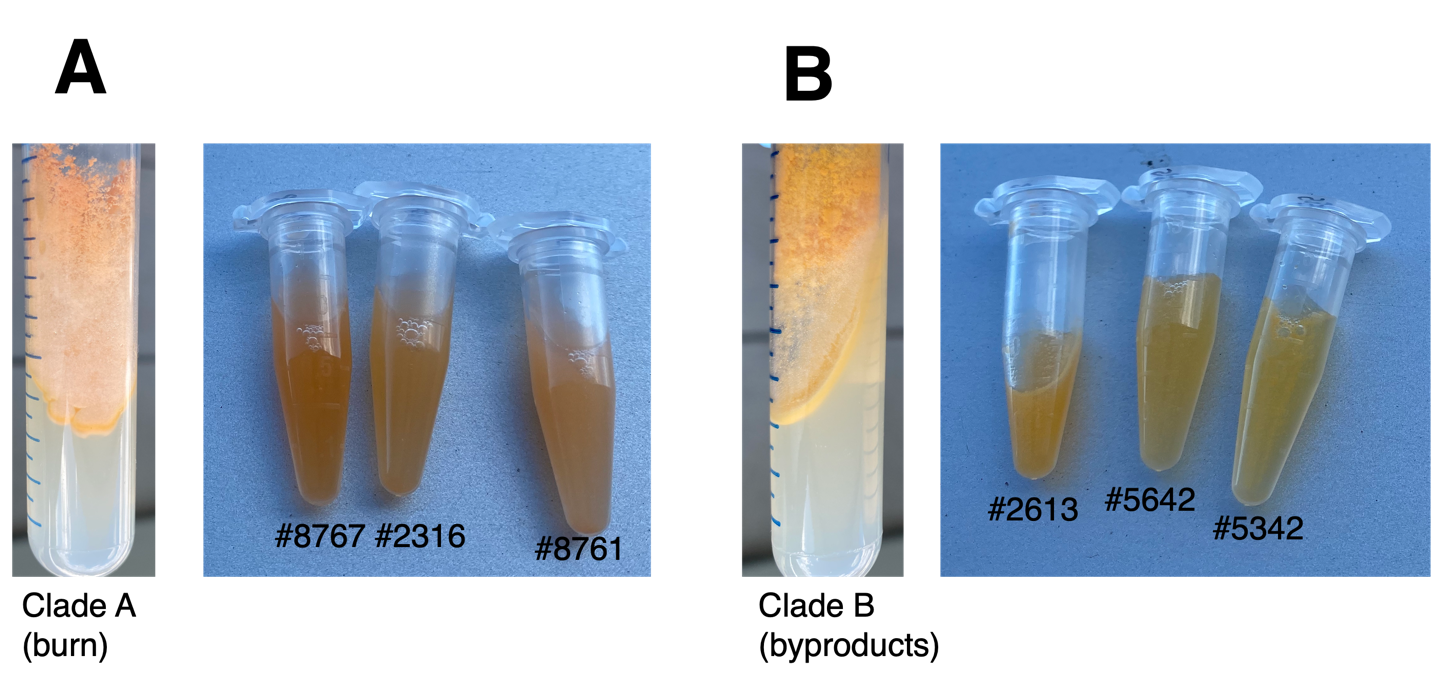

**Figure S12. Representatives from the two clades of *Neurospora intermedia* have previously reported differences in conidial pigmentation*.*** Prior to our work, two types of phenotypically distinct *N. intermedia* strains had been described, which differed in conidial size and pigmentation^11^. We found direct evidence of two genetically distinct clades of *N. intermedia,* which associated with different ecological origins (Figure 3 in main text). Phenotypic analysis confirmed that these two clades had the previously reported differences in conidial pigmentation, suggesting that the spore coloration is consistent with underlying genetic differences between the two subpopulations. A) A representative strain (#2316) from clade A (burn-associated) growing on a slant indicates an orange pigmentation, which is also clearly visible in conidial suspensions of representative members. B) In contrast, byproduct-associated strains have a more yellow-orange coloration, as can be seen during growth on a slant as well as in conidial suspensions of representative strains.

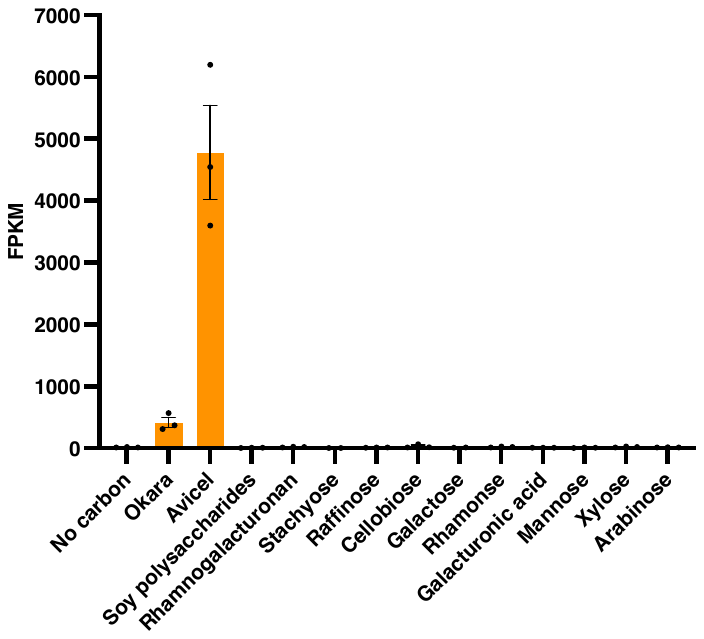

**Figure S13. Expression of predicted cellulase belonging to the glycosyl hydrolase 7 family (#480919) across carbon sources.** Expression of the predicted cellulase belonging to the glycosyl hydrolase 7 family (#480919) that was highly mutated in the burn-associated clade (clade A in Figure 3 in main text) relative to the byproduct-associated clade (clade B in Figure 3 in main text). Results are shown as fragments per kilobase million (FPKM) and are mean and SEM of three biological replicates. The gene was highly expressed on okara, and even higher on avicel, the purified cellulose substrate, implicating this gene in cellulose metabolism.

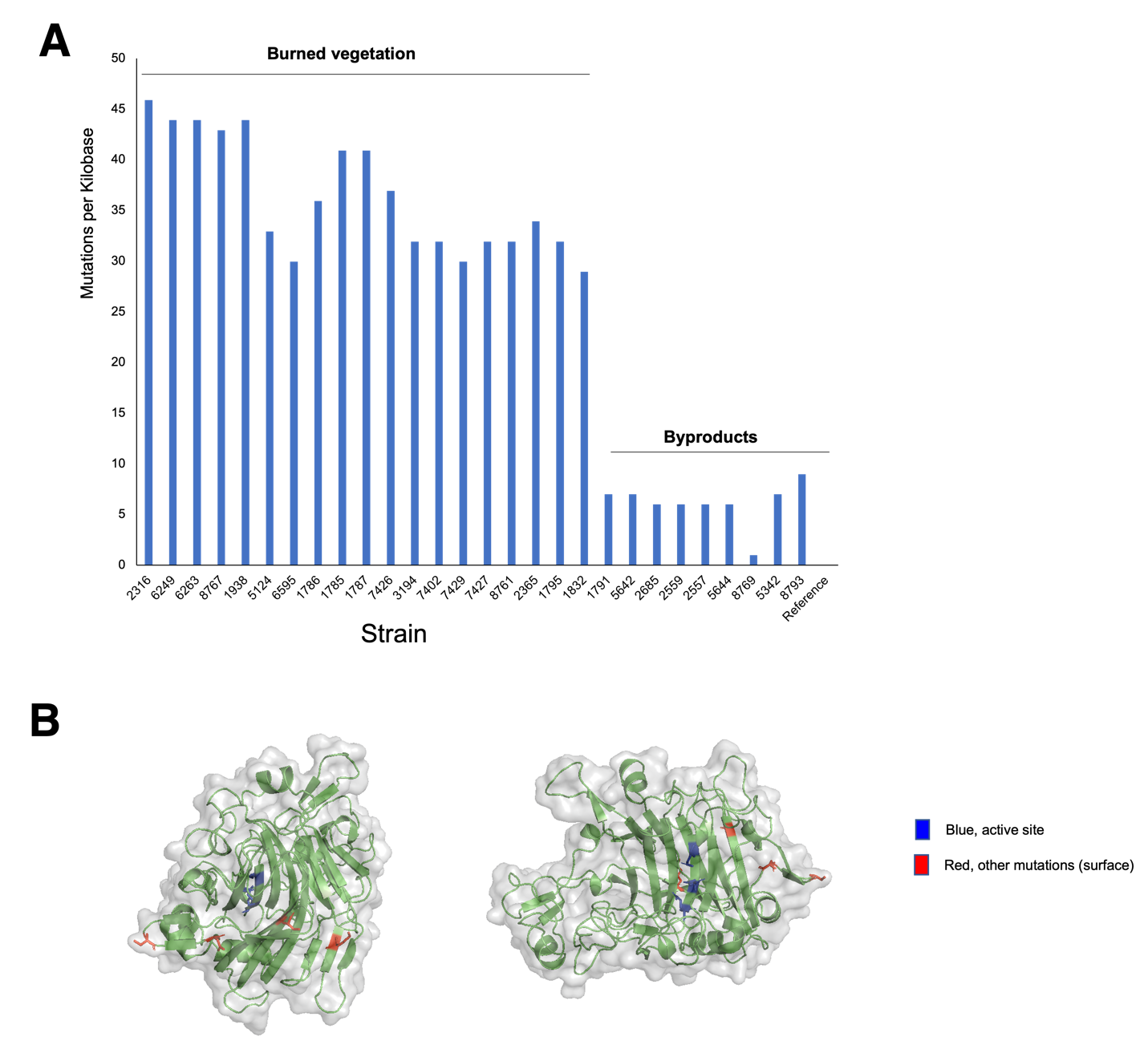

**Figure S14. Mutational profile and structural modeling in the predicted cellulase (#480919) belonging to the glycosyl hydrolase 7 (GH 7) family.** A) SNPs in the enzyme among all *N. intermedia* strains analyzed in this study. Data are normalized per kb. There was a clear difference in the mutational profile between the two groups. B) Structural modeling of predicted mutations in the GH 7 enzyme, based on *T. reesei Cel7B.* Alphafold was used to generate the model. The predicted mutations were located away from the active site and are located on the surface.

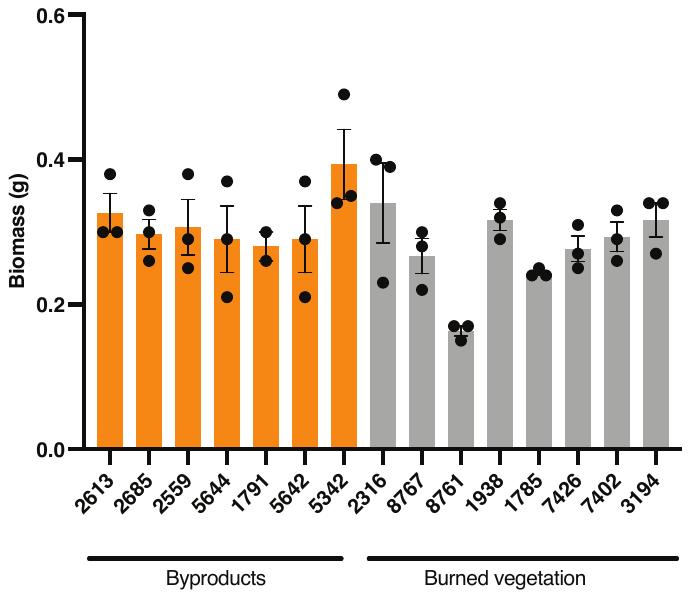

**Figure S15. Biomass yield of byproduct-associated and burned-associated strains on okara.** Strains were grown in liquid cultures using okara as the sole carbon source. Biomass yield was measured after 72 hours of growth. Results are mean + SEM of three biological replicates. There was no significant difference in mean biomass yield between the two groups.

**Table S6. Growth of *N. intermedia* on diverse byproducts.** We quantified the growth by visual inspection and assigned a growth score based on the amount of mycelia that had formed within and around the substrate (0=no growth, 1=minimal growth, 2=moderate growth, 3=high growth). Okara was used as a control to benchmark the high growth. Photos are found in Fig. 4 in main text and Supplementary Data S3.

| **Substrate** | **Growth score** |
| --- | --- |
| Soy milk (okara) waste | 3 |
| Oat milk waste | 3 |
| Black rice milk waste | 3 |
| Carrot pomace | 3 |
| Apple pomace | 3 |
| Malt rootlets | 3 |
| Pumpkin seed presscake | 3 |
| Almond skins | 3 |
| Tiger nut milk waste | 3 |
| Sorghum bagasse | 3 |
| Cashew milk waste | 3 |
| Sunflower seed presscake | 3 |
| Hemp milk waste | 3 |
| Coconut milk waste | 3 |
| Rice hulls | 3 |
| Wheat bran | 3 |
| Barley husks and hulls | 3 |
| Orange peels | 3 |
| Tomato pomace | 3 |
| Almond hulls | 2 |
| Coffee grounds | 2 |
| Banana peels | 2 |
| Pineapple core | 2 |
| Oat bran | 2 |
| Almond milk waste | 2 |
| Spent grain | 1 |
| Hazelnut skins | 1 |
| Grape pomace | 0 |
| Olive pomace | 0 |
| Buckwheat hulls | 0 |

**Table S7. Figure S13. Protein content in a subset of byproducts before and after SSF by *N. intermedia*.** The growth of *N. intermedia* #2613 across byproducts is shown in Figure 4 in the main text. We analyzed the protein content of a subset of byproducts on which *N. intermedia* displayed high growth after 72 hours in SSF. Results shown are % protein by dry weight pre and post fermentation for lyophilized samples.

| **Substrate** | **Pre** | **Post** |
| --- | --- | --- |
| Oat milk byproduct | 22.1 | 28.6 |
| Rice milk byproduct | 18.3 | 33.4 |
| Carrot pomace | 7.5 | 11.6 |

**Table S8. Predicted natural products gene clusters in edible ascomycete fungi.** Predicted natural products production capacity of *N. intermedia* and other sequenced Ascomycete molds used for food production. The results are graphically represented in Figure 4 in the main text. RiPP = ribosomally synthesized and post-translationally modified peptide; NRPS = nonribosomal peptide synthase; PKS = polyketide synthase. Predictions were generated using ANTISMASH.

| **Strain** | **terpene** | **PKS** | **NRPS** | **siderophore** | **indole** | **RiPP** | **Genomes analyzed** |
| --- | --- | --- | --- | --- | --- | --- | --- |
| *Saccharomyces cerevisiae* | 1 | 0 | 2 | 0 | 0 | 0 | 35 |
| *Neurospora intermedia* | 3 | 8 | 9 | 0 | 1 | 1 | 25 |
| *Penicillium roqueforti* | 9 | 16 | 26 | 1 | 4 | 0 | 3 |
| *Penicillium camemberti* | 13 | 23 | 36 | 1 | 4 | 0 | 2 |
| *Fusarium venenatum* | 10 | 13 | 29 | 1 | 2 | 0 | 2 |
| *Aspergillus oryzae* | 14 | 29 | 50 | 1 | 6 | 1 | 96 |
| *Aspergillus sojae* | 16 | 31 | 56 | 2 | 7 | 1 | 4 |

**Table S9. Check all that apply (CATA) sensory analysis of cooked okara fermented with *N. intermedia.*** N=61 Danish consumers who had never before consumed oncom were asked to label the sensory attributes of okara subjected to solid-state fermentation by *N. intermedia* and then cooked in oil in a pan. The sensory attributes are shown alongside their frequency. Attributes with the same frequency are shown in the same cell. Overall, the fermented product was assigned positive attributes such as nutty, mushroom, fermented, fried, earthy, brown, floury, sweet and bready. Together with the liking score in Figure 1 in the main text, those positive attributes suggest that okara fermented with *N. intermedia* can be accepted outside the context in which oncom is traditionally produced and consumed. The sensory attributes were derived from similar trials with Tempeh^12^.

| **Attribute** | **Frequency** |
| --- | --- |
| Nutty | 52 |
| Mushroom | 39 |
| Fried | 32 |
| Fermented | 31 |
| Brown, Earthy | 29 |
| Floury | 26 |
| Bready | 24 |
| Sweet | 23 |
| Long-Lived | 20 |
| Bitter | 16 |
| Dry | 13 |
| Crispy | 11 |
| Sticky | 9 |
| Intense | 8 |
| Crumbly | 7 |
| Sour | 6 |
| Orange | 5 |
| Smoke, Astringent, Rancid | 3 |
| Soft, Moldy | 2 |
| Squishy, Grainy, Cereal, Hay, Mild, Yeasty, Fruity, Umami | 1 |
| Corked, Oily, Crunchy, Metallic, Cheesy, Foie | 0 |

**Table S10. Data availability for *N. intermedia* FGSC #2613 whole-genome sequences and transcriptomics in the public Sequence Read Archive.** Data can be accessed through <https://www.ncbi.nlm.nih.gov/bioproject>. The *N. intermedia* #2613 whole-genome sequence as well as the assembly and annotation are also available through the MycoCosm portal at <https://mycocosm.jgi.doe.gov/Neuin1>.

| **Sequencing type** | **Library Name** | **BioProject** |
| --- | --- | --- |
| Whole-genome sequence | HNZZG | SRP444964 |
| Transcriptomics across C sources | HWOPA | SRP445001 |
| Transcriptomics across C sources | HWOPA | SRP445001 |
| Transcriptomics across C sources | HWOPB | SRP445000 |
| Transcriptomics across C sources | HWOPB | SRP445000 |
| Transcriptomics across C sources | HWOPC | SRP444999 |
| Transcriptomics across C sources | HWOPG | SRP444998 |
| Transcriptomics across C sources | HWOPH | SRP445004 |
| Transcriptomics across C sources | HWOPN | SRP445003 |
| Transcriptomics across C sources | HWOPO | SRP445002 |
| Transcriptomics across C sources | HWOPP | SRP445009 |
| Transcriptomics across C sources | HWOPS | SRP445008 |
| Transcriptomics across C sources | HWOPS | SRP445008 |
| Transcriptomics across C sources | HWOPT | SRP445007 |
| Transcriptomics across C sources | HWOPU | SRP445006 |
| Transcriptomics across C sources | HWOPU | SRP445006 |
| Transcriptomics across C sources | HWOPW | SRP445005 |
| Transcriptomics across C sources | HWOPX | SRP445012 |
| Transcriptomics across C sources | HWOPY | SRP445011 |
| Transcriptomics across C sources | HWOPZ | SRP445010 |
| Transcriptomics across C sources | HWOSA | SRP444979 |
| Transcriptomics across C sources | HWOSB | SRP444977 |
| Transcriptomics across C sources | HWOSB | SRP444977 |
| Transcriptomics across C sources | HWOSC | SRP444978 |
| Transcriptomics across C sources | HWOSC | SRP444978 |
| Transcriptomics across C sources | HWOSH | SRP444975 |
| Transcriptomics across C sources | HWOSO | SRP444976 |
| Transcriptomics across C sources | HWOSP | SRP444971 |
| Transcriptomics across C sources | HWOSP | SRP444971 |
| Transcriptomics across C sources | HWOST | SRP444973 |
| Transcriptomics across C sources | HWOST | SRP444973 |
| Transcriptomics across C sources | HWOSU | SRP444974 |
| Transcriptomics across C sources | HWOSU | SRP444974 |
| Transcriptomics across C sources | HWOSW | SRP444972 |
| Transcriptomics across C sources | HWOSW | SRP444972 |
| Transcriptomics across C sources | HWOSX | SRP444969 |
| Transcriptomics across C sources | HWOSY | SRP444970 |
| Transcriptomics across C sources | HWOSY | SRP444970 |
| Transcriptomics across C sources | HWOTA | SRP444993 |
| Transcriptomics across C sources | HWOTA | SRP444993 |
| Transcriptomics across C sources | HWOTB | SRP444992 |
| Transcriptomics across C sources | HWOTC | SRP444991 |
| Transcriptomics across C sources | HWOTC | SRP444991 |
| Transcriptomics across C sources | HWOTG | SRP444994 |
| Transcriptomics across C sources | HWOTG | SRP444994 |
| Transcriptomics across C sources | HWOTH | SRP444995 |
| Transcriptomics across C sources | HWOTH | SRP444995 |
| Transcriptomics across C sources | HWOTN | SRP444997 |
| Transcriptomics across C sources | HWOTN | SRP444997 |
| Transcriptomics across C sources | HWOTO | SRP444996 |
| Transcriptomics across C sources | HWOTO | SRP444996 |
| Transcriptomics across C sources | HWOTP | SRP444984 |
| Transcriptomics across C sources | HWOTS | SRP444983 |
| Transcriptomics across C sources | HWOTS | SRP444983 |
| Transcriptomics across C sources | HWOTT | SRP444987 |
| Transcriptomics across C sources | HWOTT | SRP444987 |
| Transcriptomics across C sources | HWOTU | SRP444986 |
| Transcriptomics across C sources | HWOTU | SRP444986 |
| Transcriptomics across C sources | HWOTW | SRP444985 |
| Transcriptomics across C sources | HWOTW | SRP444985 |
| Transcriptomics across C sources | HWOTX | SRP444990 |
| Transcriptomics across C sources | HWOTY | SRP444989 |
| Transcriptomics across C sources | HWOTZ | SRP444988 |
| Transcriptomics across C sources | HWOTZ | SRP444988 |
| Transcriptomics across C sources | HWOUA | SRP444980 |
| Transcriptomics across C sources | HWOUB | SRP444981 |
| Transcriptomics across C sources | HWOUB | SRP444981 |
| Transcriptomics across C sources | HWOUC | SRP444982 |
| Transcriptomics across C sources | HWOUC | SRP444982 |

**Table S11. Genome information for additional *N. intermedia* strains sequenced as part of this study and deposited to the Sequence Read Archive.** FGSC numbers are shown next to the species name.

| **Strain** | **Reference accession** | **Length (bp)** | **Contigs** | **G+C content** | **N50** |
| --- | --- | --- | --- | --- | --- |
| *N. intermedia #*2557 | PRJNA996151 | 39841043 | 430 | 0.49 | 158600 |
| *N. intermedia #*2559 | PRJNA996151 | 39871961 | 448 | 0.49 | 162522 |
| *N. intermedia #*2685 | PRJNA996151 | 39952090 | 478 | 0.49 | 146866 |
| *N. intermedia #*1791 | PRJNA996151 | 40039496 | 535 | 0.49 | 136886 |
| *N. intermedia #*5342 | PRJNA996151 | 39544159 | 427 | 0.49 | 183410 |
| *N. intermedia #*5642 | PRJNA996151 | 39932555 | 370 | 0.49 | 199598 |
| *N. intermedia #*5644 | PRJNA996151 | 39943950 | 467 | 0.49 | 153457 |

**Materials and methods**

**Collection and characterization of oncom samples from traditional producers in Java, Indonesia**

Oncom samples were collected from 10 red oncom and 6 black oncom producers throughout Western Java. Prior to sampling, a survey was carried out on the location of the oncom industry and observations related to detailed information on raw materials, the starter culture used, and the production process method used. Samples were taken aseptically and placed in sterile plastic, then the samples were brought to the laboratory using an ice box and immediately used for analysis. Two replicates were collected. One sample was lyophilized for subsequent sequencing analysis. For pH measurements, a total of 50 grams of wet sample was aseptically put into 450 mL of buffer, then crushed and homogenized to obtain a sample suspension of 10% dilution. The pH value was measured using a standardized pH-meter using a buffer. A total of 50 mL of suspension derived was measured for pH and read when the value was stable. Measurements were made twice, then the average was taken. Texture analysis of oncom samples was carried out by visual observation and by using a penetrometer. Visual observation based on the method descried in^13^ by observing the growth of mold covering the surface of oncom and the texture density of oncom with the following scale: + (poor mold growth, non-compact texture); ++ (decent growth of mold, texture is quite compact and dense); +++ (good mold growth, compact and dense texture); +++ (very good mold growth, very compact and dense texture). The measurement of the hardness level uses the Precision Scientific Penetrometer #73501 based on the level of penetration of the penetrometer needle into the sample for a certain time^14^. The oncom sample was placed in the space provided, then the needle was inserted vertically above the surface of the sample at five different points. The measurement was carried out for 5 seconds, so the hardness was expressed as mm/5 seconds.

**16S and ITS amplicon sequencing of oncom samples**

***DNA extraction, PCR, sequencing, and sequence processing:*** Lyophilized oncom samples were ground into a fine powder using a mortar and pestle. Powder was placed into a DNA Isolation Bead Plate. DNA was extracted following Qiagen’s instructions (MagAttract PowerSoil DNA Kit) on a KingFisher robot. Bacterial 16S rRNA genes were PCR-amplified with dual-barcoded primers targeting the V4 region (515F 5’-GTGCCAGCMGCCGCGGTAA-3’, and 806R 5’-GGACTACHVGGGTWTCTAAT-3’) ITS: (ITSF. 5 -CCTCCGCTTATTGATATGC-3  ITS2-Reverse CCGTGARTCATCGAATCTTTG), as per the protocol of Kozich et al. (2013) ^15^. Input DNA concentration was normalized prior to sequencing. Amplicons were sequenced with an Illumina MiSeq using the 250-bp paired-end kit (v.3). Sequenced were analyzed using DADA2^16^ and Phyloseq^17^ following the recommended procedures.

**Metagenome sequencing and analysis of oncom samples**

### *DNA extraction and library preparation:* Lyophilized oncom samples were ground into a fine powder using a mortar and pestle. DNA from the lyophilized material was extracted using the Qiagen MagAttract PowerSoil DNA KF kit (Formerly MOBio PowerSoil DNA Kit) using a KingFisher robot. DNA quality was evaluated visually via gel electrophoresis and quantified using a Qubit 3.0 fluorometer (Thermo-Fischer, Waltham, MA, USA). Libraries were prepared using an Illumina Nextera library preparation kit (Illumina, San Diego, CA, USA) with an in-house protocol from Microbiome Insights, Inc, which performed the sequencing services.

*Sequencing, data curation, and sequence processing:* Paired-end sequencing (150 bp x 2) was done on a NextSeq 500. Shotgun metagenomic sequence reads were processed with the Sunbeam pipeline. Initial quality evaluation was done using FastQC v0.11.5 (Bioinformatics Group at the Babraham Institute. Software available at: <http://www.bioinformatics.babraham.ac.uk/projects/fastqc/>). Processing took part in four steps: adapter removal, read trimming, low-complexity-reads removal, and host-sequence removals. Adapter removal was done using cutadapt v2.6^18^. Trimming was done with Trimmomatic v0.36^19^ using custom parameters (LEADING:3 TRAILING:3 SLIDINGWINDOW:4:15 MINLEN:36). Low-complexity sequences were detected with Komplexity v0.3.6^20^. For metagenomics analysis, the concatenated paired-end read fastq files were analyzed using kaiju v1.9.0^21,22^ (using the NCBI BLAST nr+euk database, which contains bacteria, yeasts, viruses, and microbial eukaryotes) was to perform taxonomic analysis. To reduce noise caused by false positives in identification, a 1% abundance filter was applied to define the presence and absence of each taxon^23^. To evaluate how much of the *N. intermedia* genome was recovered in the sequencing of the oncom samples, all fastq files were concatenated and mapped to the *N. intermedia* genome using HiSAT2.

**General growth conditions and husbandry of *N. intermedia* strains**

*N. intermedia* was grown on Vogels Minimal Medium (VMM) ^24^ unless otherwise indicated. The medium (1 L) was made by combining the 50X salt solution with 15 grams sucrose (unless other carbon source was used) and 1 L distilled water. 15 grams of agar were added for solid medium. To generate spore suspensions, which were used for inoculation in liquid and solid-state cultures, *N. intermedia* was grown from glycerol stocks on VMM agar slants, for at least 72 hours at 30 C, or until robust pigmented conidial growth was established. Conidial suspensions were generated by adding sterile water to the slants followed by vortexing. Conidial concentrations were established using a hemocytometer. Liquid and solid-state cultures were maintained in the dark.

**Preparation of solid-state fermented okara and subsequent sensory analysis and GC-MS analysis of volatile aromas**

***Fermented okara preparation:*** 250 g of raw soybeans were soaked with 4 L of water at 4°C overnight and rinsed with water before processing. It was mixed with three parts water (1:3 w/w), processed in the Thermomix (Thermomix ® TM6) (high speed; 20 s) and drained. Immediately, the solid (okara) was separated from the liquid and steamed in the oven for 30 min at 100 °C. Okara was cooled to 30 °C before inoculating with *Neurospora intermedia* #2613. For inoculation, *N. intermedia* was grown on two separate slants on Vogels Minimal Medium (VMM) for 1 week at 30 °C. A 12 mL conidial suspension was created by adding distilled water to each slant and the resulting 12 mL solution was added to 300 g of steamed grains. The oncom was covered loosely with a sterile cloth and incubated for 24 h at 29 ⁰C and 60 % of relative humidity. After 24 h the spores germinated, and the oncom was fermented without cloth for 24 h. After an additional 24 h, the aerial orange mycelium covered all the surface. The samples were kept at -80 °C until analysis.

***Volatile aroma extraction:*** To 5 g of each sample (wet weight, okara, oncom, cooked oncom), 11 g of water (containing 0.9 % NaCl) was added. The samples were vortexed, placed in stomacher bags (BagPage, Interscience, France), and homogenized in a laboratory blender (Stomacher 400, Seward Co, Worthing, West Sussex, UK) for 2 min at maximum speed. Next, 8 mL were transferred to 20 mL gas tight vials, 2.88 g of NaCl was added, and the samples stored at -80 °C. Ten microlitres of 2-methyl-3-heptanone was used as an internal standard (diluted to 27.2 mg/L in LC/MS-grade methanol). The negative control consisted of distilled water with the same concentration of salt and internal standard^25^. The volatile composition of the samples was determined using headspace solid phase micro-extraction (HS-SPME) in combination with Gas Chromatography-Mass Spectrometry (GC-MS).

***SPME-GC/MS analysis:*** We used the method by (Feng et al., 2013)^25^ with minor modifications. Solid-phase microextraction (SPME) fiber, polydimethylsiloxane/divinylbenzene (PDMS/DVB), 65 µm was used for the extraction of volatile compounds. After the extraction, the samples were incubated at 45 ºC for 15 min, before inserting into the GC injection port. The GC-MS analysis was performed on a Thermo Scientific TRACE 1310 Gas Chromatograph equipped with a Thermo Scientific Q Exactive Orbitrap mass spectrometry system with a Thermo fused-silica capillary column of cross-linked TG-5SILMS (30 m x 0.25 mm x 0.25 µm) (ThermoFisher Scientific, Waltham, MA, USA). The GC conditions were as follows: inlet and transfer line temperatures, 250˚C; oven temperature program, 40˚C for 2 min, 5˚C/min to 120˚C for 2 min, 7˚C/min to 220˚C for 5 min, 50˚C/min to 325˚C for 3 min; inlet helium carrier gas flow rate, 1 mL/min; split ratio, 20:1. The electron impact (EI)-MS conditions were as follows: ionization energy, 70 eV; ion source temperature, 250˚C; full scan m/z range, 30 - 350 Da; resolution, 60,000; AGC target, 1e6; maximum IT, 200ms. Method of identification: (1) by comparison of the MS spectra with the NIST library and (2) by comparison of RI (Kovat indices). The areas were normalized to the ISTD 2-methyl-3-heptanone. Retention index was based on Agilent DB-5MS column using C7-C27 as external references. Data were acquired and analyzed with Thermo TraceFinder 4.1 software package (ThermoFisher Scientific, Waltham, MA, USA).

***Sensory analysis:***

Sensory analysis was conducted by a hedonic test with a total of 61 consumers from Copenhagen, Denmark. Sensory analysis was designed by the Declaration of Helsinki and the 2016/679 EU Regulation on the protection of natural persons regarding the processing of personal data. The study protocol was reviewed and approved by the Ethics Committee at Mondragon Unibertsitatea Before analysis, the experimental procedure was explained and participants completed a written consent form indicating voluntary participation in novel food sensory analysis, as well giving permission for data processing. Okara oncom was prepared to conduct the sensory analysis, following the method above. Approximately 10 g of each sample, was cut in squares, fried, and served. The test was conducted in the same room, under controlled temperature and relative humidity (20 ± 2 ◦C; 95 ± 5% RH) and natural illumination. The nine-point hedonic scale (1 = “dislike extremely”, 9 = “like extremely”) was used to evaluate the participants’ level of liking, as well as three different characteristics (flavour, texture, and appearance). The CATA (Check-All-That-Apply) survey was conducted after the consumer test, with 21 attributes, as shown in Table S3^26^. The attributes used for the sensory attributes were selected based on trials with Tempeh, a similar fungal fermented Indonesian food^27^. All consumers were instructed to rinse their mouths with water between samples.

**Nutritional analysis of raw and fermented okara and other byproducts**

Nutritional analyses were performed by Cumberland Valley Analytical Services, Inc (Waynesboro, PA). All samples analyzed were lyophilized and ground into a fine powder prior to analysis.Fiber (Crude) was analyzed according to the protocol in: Fiber (Crude) in Animal Feed and Pet Food (978.10) described in^28^. Protein (crude) was analyzed by combustion method, following the method of Protein (Crude) in Animal Feed (990.03) described in^28^. The combustion was performed using Leco FP-528 Nitrogen Combustion Analyzer (Leco, 3000 Lakeview Avenue, St. Joseph, MI). Crude Protein was calculated as Nitrogen * 6.25. Fat (crude) was analyzed using the protocol described in Crude Fat in Feeds, Cereal Grains, and Forages (2003.05) described in ^29^. Tecator Soxtec System HT 1043 Extraction unit was used for lipid extraction (Tecator Eden Prairie, MN). Amino acid composition of all amino acids except tryptophan was analyzed using acid hydrolysis of the material and following the protocols described in AOAC Official Method 994.12 and 982.30 described in^29^ , as well as the method described in^30^.

**Tryptophan** was analyzed by alkaline hydrolysis following the protocols in Alkaline hydrolysis (Official Method 988) described in^29^, as well as in^31^. High-performance Liquid Chromatography (HPLC) of amino acids was used for detection, either with an Ion exchange column and Ninhydrin Derivatization^29^, or reverse phase C18 column and fluorescence detector (ISO: 13904:2005)

**Extraction and Liquid Chromatography – Mass Spectrometry (LC-MS) analysis of ergothioneine and free amino acids in raw and fermented okara samples**

***Extraction.*** All samples were lyophilized prior to analysis. For extraction, samples were ground into a fine powder using a mortar and pestle and then approximately 30 mg was transferred to bead beating tubes for homogenization (Lysing Matrix Z, MP Biomedicals, catalog#: 116961050-CF). 1 mL of 20% methanol with 0.1% formic acid was added and samples were subjected to bead beating for 2x1 minutes using the Biospec Mini Beadbeater. Following bead-beating, samples were spun down at 12,000 RCF for 10 minutes to separate the solids. 500 µL supernatant was transferred to a centrifugal spin filter (Amicon Ultra, Sigma-Aldrich, Catalog # UFC500324) to remove any particulates larger molecules (3 kDa cutoff). The flow-through was collected and subjected to analysis by LC-MS.

***LC-MS analysis:*** For LC-MS, analytes were chromatographically separated with a Kinetex HILIC column (100-mm length, 4.6-mm internal diameter, 2.6-µm particle size; Phenomenex, Torrance, CA) at 20°C using a 1260 Infinity HPLC system (Agilent Technologies, Santa Clara, CA, USA). The injection volume for each measurement was 2 µL. The mobile phase was composed of 10 mM ammonium formate (prepared from a pre-made solution from Sigma-Aldrich, St. Louis, MO, USA) and 0.2% formic acid (from an original stock at ≥98% chemical purity from Sigma-Aldrich) in water (as mobile phase A) and 10 mM ammonium formate and 0.2% formic acid in 90% acetonitrile with the remaining solvent being water (as mobile phase B). The solvents used were of LC-MS grade and purchased from Honeywell Burdick & Jackson, CA, USA. Analytes were separated via the following gradient: linearly decreased from 90%B to 70%B in 4 min, held at 70%B for 1.5 min, linearly decreased from 70%B to 40%B in 0.5 min, held at 40%B for 2.5 min, linearly increased from 40%B to 90%B in 0.5 min, held at 90%B for 2 min. The flow rate was changed as follows: 0.6 mL/min for 6.5 min, linearly increased from 0.6 mL/min to 1 mL/min for 0.5 min, held at 1 mL/min for 4 min. The total run time was 11 minutes. The HPLC system was coupled to an Agilent Technologies 6520 Quadrupole Time-of-Flight Mass Spectrometer (QTOF-MS) with a 1:4 post-column split. Nitrogen gas was used as both the nebulizing and drying gas to facilitate the production of gas-phase ions. Drying and nebulizing gases were set to 12 L/min and 25 psi, respectively, and a drying gas temperature of 350°C was used throughout. Fragmentor, skimmer and OCT1 RF voltages were set to 100 V, 50 V and 250 V, respectively. Electrospray ionization (ESI) was conducted in the positive-ion mode with a capillary voltage of 3.5 kV. MS experiments were carried out in the full-scan mode (*m/z* 70–1100) at 0.86 spectra per second for the detection of [M + H]^+^ ions. The instrument was tuned for a range of *m/z* 50 – 1700. Prior to LC-ESI-TOF MS analysis, the TOF MS was calibrated with the Agilent ESI-Low TOF tuning mix. Mass accuracy was maintained via reference ion mass correction, which was performed with purine and HP-0921 (Agilent Technologies). Data acquisition was carried out by MassHunter Workstation Software version B.08.00 (Agilent Technologies). Data processing was carried out by MassHunter Workstation Qualitative Analysis version B.06.00 and MassHunter Quantitative Analysis version 10.00. External calibration curves were used to quantify the analytes.

**High performance anion exchange chromatography (HPAEC) for sugar profiling**

***Extraction of free sugars.*** All samples were lyophilized prior to analysis. For extraction from solid, samples were ground into a fine powder using a mortar and pestle and then approximately 30 mg was transferred to tubes for homogenization (Lysing Matrix Z, MP Biomedicals, catalog#: 116961050-CF). 1 mL of 20% methanol with 0.1% formic acid was added and samples were subjected to bead beating for 2x1 minutes. Following bead-beating, samples were spun down at 12,000 RCF for 10 minutes to separate the solids. 500 µL supernatant was transferred to a centrifugal spin filter to remove any particulates larger molecules (3 kDa cutoff) (Amicon Ultra, Sigma-Aldrich, Catalog # UFC500324). The flow-through was collected and subjected to analysis by HPAEC. For extraction from culture supernatants, 1 mL culture supernatant was collected and subjected to lyophilization. 1 mL of 20% methanol with 0.1% formic acid was added and samples were extracted by continuous vortexing for 10 minutes. Samples were then spun down at 12,000 RCF for 10 minutes and 500 µL supernatant was transferred to a centrifugal spin filter to remove any particulates larger molecules (3 kDa cutoff) (Amicon Ultra, Sigma-Aldrich, Catalog # UFC500324). The flow-through was collected and subjected to analysis by HPAEC.

***Extraction of pectin-bound sugars:*** Method was adapted from Rautengarten et al (2016) ^32^. Lyophilized material was ground to a fine powder before boiling in 96% ethanol for 30 min. After a centrifugation step of 5 min at 20,000g, the supernatant was discarded. The resultant pellet was washed with 70% ethanol until the supernatant was clear. Last, the pellet was washed with 100% acetone and dried in a vacuum concentrator. Samples were hydrolysed in 2 N trifluoroacetic acid (TFA) for 1 h at 120 C.

***High performance anion exchange chromatography (HPAEC) for sugar profiling:*** Method was adapted from Rautengarten et al (2016) ^32^ with minor modifications. High-performance anion exchange chromatography with pulsed amperometric detection was performed on an ICS-6000 (Dionex Corporation, Sunnyvale, CA) using a CarboPac PA20 (3 150 mm, Dionex Corporation, Sunnyvale, CA) anion exchange column at a flow rate of 0.4 ml min 1. Before sample injection, the column was equilibrated with 5 mM NaOH for 5 min. The elution program involved two isocratic elution steps with 5 mM NaOH from 0 to 23 min to separate the neutral sugars followed by a ramp step to 450 mM NaOH from 23.1 to 41 min, which allowed separation of uronic acids and washing of the column. Monosaccharide standards comprised L-Fuc, L-Rha, L-Ara, CGal, D-Glc, D-Xyl, D-GalA and D-GlcA, as well as GlcNac and D-Man. A run of a standard mixture containing the eight monosaccharides was performed with each sample set to enable sample quantitation by linear regression.

**Library preparation and sequencing to generate the high-quality genome of oncom-derived *N. intermedia* FGSC #2613**

***Extraction of high-quality genomic DNA from N. intermedia:*** *N. intermedia* was grown in VMM with sucrose as the sole carbon source at 25 °C, 120 rpm for 72 hours. Mycelia were collected by vacuum filtration and was immediately ground in a mortar and pestle with liquid nitrogen to generate a fine powder. Approximately 12 grams of finely ground mycelium was subjected to further DNA extraction. The mycelium was transferred this to a 50 mL falcon tube containing 10 mL lysis buffer (0.15 M NaCl, 0.1 M EDTA, 2% SDS at pH 9.5). Protease K (Thermo Fisher, #EO0491) was then added at a final amount of 1 mg. The tube was left at 37C shaking at 90 rpm overnight. Samples were then centrifuged at 6000 rcf for 10 minutes to pellet the cellular debris. After centrifugation, avoiding the bottom pellet of debris, 15 mL of the supernatant was transferred to a new tube. 10 ml distilled water was then added to the supernatant, as well as 25 mL of phenol:chloroform:isoamyl alcohol 25:24:1 reagent was then added (Sigma-Aldrich, #P3803). The sample was rotated and shaken to precipitate proteins and was then spun down at 7000 rcf for 15 minutes to separate the layers.  23 mL of the aqueous layer (the top layer) was moved to a new 50 mL falcon tube. Nucleic acids present in this fraction were precipitated with 0.6 volumes (14 mL) of ice-cold isopropanol, which had been frozen at -80 °C for 15 minutes. The precipitated solution was spun down at 7000 rcf for 10 minutes to pellet the DNA. Then 6 mL TE buffer (10 mM Tris, 1 mM EDTA pH 8.0) was added, and the sample was resuspended. RNA was digested by overnight incubation at 37 °C with RNAase A (300 μg, Sigma-Aldrich, #10109142001). Then, an additional protein digestion step was performed by adding protease K (300 μg) and incubating at 37 °C for 2 hours. 6 mL of the solution was extracted once with 6 mL of phenol:chloroform:isoamyl alcohol 25:24:1 reagent (Sigma-Aldrich, #P3803). DNA in the top aqueous layer was precipitated with 0.6 volumes (3.5 mL) of ice-cold isopropanol, and, following centrifugation, the pellet was dried at room temperature, followed by resuspension in TE buffer. To further purify the DNA for PacBio sequencing, we performed an additional cleanup step, which removed contaminating carbohydrates and organic reagents. In this protocol, 150 µL DNA suspension was mixed with 150 µL chloroform and 50 µL TE buffer. Samples were inverted several times to mix. 150 µL of the top layer was removed to a new Eppendorf tube, and 15 µL sodium acetate (pH 5.2) and 450 µL 100% ethanol (ice cold) was added. Following centrifugation to pellet the precipitated DNA, the supernatant was removed, and the pellet was washed twice with 800 µL 70% ethanol, followed by air-drying for 30 minutes at room temperature. The final pellet was re-suspended in 150 µL TE buffer. DNA quality was assessed by agarose gel electrophoresis and nanodrop. DNA concentration was assessed by the Qubit DNA BR assay quantification kit (Q32850). The DNA was deemed sufficiently high quality for reference genome sequencing by PacBio and Illumina.

**Library preparation and genome sequencing of *N. intermedia* #2613:** The high-quality draft genome and full annotations is available through the Joint Genome Institute Website. It has also been deposited to the Sequence Read Archive (SRA) under # 1294447. For the PacBio sequencing (Multiplexed >10kb w/ Blue Pippin Size Selection, Tubes), an input of 1.5 µg of high-quality genomic DNA (extracted according to the protocol above) was sheared around 10 kb using the Megaruptor 3 (Diagenode) or g-TUBE(Covaris). The sheared DNA was treated with exonuclease to remove single-stranded ends, DNA damage repair enzyme mix, end-repair/A-tailing mix and ligated with barcoded overhang adapters using SMRTbell Express Template Prep Kit 2.0 (PacBio). Up to sixteen libraries were pooled in equimolar concentrations and purified with AMPure PB Beads (PacBio). Pooled libraries were size selected using the 0.75% agarose gel cassettes with Marker S1 and High Pass protocol on the BluePippin (Sage Science). PacBio Sequencing primer was then annealed to the SMRTbell template library and sequencing polymerase was bound to them using Sequel II Binding kit 2.0. The prepared SMRTbell template libraries were then sequenced on a Pacific Biosystems' Sequel IIe sequencer using sequencing primer, 8M v1 SMRT cells, and Version 2.0 sequencing chemistry with 1x1800 sequencing movie run times.

**Genome assembly and annotation:** Filtered subread data was processed with the JGI QC pipeline to remove artifacts. The mitochondria-filtered CCS reads were then assembled with Flye version 2.8.1-b1676 (<https://github.com/fenderglass/Flye>) [-g 40M --asm-coverage 50 -t 32 --pacbio-hifi] and subsequently polished with two rounds of RACON version 1.4.13 racon [-u -t 36] (<https://github.com/lbcb-sci/racon>). Small scaffolds (<1kb) were removed from the assembly. The cleaned nuclear genome assembly was annotated using the JGI Annotation Pipeline (Grigoriev et al., 2014). The annotation pipeline uses a combination of ab initio, homology-based, and transcriptome-based gene predictors. The best representative gene model at each locus was selected through automated filtering based on homology and transcriptome support to produce the gene model sets available on MycoCosm^33^ (Grigoriev et al., 2014).

**Growth of *N. intermedia* FGSC #2613 across carbon sources and extraction of RNA for sequencing**

***Growth experiment across carbon sources:*** We adapted the approach used in *Neurospora crassa* by Wu et al (2020) ^34^, with minor modifications. For all RNAseq experiments, we first grew *N. intermedia* #2613 from a glycerol stock for 7 days on VMM slants harboring sucrose as the sole carbon source. Conidia were then harvested by addition of 4 mL sterile water to the slant, followed by vortexing. Conidia were counted using a hemocytometer and were added at a final concentration of 5*10^5^ conidia / mL into 3 mL VMM medium harboring 2% (w/v) sucrose at in 24 well Whatman Uniplates. Each well harbored 3 mL medium. Importantly, the bottoms of the wells for each Uniplate were initially scratched with a sharp syringe needle to allow adherence and formation of mycelial mat. The plates were put in a shaker (light on) for 2 hours to enable conidial germination and initial mat formation. This step was necessary to establish a mat that could be manipulated in subsequent steps. Then, the shaker was turned on (200 rpm, light on). After 15 hours of growth, mycelial mats were removed from the bottom of the plate where they had attached and were then washed three times in 3 mL of VMM without any added carbon, to remove the sucrose medium. The mycelial mats were finally transferred to 1 x VMM with the indicated carbon source for induction. All conditions were done in triplicate. In addition to the carbon sources, a set of samples were “induced” in a no carbon control, which facilitated downstream analysis to identify genes that were uniquely induced on the carbon source of interest. For induction conditions with carbon sources, 2mM mono and disaccharides were used, while for complex carbon sources, including complex polysaccharides and plant biomass, 1% (w/v) was used. The following complex carbon sources were used in the experiments at 1% (w/v): Okara flour (Renewal Mill, Okland, CA), Soluble soy polysaccharies (Creative Enzymes Inc, #NATE-1284), Avicel PH-101 (Sigma-Aldrich, #11365), rhamonogalacturonan from soy pectic fibre (MEGAZYME, #P-RHAGN), and Stachyose hydrate (Sigma-Aldrich, #S4001-100MG). The following carbon sources were used in the experiment at 2 mM: D-(+)-RAFFINOSE PENTAHYDRATE (TCI, #R0002), D-(+)-Galactose (Sigma-Aldrich, #G0750-10G), L-(+)-Arabinose (Sigma-Aldrich, #A3256-25G), D-(+)-Xylose (Sigma-Aldrich, #X3877-25G), D-(+)-Galacturonic acid monohydrate (Sigma-Aldrich, #48280-25G-F), L-Rhamnose Monohydrate (Sigma-Aldrich, #83650-50G), D-(+)-Cellobiose (Sigma-Aldrich, #C7252-100G), and D-(+)-Mannose (MP Biomedicals, # 02102250-CF). After 4 hours of induction, mycelia were harvested over Miracloth filter paper and flash frozen in liquid nitrogen for storage at -80˚C in Lysing Matrix Z tubes (MP Biomedicals, catalog#: 116961050-CF), which allowed subsequent bead beating upon defrosting for RNA extraction. Three biological replicates were used for each condition.

***RNA extractions.*** RNA extractions were performed on -80˚C stored biomass using TRIzol (Invitrogen, # 15596026) and chloroform. Half of the mycelial biomass from a 3 mL culture was used for RNA extractions. 1 mL of TRIzol reagent was added to the bead beating tube. Tubes containing biomass, TRIzol, and beads were bead beaten for 1 minute, then allowed to incubate at room temperature for 5 minutes on a rocker. 200 µL of chloroform was then added to each tube, which was vortexed and centrifuged for phase separation. 400 µL of the aqueous phase from each sample was combined with 400 µL isopropanol and incubated at room temperature for 10 min on a rocker for RNA precipitation. Samples were centrifuged at 4˚C for 10 mins. RNA pellets that formed were washed with 75% ethanol and centrifuged at 4˚C for 2 min. Ethanol was removed via pipet, and the RNA pellet was allowed to dry for several minutes with the cap open. The RNA pellet was resuspended in 40 µL water and treated with 2 µL of Turbo DNAse (Thermo Fisher), in a 50ul reaction. After 20 min incubation at 37˚C, RNA was cleaned up using Qiagen RNeasy Mini Kit (Qiagen), eluting at the final step in 30 µL RNAse-free water. RNA was tested for quality using agarose gel electrophoresis and nanodrop. RNA was quantified using the Qubit RNA quantification kit (Thermo Fisher, #Q10210).

**Library preparation and sequencing of *N. intermedia* FGSC #2613 transcriptomes**
Illumina sequencing was used to capture the transcriptomic profile of *N. intermedia* #2613 grown across carbon sources. RNA was extracted according to the protocol above. Plate-based RNA sample prep was performed on the PerkinElmer Sciclone NGS robotic liquid handling system using Illumina's TruSeq Stranded mRNA HT sample prep kit utilizing poly-A selection of mRNA following the protocol outlined by Illumina in their user guide, and with the following conditions: total RNA starting material was 1 µg per sample and 8 cycles of PCR was used for library amplification. The prepared libraries were then quantified using KAPA Illumina library quantification kit (Roche) and run on a LightCycler 480 real-time PCR instrument (Roche). The quantified libraries were then multiplexed and the pool of libraries was then prepared for sequencing on the Illumina NovaSeq 6000 sequencing platform using NovaSeq XP v1.5 reagent kits (Illumina), S4 flow cell, following a 2x150 indexed run recipe. The transcriptomics data from *Neurospora intermedia* FGSC #2613 grown across carbon sources has been deposited to the Sequence Read Archive (SRA) under PRJNA982896-PRJNA982905, and PRJNA982926-PRJNA982963. The access information is specified in Table S10 in the supplemental materials.

**Analysis of transcriptome data from *N. intermedia* FGSC #2613 grown across carbon sources**

***RNA-seq data processing:*** Raw fastq files reads were filtered and trimmed using the JGI QC pipeline resulting in the filtered fastq file (*.filter-RNA.gz files). Using BBDuk^35^, raw reads were evaluated for artifact sequence by kmer matching (kmer=25), allowing 1 mismatch and detected artifact was trimmed fro the 3' end of the reads. RNA spike-in reads, PhiX reads and reads containing any Ns were removed. Quality trimming was performed using the phred trimming method set at Q6. Finally, the reads under the length threshold were removed. Filtered reads from each library were aligned to the reference genome using HISAT2 version 2.2.0^36^. Strand-specific coverage bigWig files (fwd and rev) were generated using deepTools v3.1^37^. featureCounts^38^ was used to generate the raw gene counts (counts.txt) file using gff3 annotations. Only primary hits assigned to the reverse strand were included in the raw gene counts (-s 2 -p -- primary options). Raw gene counts were used to evaluate the level of correlation between biological replicates using Pearson's correlation.

***Differential expression analysis****:* To determine those genes that change significantly, we only consider as expressed those genes that have at least a sum of 10 counts per million in at least fifteen libraries. All the comparisons were paired and the profile in each carbon source was compared against the non-carbon source condition; this was done with the edgeR package^39^. We used the generalized linear model (GLM) likelihood ratio test. To determine if there is consistency between biological replicates we performed a multidimensional scaling plot of distances between gene expression profiles. The parameters used to call a gene DE between conditions were FDR < 0.05 and log2 fold-change > |1|. A common dispersion between replicates of 0.01339982 was calculated and a tagwise dispersion was also calculated. All the data generated in this work were deposited on the page <https://genome.jgi.doe.gov/portal/pages/dynamicOrganismDownload.jsf?organism=Neuin1>. Note: Raw gene counts (counts.txt), not normalized counts, were used for differential gene expression analysis.

***Annotation of the genes****:* To annotate genes, orthologous genes between *N. intermedia* and *N. crassa* were obtained. This was done with the OrthoFinder program^40^. The curated annotation was inherited from *N. crassa* to *N. intermedia.* The general classification as “PlantDegradBio” is based on the *Neurospora crassa* genes predicted to code for plant biomass degrading enzymes and transporters, presented in Supplementary Table 2 published by Wu et al (2020) ^34^.

***Co-expression network****:* With the matrix of accounts, we built a co-expression network using the WGCNA software^41^. The transformation of the count table was performed with the VST (Variance Stabilizing Transformation) function of DESeq2^42^. Subsequently, the Pearson correlation matrix and the weighted adjacency matrix with continuous values of 0 and 1 were obtained. To evaluate scale independence and mean connectivity, we utilized a gradient method by systematically adjusting the power value from 1 to 20. After identifying a degree of independence surpassing 0.90, the construction of a scale-free network was initiated using the blockwiseModules function, employing a power value of 8. To define the modules of the network, the minimum size was established at 30 genes per module and the threshold for merging similar modules was established at 0.15. Furthermore, the weighted adjacency matrix was transformed into a topological overlap measurement matrix (TOM) to estimate connectivity in the network. Finally, we export the network with the function exportNetworkToCytoscape with a correlation threshold of 0.25. The network was visualized using the Cytoscape v.3.9.1 program. Network metrics were obtained by Cytoscape v.3.9.1. To determine enriched metabolic pathways in the modules we used the KEGG tool (https://www.genome.jp/kegg/).

**Time-course profiling of galactose and arabinose in liquid cultures of of *N. intermedia* #2613 using okara as the sole carbon source**

5*10^5^ conidia from *N. intermedia* #2613 were inoculated into 50 mL VMM medium harboring okara flour as the sole carbon source (1% w/v). 250 mL flasks were used. Flasks were then left to shake at 30 C at 160 rpm in the dark. 1 mL culture supernatants were removed at regular intervals, spun down to remove the solid material and mycelium, and immediately flash-frozen at -80 C. Clarified culture supernatants were then lyophilized and were then extracted for sugar analysis using HPAEC (see below).

**Enzyme assays for cellulase detection**

For all cellulase assays, we used the fluorescence-based cellulase kit from Abcam (Abcam, #ab189817) and followed the standard protocol. All cellulase assays were performed in 96-well plates (Corning, Falcon Tissue Culture Plate, #353072) and fluorescence was measured at 550 nm (excitation) and 595 nm (emission) using a BIOTEK Synergy H1 microplate reader (Agilent). Readings were recorded every 3 minutes for 18 minutes total. For enzyme assays from liquid cultures, 1 mL culture supernatants were removed from cultures by pipetting. These were then spun down to pellet the insoluble material and mycelium. 20 or 25 µL of the clarified supernatant was then used as the enzyme source in cellulase assays in the 100 µL reactions. For cellulase assays from solid state samples, approximately 300 mg (wet weight) of raw or fermented okara was placed in an Eppendorf tube. 1 mL water was added followed by vortexing. To homogenize the sample, sonication was used (10 s on, 25 s OFF, 2 min total, 25 % amplitude). The resulting slurry was spun down at max speed to clarify the supernatant, and 25 µL of the resulting supernatant was used as the enzyme source in the 100 µL enzyme reactions. Cellulase activity was calculated using the formula (𝐵/(Δ𝑇∗ 𝑉))∗𝐷, where B = amount of resorufin in sample well calculated from Standard Curve (µmol); ΔT = linear phase reaction time T2 – T1 (minutes); V = original sample volume added into the reaction well (µL); D = sample dilution factor if sample is diluted to fit within the standard curve range. Results from calculations were expressed as mU / mL. 1 Unit (U) was defined as the amount of enzyme that cleaves substrate to generate 1.0 μmol of molecule per min.

**Cellulase activity in liquid cultures of *N. intermedia* strains**

***Confirmation of cellulase activity in liquid cultures of N. intermedia #2613:*** to confirm the presence of cellulases suggested by the RNA-seq data, we replicated the RNA-seq culture setup in which the cellulase gene expression was first detected. We first grew *N. intermedia* #2613 from a glycerol stock for 7 days on VMM slants harboring sucrose as the sole carbon source. Conidia were then harvested by addition of 4 mL sterile water to the slant, followed by vortexing. Conidia were counted using a hemocytometer and were added at a final concentration of 5*10^5^ conidia / mL into 3 mL VMM medium harboring 2% (w/v) sucrose at in 24 well Whatman Uniplates. The bottoms of the wells for each Uniplate were initially scratched with a sharp syringe needle to allow adherence and formation of mycelial mat. The plates were put in a shaker (light on) for 2 hours to enable conidial germination and initial mat formation. This step was necessary to establish a mat that could be manipulated in subsequent steps. Then, the shaker was turned on (200 rpm, light on). After 15 hours of growth, mycelial mats were removed from the bottom of the plate where they had attached and were then washed three times in 3 mL of VMM without any added carbon, to remove the sucrose medium. The mycelial mats were finally transferred to 1 x VMM with either sucrose (1.5% w/v), avicel (1% w/v), okara flour (1% w/v), or rhamnogalacturonan (1% w/v). All conditions were done in triplicate. Cultures were left for 72 hours, and supernatants were then analyzed for cellulase activity using the protocol above.

***Screen for secreted cellulase activity in liquid cultures among strains of N. intermedia grown on okara:*** to confirm screen for cellulase activity across burn-associated and byproduct-associated *N. intermedia* strains, we grew suggested by the RNA-seq data, we replicated the RNA-seq culture setup. We first grew *N. intermedia* strains from glycerol stocks for 7 days on VMM slants harboring sucrose as the sole carbon source. Conidia were then harvested by addition of 4 mL sterile water to the slant, followed by vortexing. The following strains were used from the byproduct-associated clade: #2613, #2685, #2559, #5644, #1791, #5642, #5342). The following strains were used from the burn-associated clade: #1316, #8767, #8761, #1938, #1785, #7426, #7402, #3194). Conidia were counted using a hemocytometer and were added at a final concentration of 5*10^5^ conidia L into 250 mL flasks with 50 mL VMM medium harboring 1% (w/v) okara flour as the sole carbon source. All strains were grown in triplicate and the experiment was repeated three times to verify the consistency of the results. Cultures were left for 72 hours, and supernatants were analyzed for cellulase activity using the protocol above. Biomass was harvested and dried at the end of the experiment. The biomass was dried for 7 days at 50 C prior to the weight being recorded.

**Genome sequencing of additional *N. intermedia* strains beyond the oncom-derived reference strain *N. intermedia* #2613**

***Extraction and sequencing:*** *N. intermedia* strains subjected to sequencing (FGSC #2559, #2685, #5644, #2557, #1791, #5642, #5342) were grown in VMM-sucrose medium (50 mL medium in 250 mL flasks) for 72 hours prior to harvesting by vacuum filtration and flash-freezing in liquid nitrogen. gDNA extraction, sample quality assessment, DNA library preparation, sequencing and bioinformatics analysis were conducted at Azenta Life Sciences (South Plainfield, NJ, USA). Genomic DNA was extracted using DNeasy Plant Mini Kit following manufacturer’s instructions (Qiagen, USA). Genomic DNA was quantified using the Qubit 2.0 Fluorometer (ThermoFisher Scientific, Waltham, MA, USA). NEBNext® Ultra™ II DNA Library Prep Kit for Illumina, clustering, and sequencing reagents was used throughout the process following the manufacturer’s recommendations. Briefly, the genomic DNA was fragmented by acoustic shearing with a Covaris S220 instrument. Fragmented DNA was cleaned up and end repaired. Adapters were ligated after adenylation of the 3’ends followed by enrichment by limited cycle PCR. DNA libraries were validated using a High Sensitivity D1000 ScreenTape on the Agilent TapeStation (Agilent Technologies, Palo Alto, CA, USA), and were quantified using Qubit 2.0 Fluorometer. The DNA libraries were also quantified by real time PCR (Applied Biosystems, Carlsbad, CA, USA). The sequencing library was clustered onto lanes of an Illumina HiSeq 4000 (or equivalent) flowcell. After clustering, the flowcell was loaded onto the Illumina HiSeq instrument according to manufacturer’s instructions. The samples were sequenced using a 2x150bp Paired End (PE) configuration. Image analysis and base calling were conducted by the HiSeq Control Software (HCS). Raw sequence data (.bcl files) generated from Illumina HiSeq was converted into fastq files and de-multiplexed using Illumina bcl2fastq 2.17 software. One mismatch was allowed for index sequence identification.

***Assembly and annotation:*** the reads were filtered with TrimmomaticPE version 0.39^19^ with the following parameters: LEADING:30 TRAILING:30 MINLEN:120. The filtered reads were used for *de novo* assembly using the SPAdes genome assembler v3.13.1-1^43^ with the following parameters --careful --cov-cutoff 100. The resulting assemblies were then processed with AUGUSTUS v3.4.0^44^, to obtain coding sequences and protein predictions. Augustus was executed using a gene model for *Neurospora crassa* to identify start and stop codons, introns, and exons.

**Genomic and Phylogenomic analysis of *N. intermedia* strains**

***Core/ pan genome analysis for N. intermedia:*** in addition to the seven *N. intermedia* draft genomes obtained in our study using short reads, and the genome of *N. intermedia* strain 2613 obtained using long and short reads, we retrieved the genomes of 29 *Neurospora* spp. strains from a previous study^1^, and 5 strains which were obtained from the public GenBank database. These genomes were annotated as following the protocol described above and their protein sequences were used for core genome calculation using BPGA^45^ with a sequence identity cutoff of 0.5, this analysis led to a set of 6416 conserved orthologous proteins. The functional analysis of the resulting sets of core and unique genes was done with blast2go^46^ implemented in omicsBox v3.029 (<https://www.biobam.com/omicsbox>).

The taxonomic affiliation of the *N. intermedia* strains used in this study (tree in figure 4) was defined by a multi-locus phylogenetic tree constructed with the set of conserved orthologs proteins found in their core genome. For each set of orthologs, the amino acid sequences were aligned^1^ and trimmed^47^, after this process 6353 alignments were kept. They were then concatenated to form a matrix with 3203239 sites in 6353 partitions. An evolutionary model was calculated for each partition. Then a phylogenetic tree was calculated with IQtree2 v2.0.7^48^ using maximum likelihood with 10,000 bootstrap replicates. The entire process was executed automatically using a script available at <https://github.com/WeMakeMolecules/Core-to-Tree>

***SNP analysis:*** for the identification and analysis of Single Nucleotide polymorphisms (SNPs) we assembled a set of DNA sequencing reads for a total of 28 *N. intermedia* strains obtained with the Illumina Hi-seq platform in the 2X150 paired end read format. This dataset included the reads obtained for the six *N. intermedia* strains sequenced in this project and 22 more from a previous study^1^. Strain FGSC #2613 (SRA accession number #1294447) obtained for this project was used as reference.

The reads were downloaded from the genbank FTP using the fastq-dump v2.113 (<https://github.com/ncbi/sra-tools>). Each of the 28 paired reads were aligned with Bowtie 2 v2.4.4^49^, the Sequence Alignment and Map (SAM) formatted files where then converted to binary alignment and map format (BAM) and sorted using samtools v1.13^50^. The sorted and bam-formatted alignment and mapping files were then used for variant calling, using the *mpileup*, *call*, *view*, and *filter* utilities of the bcftools package v1.13^50^. For this we set the ploidy to 1 and a quality cutoff for the SNPS of QUAL=>50. A simple script to perform these operations is available at: <https://github.com/WeMakeMolecules/SNP_CALL_and_MAPPING/raw/main/map_and_vcf.sh>

We used a simple script to process the variant call files (vcf formatted) to obtain a table (Supplementary Data S2) with the number of SNPs per gene in each of the analyzed strains when compared with strain 2613. The script is available at: <https://github.com/WeMakeMolecules/SNP_CALL_and_MAPPING/raw/main/raw_snps_counter.pl>.

The raw SNP counts were divided by the length of their corresponding gene in kilobases to obtain the SNP per Kilobase value shown in Supplementary Data S2. Finally, to select for genes that have accumulated significantly more mutations and that may be implied in the emergence of new food-assocaited traits, e. g. outliers, we (i) identified genes which SNP/Kb value is equal or more than the average SNP/Kb value for that gene plus two standard deviations (e. g. 2 X STDEV + average) then (ii) we selected genes which values passed this filter in at least 10 of the strains locate in the burn-vegetation clade (figure 4B). The resulting list of genes is in Supplementary data S2. The functional analysis of the resulting set of outlier genes was done with blast2go^46^implemented in omicsBox v3.029 (<https://www.biobam.com/omicsbox>).

**Screen for growth of *N. intermedia* across okara and other byproducts in solid state cultures**

Byproducts were received from restaurants and large-scale industrial food producers who wished to remain anonymous. Byproducts that were received in dry form soaked overnight and then drained to remove excess moisture and accomplishing a % moisture of approximately 50 % w/w. Byproducts were then autoclaved for 15 minutes at 121 C for sterilization. For inoculation, we first grew *N. intermedia* #2613 from a glycerol stock for 7 days on VMM slants harboring sucrose as the sole carbon source. Conidia were then harvested by addition of 4 mL sterile water to the slant, followed by vortexing. 2 mL of this conidial suspension was added to approximately 50 g of the solid material, which was then mixed by hand to distribute the conidia. The inoculated byproducts were placed in square petri dishes with the lid on (Greiner Bio, #688102) and grown at 30 C for 72 hours. Autoclaved, uninoculated byproducts were saved for comparison. Growth was assessed by visual inspection, and a growth score was assigned based on the development of mycelia and conidia within and around the substrate, and compared with okara, which served as a positive control to benchmark the high growth. The score was as followed: 0=no growth, 1=minimal growth, 2=moderate growth, 3=high growth. Samples were lyophilized for storage, and a subset of samples were subjected to protein analysis.

**Natural products prediction across *N. intermedia* and other edible ascomycetes**

For the prediction of natural product production capacity (Fig. 4C) the genome assembly and gene calling file for strain 2613 was then used for functional annotation and mining for natural products biosynthetic gene clusters using Antismash version 7^51^. The same gene calling, and annotation process described above in *“Genome sequencing of additional N. intermedia strains beyond the oncom-derived reference strain N. intermedia #2613”* was done for the genomes of *Saccharomyces cerevisiae* YJM1342 (GenBank accession GCA_000977265.3), *Fusarium venenatum* A3/5 (GenBank accession GCA_900007375.1), *Penicillium roqueforti* JCM 22842 (GenBank accession GCA_001599855.1), *Penicillium camemberti* FM013 (GenBank accession GCA_014839975.1), *Aspergillus oryzae* RIB40 (GenBank accession GCF_000184455.2) and *Aspergillus sojae* SMF134 (GenBank accession GCA_008274985.1).

**Metabolomic analysis of secondary metabolite production of *Neurospora intermedia***

***Extraction:*** lyophilized, powdered samples of *Neurospora Intermedia* (2 mg) grown in okara obtained by SSF were transferred to vials, followed by the addition of 2 ml of methanol. The vials were placed in an ultrasound bath for 60 min. At the end of this period, centrifugation was performed, and the supernatant was then transferred to a 1.5 mL glass vial in preparation for subsequent LC-MS/MS analysis.

***LC-MS/MS data acquisition*** ***and*** ***untargeted metabolomics analysis:*** untargeted metabolomic analysis of extracts was performed using LC-MS/MS on a Vanquish Duo UHPLC binary system (Thermo Fisher Scientific, USA) coupled to an IDX-Orbitrap Mass Spectrometer (Thermo Fisher Scientific, USA). The chromatographic separation was performed using a Waters ACQUITY BEH C18 column (10 cm × 2.1 mm, 1.7 μm) equipped with an ACQUITY BEH C18 guard column kept at 40°C, with a flow rate of 0.35 mL/min. The mobile phases consisted of MilliQ© water + 0.1% formic acid (A) and acetonitrile + 0.1% formic acid (B). The initial composition was 2% B held for 0.8 min, followed by a linear gradient to 5% in 3.3 min, and then a second gradient until reaching 100% B in 10 min. This condition was then held for 1 min. The system re-equilibration time was 2.7 min before loading the next sample into the instrument. The MS measurement was done in positive and negative-heated electrospray ionization (HESI) mode with voltages of 3500 V and 2500 V, respectively. Full MS/MS spectra DDA (Data-dependent Acquisition-driven MS/MS) were acquired within the mass range of 70–1000 Da. The DDA acquisition settings were as follows: automatic gain control (AGC) target value set at 4e^5^ for full MS and 5e^4^ for MS/MS spectral acquisition, and the mass resolution was set to 120,000 for full scan MS and 30,000 for MS/MS events. Precursor ions were fragmented by stepped High-energy collision dissociation (HCD) using collision energies of 20, 40, and 60. To search for known natural products and mycotoxins in *Neurospora intermedia* grown in okara the tandem mass spectrometry data was analyzed using the Global Natural Product Social Molecular Networking (GNPS) ^52^ platform (available online through <https://gnps.ucsd.edu/ProteoSAFe/static/gnps-splash.jsp>; online portal accessed on 24 October, 2022) and SIRIUS 4^53^. For MS/MS dereplication via molecular networking analysis and SIRIUS 4, MS/MS data were converted to mzXML format using MS-Convert software, which is part of ProteoWizard (Palo Alto, CA, USA).

**References**

1	Svedberg, J. *et al.* Convergent evolution of complex genomic rearrangements in two fungal meiotic drive elements. *Nat Commun* **9**, 4242 (2018). https://doi.org:10.1038/s41467-018-06562-x

2	Zhaojun Wang, T. G., Zhiyong He, Maomao Zeng, Fang Qin, Jie Chen. Reduction of off-flavor volatile compounds in okara by fermentation with four edible fungi. *LWT* **55** (2022). https://doi.org:https://doi.org/10.1016/j.lwt.2021.112941

3	Lan, Y., Xu, M., Ohm, J. B., Chen, B. & Rao, J. Solid dispersion-based spray-drying improves solubility and mitigates beany flavour of pea protein isolate. *Food Chem* **278**, 665-673 (2019). https://doi.org:10.1016/j.foodchem.2018.11.074

4	Cavinato, C., Da Ros, C., Pavan, P. & Bolzonella, D. Influence of temperature and hydraulic retention on the production of volatile fatty acids during anaerobic fermentation of cow manure and maize silage. *Bioresour Technol* **223**, 59-64 (2017). https://doi.org:10.1016/j.biortech.2016.10.041

5	Lin, J. *et al.* Qualitative and quantitative analysis of volatile constituents from latrines. *Environ Sci Technol* **47**, 7876-7882 (2013). https://doi.org:10.1021/es401677q

6	Siddique, R., Zahoor, A. F., Ahmad, H., Zahid, F. M. & Karrar, E. Impact of different cooking methods on polycyclic aromatic hydrocarbons in rabbit meat. *Food Sci Nutr* **9**, 3219-3227 (2021). https://doi.org:10.1002/fsn3.2284

7	Guo, S., Na Jom, K. & Ge, Y. Influence of Roasting Condition on Flavor Profile of Sunflower Seeds: A flavoromics approach. *Sci Rep* **9**, 11295 (2019). https://doi.org:10.1038/s41598-019-47811-3

8	McCluskey, K., Wiest, A. & Plamann, M. The Fungal Genetics Stock Center: a repository for 50 years of fungal genetics research. *J Biosci* **35**, 119-126 (2010). https://doi.org:10.1007/s12038-010-0014-6

9	Galagan, J. E. *et al.* The genome sequence of the filamentous fungus Neurospora crassa. *Nature* **422**, 859-868 (2003). https://doi.org:10.1038/nature01554

10	Ellison, C. E. *et al.* Massive changes in genome architecture accompany the transition to self-fertility in the filamentous fungus Neurospora tetrasperma. *Genetics* **189**, 55-69 (2011). https://doi.org:10.1534/genetics.111.130690

11	Turner, B. C. Two Ecotypes of Neurospora Intermedia. *Mycologia* **79**, 425-432 (1987). https://doi.org:10.1080/00275514.1987.12025400

12	Rahmawati, D., Astawan, M., Putri, S. P. & Fukusaki, E. Gas chromatography-mass spectrometry-based metabolite profiling and sensory profile of Indonesian fermented food (tempe) from various legumes. *J Biosci Bioeng* **132**, 487-495 (2021). https://doi.org:10.1016/j.jbiosc.2021.07.001

13	Sastraatmadja, D. D., Tomita, F. & Kasai, T. Production of High-Quality Oncom, a Traditional Indonesian Fermented Food, by the Inoculation with Selected Mold Strains in the Form of Pure Culture and Solid Inoculum. *J Grad Sch Agr Hokkaido Univ* **70**, **111–127** (2002). https://doi.org:10.11501/3178528

14	Weliana, S., Sari, E. R., & Wahyudi, J. PENGGUNAAN CaCO3 UNTUK MEMPERTAHANKAN KUALITAS TEKSTUR DAN SIFAT ORGANOLEPTIK PISANG AMBON (Musa acuminata) SELAMA PENYIMPANAN. *AGRITEPA* **1** (2015). https://doi.org:https://doi.org/10.37676/agritepa.v1i1.110

15	Kozich, J. J., Westcott, S. L., Baxter, N. T., Highlander, S. K. & Schloss, P. D. Development of a dual-index sequencing strategy and curation pipeline for analyzing amplicon sequence data on the MiSeq Illumina sequencing platform. *Appl Environ Microbiol* **79**, 5112-5120 (2013). https://doi.org:10.1128/AEM.01043-13

16	Callahan, B. J. *et al.* DADA2: High-resolution sample inference from Illumina amplicon data. *Nat Methods* **13**, 581-583 (2016). https://doi.org:10.1038/nmeth.3869

17	McMurdie, P. J. & Holmes, S. phyloseq: an R package for reproducible interactive analysis and graphics of microbiome census data. *PLoS One* **8**, e61217 (2013). https://doi.org:10.1371/journal.pone.0061217

18	Martin, M. Cutadapt removes adapter sequences from high-throughput sequencing reads. *2011* **17**, 3 (2011). https://doi.org:10.14806/ej.17.1.200

19	Bolger, A. M., Lohse, M. & Usadel, B. Trimmomatic: a flexible trimmer for Illumina sequence data. *Bioinformatics* **30**, 2114-2120 (2014). https://doi.org:10.1093/bioinformatics/btu170

20	Clarke, E. L. *et al.* Sunbeam: an extensible pipeline for analyzing metagenomic sequencing experiments. *Microbiome* **7**, 46 (2019). https://doi.org:10.1186/s40168-019-0658-x

21	Menzel, P., Ng, K. L. & Krogh, A. Fast and sensitive taxonomic classification for metagenomics with Kaiju. *Nature Communications* **7**, 11257 (2016). https://doi.org:10.1038/ncomms11257

22	Landis Elizabeth, A. *et al.* Microbial Diversity and Interaction Specificity in Kombucha Tea Fermentations. *mSystems* **7**, e00157-00122 (2022). https://doi.org:10.1128/msystems.00157-22

23	Tovo, A., Menzel, P., Krogh, A., Cosentino Lagomarsino, M. & Suweis, S. Taxonomic classification method for metagenomics based on core protein families with Core-Kaiju. *Nucleic Acids Research* **48**, e93-e93 (2020). https://doi.org:10.1093/nar/gkaa568

24	Vogel, H. J. A convenient growth medium for Neurospora (Medium N). *Microbial Genet. Bull.* **13**, 42-43 (1956).

25	Feng, Y. *et al.* Effect of koji fermentation on generation of volatile compounds in soy sauce production. *International Journal of Food Science & Technology* **48**, 609-619 (2013). https://doi.org:https://doi.org/10.1111/ijfs.12006

26	Jaeger, S. R. *et al.* Check-all-that-apply (CATA) questions for sensory product characterization by consumers: Investigations into the number of terms used in CATA questions. *Food Quality and Preference* **42**, 154-164 (2015). https://doi.org:https://doi.org/10.1016/j.foodqual.2015.02.003

27	Rahmawati, D., Astawan, M., Putri, S. P. & Fukusaki, E. Gas chromatography-mass spectrometry-based metabolite profiling and sensory profile of Indonesian fermented food (tempe) from various legumes. *Journal of Bioscience and Bioengineering* **132**, 487-495 (2021). https://doi.org:https://doi.org/10.1016/j.jbiosc.2021.07.001

28	The Association of Official Analytical Chemists. *Official Methods of Analysis*. 17th edn, (2000).

29	The Association of Official Analytical Chemists. *Official Methods of Analysis*. 18th edn, (2006).

30	Gehrke, C. W., Rexroad, P. R., Schisla, R. M., Absheer, J. S. & Zumwalt, R. W. Quantitative analysis of cystine, methionine, lysine, and nine other amino acids by a single oxidation--4 hour hydrolysis method. *J Assoc Off Anal Chem* **70**, 171-174 (1987).

31	Landry, J. & Delhaye, S. Simplified procedure for the determination of tryptophan of foods and feedstuffs from barytic hydrolysis. *Journal of Agricultural and Food Chemistry* **40**, 776-779 (1992). https://doi.org:10.1021/jf00017a014

32	Rautengarten, C. *et al.* The Arabidopsis Golgi-localized GDP-L-fucose transporter is required for plant development. *Nature Communications* **7**, 12119 (2016). https://doi.org:10.1038/ncomms12119

33	Grigoriev, I. V. *et al.* MycoCosm portal: gearing up for 1000 fungal genomes. *Nucleic Acids Res* **42**, D699-704 (2014). https://doi.org:10.1093/nar/gkt1183

34	Wu, V. W. *et al.* The regulatory and transcriptional landscape associated with carbon utilization in a filamentous fungus. *Proc Natl Acad Sci U S A* **117**, 6003-6013 (2020). https://doi.org:10.1073/pnas.1915611117

35	Kechin, A., Boyarskikh, U., Kel, A. & Filipenko, M. cutPrimers: A New Tool for Accurate Cutting of Primers from Reads of Targeted Next Generation Sequencing. *J Comput Biol* **24**, 1138-1143 (2017). https://doi.org:10.1089/cmb.2017.0096

36	Kim, D., Langmead, B. & Salzberg, S. L. HISAT: a fast spliced aligner with low memory requirements. *Nat Methods* **12**, 357-360 (2015). https://doi.org:10.1038/nmeth.3317

37	Ramírez, F. *et al.* deepTools2: a next generation web server for deep-sequencing data analysis. *Nucleic Acids Res* **44**, W160-165 (2016). https://doi.org:10.1093/nar/gkw257

38	Liao, Y., Smyth, G. K. & Shi, W. featureCounts: an efficient general purpose program for assigning sequence reads to genomic features. *Bioinformatics* **30**, 923-930 (2014). https://doi.org:10.1093/bioinformatics/btt656

39	Robinson, M. D., McCarthy, D. J. & Smyth, G. K. edgeR: a Bioconductor package for differential expression analysis of digital gene expression data. *Bioinformatics* **26**, 139-140 (2010). https://doi.org:10.1093/bioinformatics/btp616

40	Emms, D. M. & Kelly, S. OrthoFinder: phylogenetic orthology inference for comparative genomics. *Genome Biology* **20**, 238 (2019). https://doi.org:10.1186/s13059-019-1832-y

41	Langfelder, P. & Horvath, S. WGCNA: an R package for weighted correlation network analysis. *BMC Bioinformatics* **9**, 559 (2008). https://doi.org:10.1186/1471-2105-9-559

42	Love, M. I., Huber, W. & Anders, S. Moderated estimation of fold change and dispersion for RNA-seq data with DESeq2. *Genome Biol* **15**, 550 (2014). https://doi.org:10.1186/s13059-014-0550-8

43	Prjibelski, A., Antipov, D., Meleshko, D., Lapidus, A. & Korobeynikov, A. Using SPAdes De Novo Assembler. *Curr Protoc Bioinformatics* **70**, e102 (2020). https://doi.org:10.1002/cpbi.102

44	Stanke, M., Diekhans, M., Baertsch, R. & Haussler, D. Using native and syntenically mapped cDNA alignments to improve de novo gene finding. *Bioinformatics* **24**, 637-644 (2008). https://doi.org:10.1093/bioinformatics/btn013

45	Chaudhari, N. M., Gupta, V. K. & Dutta, C. BPGA- an ultra-fast pan-genome analysis pipeline. *Sci Rep* **6**, 24373 (2016). https://doi.org:10.1038/srep24373

46	Götz, S. *et al.* High-throughput functional annotation and data mining with the Blast2GO suite. *Nucleic Acids Res* **36**, 3420-3435 (2008). https://doi.org:10.1093/nar/gkn176

47	Talavera, G. & Castresana, J. Improvement of phylogenies after removing divergent and ambiguously aligned blocks from protein sequence alignments. *Syst Biol* **56**, 564-577 (2007). https://doi.org:10.1080/10635150701472164

48	Minh, B. Q. *et al.* IQ-TREE 2: New Models and Efficient Methods for Phylogenetic Inference in the Genomic Era. *Mol Biol Evol* **37**, 1530-1534 (2020). https://doi.org:10.1093/molbev/msaa015

49	Langmead, B. & Salzberg, S. L. Fast gapped-read alignment with Bowtie 2. *Nat Methods* **9**, 357-359 (2012). https://doi.org:10.1038/nmeth.1923

50	Danecek, P. *et al.* Twelve years of SAMtools and BCFtools. *Gigascience* **10** (2021). https://doi.org:10.1093/gigascience/giab008

51	Blin, K. *et al.* antiSMASH 7.0: new and improved predictions for detection, regulation, chemical structures and visualisation. *Nucleic Acids Research* **51**, W46-W50 (2023). https://doi.org:10.1093/nar/gkad344

52	Wang, M. *et al.* Sharing and community curation of mass spectrometry data with Global Natural Products Social Molecular Networking. *Nat Biotechnol* **34**, 828-837 (2016). https://doi.org:10.1038/nbt.3597

53	Dührkop, K. *et al.* SIRIUS 4: a rapid tool for turning tandem mass spectra into metabolite structure information. *Nat Methods* **16**, 299-302 (2019). https://doi.org:10.1038/s41592-019-0344-8
